# Globally prevalent Kelch13 mutations increase partial artemisinin resistance and fitness in Bangladeshi *Plasmodium falciparum* parasites

**DOI:** 10.1101/2025.04.25.650672

**Authors:** Maisha Khair Nima, Nirjhar Bhattacharyya, Jungyoon Park, Douglas Shoue, Ching Swe Phru, Saiful Arefeen Sazed, Sudhir Kumar, Mohammad Shafiul Alam, Michael Ferdig, Angana Mukherjee

## Abstract

Artemisinin partial resistance (ArtR), mediated by Kelch13 (K13) gene mutations in *Plasmodium falciparum*, has caused delayed parasite clearance and, together with partner drug resistance, treatment failures in the Greater Mekong Subregion (GMS). The Chittagong Hill Tracts (CHTs), located in southeastern Bangladesh and bordering Myanmar and India, regions with prevalent K13 mutations, account for approximately 90% of the country’s malaria infections but have not yet reported ArtR-causing K13 mutations. Importantly, however, some isolates from the CHTs have demonstrated moderate *in vitro* ArtR in the absence of K13 mutations. To assess the potential threat of K13-mediated ArtR in Bangladesh, we proactively evaluated the impact of three prevalent neighboring K13 mutations (F446I, R561H, and C580Y) on ArtR and parasite fitness in parasites isolated from the CHTs. We edited these mutations into two distinct genetic backgrounds: an artemisinin-sensitive strain (CHT-S) and a K13-independent, moderately resistant strain (CHT-R). In these edited lines, we then evaluated ArtR levels and fitness using ring-stage survival assays (RSA), post-drug treatment recovery assays, and competitive fitness assays against isogenic control strains. Prior to genome editing, both isolates were characterized for baseline drug susceptibility, resistance-associated mutations, and population structure. We found that C580Y and R561H mutations, but not F446I, confer increased *in vitro* ArtR in the CHT-R background already exhibiting K13-independent moderate resistance. All three mutations incurred minimal or no fitness costs in both genetic backgrounds. Notably, R561H, the dominant allele at the Thai-Myanmar border and currently expanding in Rwanda, mediates extreme resistance in the CHT-R background, with mean RSA survival rates of 30.9±1.9%, an unprecedented resistance level among K13-engineered lines. R561H also showed fitness advantages (0.5% per generation) and the highest growth recovery post-treatment. This represents the first experimental study modeling this K13 mediated ArtR risk in Bangladeshi parasites. In conclusion, we found that indigenous *P. falciparum* isolates in the CHTs possess inherent genetic potential to sustain high-level ArtR and fitness advantages if K13 mutations emerge, raising urgent concerns for containment and surveillance strategies during Bangladesh’s malaria elimination phase.

## Introduction

In 2015, World Health Organization member states adopted a strategy to reduce malaria incidence and mortality by at least 90% by 2030 (*Global Technical Strategy for Malaria 2016-2030, 2021 Update* 2021). However, more than halfway to the target date, malaria remains a major global health threat, with 263 million cases and 597,000 deaths reported in 2023 (“World Malaria Report 2024,” n.d.). In the absence of an effective malaria vaccine, control and elimination efforts heavily rely on the clinical efficacy of artemisinin-based combination therapies (ACTs) which combine fast-acting artemisinin derivatives with partner drugs such as lumefantrine, amodiaquine, mefloquine, pyronaridine, sulfadoxine–pyrimethamine or piperaquine(“WHO Guidelines for Malaria” 2017).

In Bangladesh, where approximately 18 million people are at risk of malaria, substantial progress has been made in malaria control. From 2010 to 2020, the country achieved 93% reduction in malaria cases and is actively working towards a complete national elimination by 2030(“National Strategic Plan_Malaria Elimination_Bangladesh_2021-2025.pdf,” n.d.) by interrupting local transmission, preventing re-establishment, and impeding emergence of ACT resistance. However, despite these efforts, *Plasmodium falciparum*, the primary cause of severe malaria and responsible for 80-90% of malaria infections in Bangladesh(Noé et al. 2018; Haque et al. 2014; Maude et al. 2012; Shannon et al. 2016), remains prevalent. Transmission is concentrated in the forested, mountainous Chittagong Hill Tracts (CHTs), comprising the districts of Bandarban, Rangamati, and Khagrachhari, which account for 80-90% of the country’s malaria cases(Huwe et al. 2022; Noé et al. 2018; Sinha et al. 2020)

The spread of antimalarial drug resistance is a critical barrier to malaria elimination globally(White 2004). In the late 2000s, partial artemisinin resistance (ArtR) emerged in Cambodia, characterized by prolonged parasite clearance times in patients, following treatment with artemisinin derivatives or ACTs(Noedl et al. 2008; Dondorp et al. 2009; Denis et al. 2006; Alker et al. 2007). Genome-wide association studies subsequently linked delayed clearance to mutations in the *pfkelch13* (K13) propeller domain(Cheeseman et al. 2012; Takala-Harrison et al. 2013; Anderson et al. 2010). Although ArtR can have a multigenic basis(Miotto et al. 2015; Demas et al. 2018; Mbengue et al. 2015; Rocamora et al. 2018; Mukherjee et al. 2017; Birnbaum et al. 2020; Sutherland, Henrici, and Artavanis-Tsakonas 2021), K13 mutations are the major genetic determinant of resistance, originally identified through *in vitro* evolution and genome sequencing, confirmed through field studies(Ariey et al. 2014) and subsequently verified through reverse genetics(Straimer et al. 2015). The standard *in vitro* correlate for ArtR, the Ring-stage Survival Assay (RSA) measures early ring-stage-specific resistance by exposing parasites to pharmacologically relevant dosage of active metabolite; dihydroartemisinin (DHA), during the first six hours of the asexual cycle(Witkowski, Amaratunga, et al. 2013). To date, over 200 propeller domain K13 substitutions have been found globally, of which only 14 have been validated as contributing to ArtR as measured by *in vitro* and *ex vivo* RSA per WHO guidelines(Uwimana et al. 2020; Straimer et al. 2015; Stokes et al. 2021; Siddiqui et al. 2020; World Health Organization 2023). These mutations include P441L, F446I, C469Y, N458Y, M476I, Y493H, R539T, I543T, P553L, R561H, P574L, C580Y, R622I, and A675V.

In the Greater Mekong Subregion (GMS), the emergence and spread of ArtR-associated K13 mutations, coupled with resistance to partner drugs, have led to ACT failure rates upto 50%(van der Pluijm et al. 2019). These mutations have disseminated across eastern GMS countries countries via clonal expansion (Cambodia, Vietnam, Laos and Thailand) (Miotto et al. 2015; Ménard et al. 2016; Amato et al. 2018; Imwong et al. 2017), and have independently emerged in Myanmar, China-Myanmar border and Thai-Myanmar border(Ye et al. 2016; Takala-Harrison et al. 2015; Imwong et al. 2017). K13 mutations have also been reported in northeastern India bordering the CHTs (Chhibber-Goel and Sharma 2019; Mishra et al. 2015). This pattern resembles historical resistance trends, notably chloroquine and sulfadoxine-pyrimethamine resistance, that spread from Southeast Asia through Bangladesh and India to Africa, causing significant morbidity and mortality(Mita, Tanabe, and Kita 2009). More recently, independent K13 mutations with clinical and *in vitro* evidence of ArtR have also emerged in South America, East Africa, and the Horn of Africa, further broadening global concern (Uwimana et al. 2020, 2021; Bergmann et al. 2021; Straimer et al. 2022; Juliano et al. 2023; Conrad et al. 2023; Balikagala et al. 2021; Tumwebaze et al. 2022; Pillai et al. 2016; Mihreteab et al. 2023; Fola et al. 2023; “More on Artemisinin-Resistant *Plasmodium Falciparum* in Eastern India” 2019), (Mathieu et al. 2020) .

Despite no current detection of K13-mediated ArtR(Nima et al. 2022; Alam et al. 2017; Mohon et al. 2014), the CHTs in southeastern Bangladesh remain highly vulnerable due to their geographic proximity to K13 associated ArtR-prevalent regions in Myanmar and India, and shared climatic conditions, seasonal transmission patterns, and malaria control challenges(Lalmalsawma et al. 2023). Other malaria-endemic countries with similar timeline of of ACT usage(Takala-Harrison et al. 2015; Mathieu et al. 2020; Miotto et al. 2020; Lu et al. 2017; Chenet et al. 2016; Imwong et al. 2017; Kyaw M. Tun et al. 2015) and transmission settings(Huwe et al. 2022; Fola et al. 2017; Nkhoma et al. 2013; Kateera et al. 2016) have also witnessed expansion and spread of K13 associated ArtR. This collective evidence underscores the high likelihood that the CHTs could experience the emergence and expansion of ArtR through both regional spread and independent evolution, posing significant threats to Bangladesh’s malaria elimination efforts.

In 2018 and 2019, we assessed ACT susceptibility in malaria patients with *P. falciparum* mono infections from Bandarban and Alikadam districts of the CHTs using clinical parasite clearance assays. The median time to clear half of the initial parasite load was 5.6 hours (range: 1.5 to 9.6 hours), with 20% of patients exhibiting a median of 8 hours (Nima et al. 2022). Additionally, we also detected quantifiable *in vitro* resistance by RSA that was independent of K13 mutations.

Given these findings, the main objective of the current study was to proactively assess the phenotypic and fitness consequences of three highly prevalent K13 mutations (F446I, R561H, and C580Y) using genome editing in two distinct CHT patient isolates collected in 2018 and 2019 in our previous study(Nima et al. 2022). These K13 mutations were chosen for their highest prevalence, their historical emergence in both proximal and distal locations and their temporal expansion. We introduced each mutation into two genetically distinct CHT isolates: an artemisinin-sensitive strain (CHT-S) and a K13-independent moderately resistant strain (CHT-R). We systematically assessed the impact of these K13 mutations and the influence of the genetic backgrounds through quantitative RSAs, post-treatment recovery assays, and competitive fitness assays.

## Results

### Spatio-temporal analysis of prevalence of WHO validated K13 mutations in malaria endemic regions

Fourteen non-synonymous K13 mutations, validated and documented by WHO as markers of ArtR, exhibit varying prevalence across malaria-endemic regions worldwide. We conducted a comprehensive analysis of their frequency using data from the Worldwide Antimalarial Resistance Network(“Artemisinin Molecular Surveyor,” n.d.), supplemented by malaria threat maps and recent studies(Kagoro et al. 2022; Mihreteab et al. 2023; Ndwiga et al. 2021; Uwimana et al. 2021; Asua et al. 2021; Conrad et al. 2023; Fola et al. 2023; He et al. 2019) (Source Table S1). We analyzed 72,375 samples collected from regions where WHO-validated K13 mutations have independently emerged or spread, spanning data from 1997 to the most recent sampling year for each country, covering seven countries in southeast Asia and South Asia, fourteen countries in Africa, Papua New Guinea in Oceania, and Guyana in South America. The K13 3D7 reference sequence was present in the majority of samples (88.7%, n=72,375) (Supplementary Table S1). Cumulative prevalence of each of the K13 propeller domain mutations spanning 1997-2022 (Supplementary Table S1) demonstrate that C580Y (49.88%), F446I (14.3%), and R561H (7.65%), followed by P441L (6.58%), R539T (4.28%), and Y493H (3%) were the most frequent mutations. Next, we narrowed our focus to a detailed temporal and spatial analysis of these mutations over the last 15 years (2007–2022), a period chosen because this was the inflection point where the total number of kelch13 mutations started increasing. Supplementary Figure 1 depicts the global temporal progression of these validated K13 mutations worldwide. The spatio-temporal analysis, presented in Figure 1, highlights the dynamics of K13-mediated ArtR emergence and spread during this period. Our analysis indicates that C580Y, the most prevalent mutation worldwide, emerged in 9 countries, the highest number of countries any validated mutations have emerged, initially emerging in Thailand and Cambodia in 2007-08 reaching 80-100% prevalence in 2015-16. Other GMS countries such as Laos and Vietnam also saw emergence and expansion upto 70 and 80% prevalence in 2014 and 2019 respectively. Outside of the GMS, Angola, Papua New Guinea and Guyana have also reported emergence from 2-5% over the last decade. F446I, the second most prevalent mutation worldwide, also shows very high prevalence in the Myanmar and China-Myanmar border, increasing from a frequency of approx. 20-55% in years 2007 to 2015-16. (Figure 1). The third highest prevalent mutation R561H emerged in Thailand in 2009, reaching 30% frequency in 2014. In 2018, Rwanda saw an emergence at >10% followed by Tanzania in 2021-2022 (Figure 1). Other mutations have shown regionally restricted or declining trends. R539T peaked in Cambodia in the early 2010s but has since declined, while P441L and Y493H resurged in Myanmar and Cambodia, respectively, around 2019 (Figure 1). Sporadic GMS mutations such as P574L, N458Y, I543T, and M465I remain limited in prevalence. R622I has recently emerged in the horn of Africa in Eritrea and Ethiopia, associated with delayed clearance. Mutations such as C469Y and A675V, first observed in Uganda in 2016–2017, are rising in multiple districts in Uganda. Although these mutations provide valuable insights into localized resistance dynamics, their sporadic and regionally restricted nature contrasts with the widespread emergence and fixation of mutations like C580Y, F446I, and R561H.

**Figure 1.**
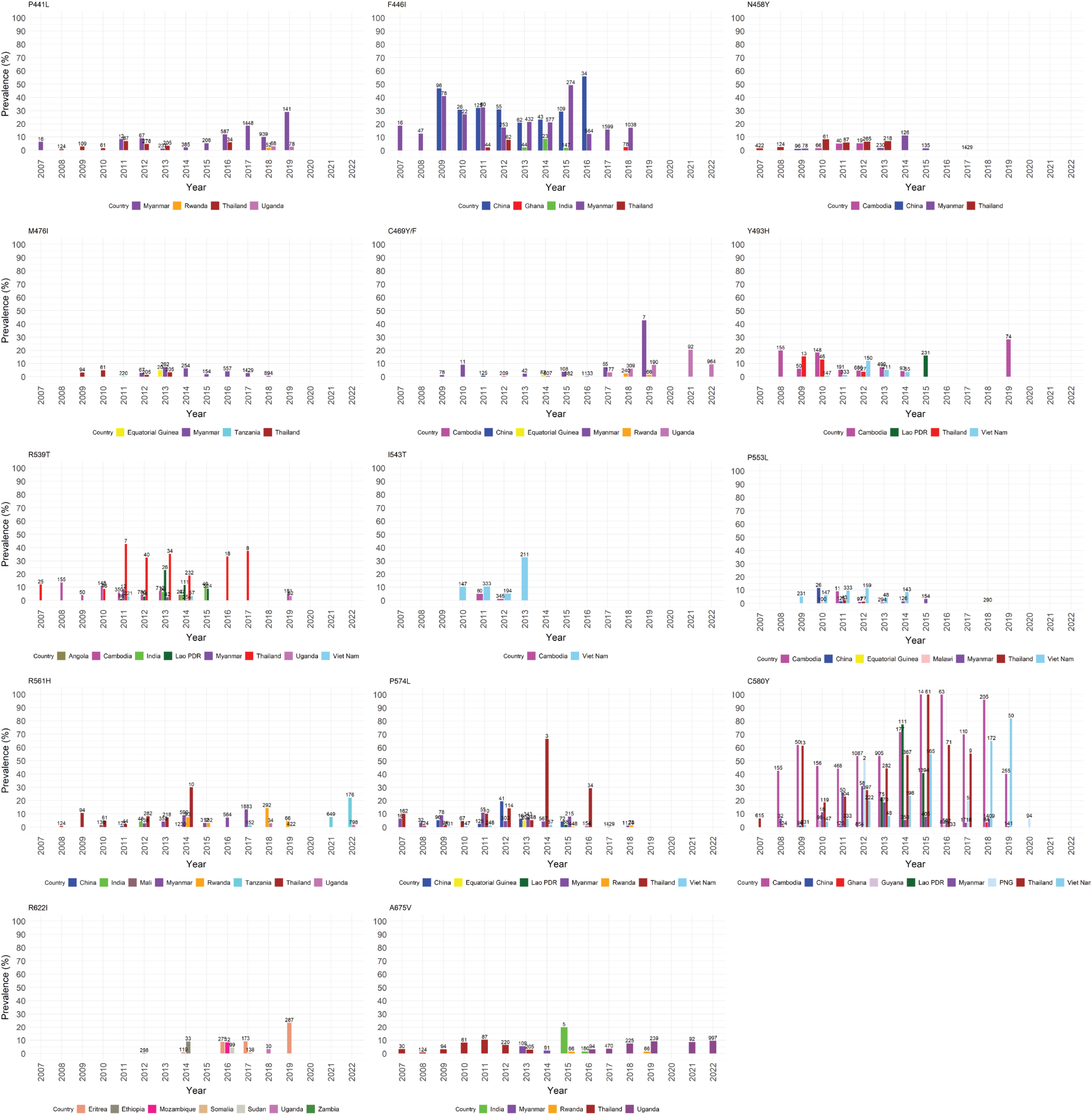
Spatio-temporal analysis highlighting the prevalence dynamics of WHO validated K13 mutations globally from 2007 to 2022.

To provide a global snapshot of these three most prevalent K13 mutations—C580Y, R561H, and F446I, we analyzed their frequency across the GMS, and regions of independent emergence, including East Africa, Guyana, and Papua New Guinea. Figure 2A presents the most recent prevalence data for each region, while Figure 2B illustrates temporal trends.

**Figure 2.**
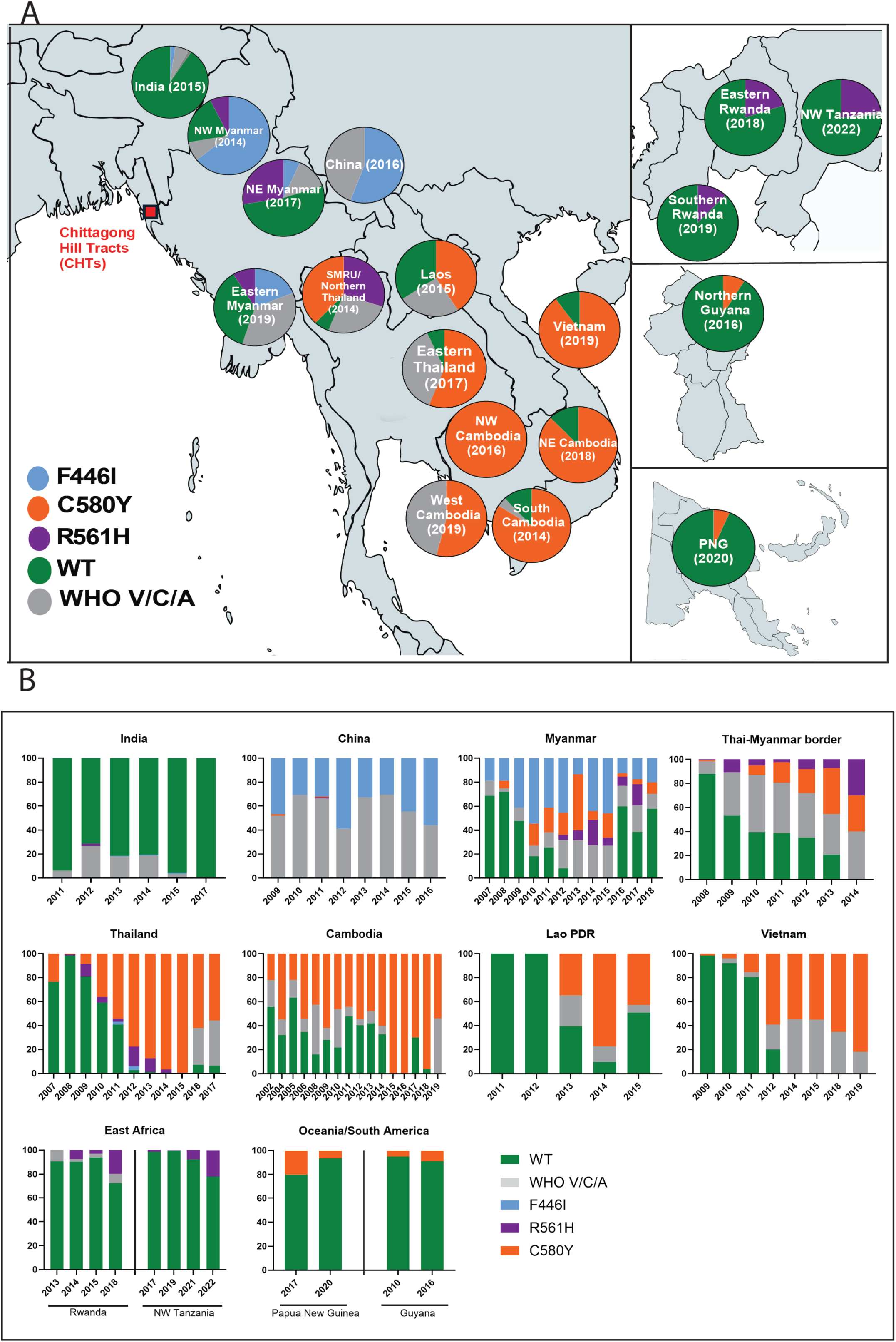
Frequency of commonly emerged and expanded pfK13 mutations observed globally. A. Colored pie charts in sectioned maps of the globe showing prevalence of K13F446I, K13R561H, K13C580Y in the GMS, K13R561H in Rwanda and Tanzania, K13C580Y in Guyana and Papua New Guinea , sampling year in parenthesis for each region. B. Temporal prevalence of K13F446I, K13R561H, K13C580Y in key regions of emergence. See Source Files Figure 2A and 2B for the raw data.

C580Y has been the predominant K13 mutation in the GMS for the past decade. Its prevalence in Cambodia ranges from 40–100%, reaching full 100% fixation in North-West Cambodia by 2016, and has reached 87% in North-East and 54% in West Cambodia in 2019 (Figure 2A). In Thailand, Laos, and Vietnam, C580Y remained prevalent at 40–80% (2015–2017) (Figure 2A). Figure 2B shows that C580Y outcompeted other mutations and WT alleles by 2016-2018 in Thailand and Cambodia. Independent emergences in Papua New Guinea and Guyana around 2010 have maintained low but stable prevalence over the past decade (Figure 2).

F446I is dominant in Myanmar, particularly near the China-Myanmar border, peaking at 50% in 2015, X in North-West Myanmar 20% in Eastern Myanmar also present in Northeast India near the Bangladesh border (Figure 2A). Temporally, unlike C580Y it did not outcompete other mutations and WT alleles and was present in the Myanmar population at 20% as early as 2007, increasing to 60% by 2010 (Figure 2B). Its dominance along the Myanmar-China border and spread to Northeast India highlight its regional impact.

R561H remains prevalent at 33% near the Thai-Myanmar border, it’s also high in Myanmar, ranking as the second most common mutation in Myanmar (Figure. 2A and Source File-Figure 2A). In eastern Thailand, the small proportion of R561H mutation was swept away by C580Y between 2009-2015. Outside the GMS, R561H has independently emerged as the most prevalent mutation in Rwanda, with a significant frequency increase observed in 2018. In North-West Tanzania, which borders Rwanda, R561H has also emerged and expanded since 2018 (Figure 2B).

### Genomic and *in vitro* Phenotypic Characterization of an artemisinin sensitive (CHT-S) and a K13 independent moderate artemisinin resistant (CHT-R)

The CHTs border northeastern India and Myanmar where K13 mediated ArtR is prevalent(Kyaw M. Tun et al. 2015; Nyunt et al. 2015; Chhibber-Goel and Sharma 2019). Approximately 90% of Bangladesh’s *P. falciparum* malaria cases occur within the forested, hilly Chittagong Hill Tracts (CHTs), comprising three districts, including our specific study site in Bandarban. Although ArtR-associated K13 mutations have not yet been reported from the CHTs or elsewhere in Bangladesh, we analyzed 1,310 samples from the broader Chittagong division (which includes the three CHT districts and eight additional districts), collected between 2008 and 2018, from the MalariaGEN Pf7 dataset(MalariaGEN et al. 2023). We identified five samples harboring the K13 mutation A578S, and seven additional K13 mutations—T573I, S711T, P475S, N523D, G674R, E605K, and A724V—each detected in only one sample. None of these mutations are associated with or validated for ArtR, and their lack of expansion suggests they are not driven by ART selection pressure in the population. Although K13 resistance-associated mutations remain unreported in Bangladesh(Nima et al. 2022; Mohon et al. 2014; Alam et al. 2017), our previous study in Bandarban (2018–2019) identified K13-independent ArtR in one *P. falciparum* isolate(Nima et al. 2022). We short-term lab-adapted a cohort of these patient isolates and selected two for genetic and phenotypic profiling: an artemisinin-sensitive isolate (I-001, CHT-S) with a ring-stage survival rate of 0.6% ± 0.13% (Nima et al. 2022), Table 1) and an artemisinin-resistant isolate (I-029, CHT-R) with a survival rate of 6.4% ± 0.53% (Table 1). Microsatellite analysis revealed that I-029 was polygenomic, clonal populations were derived from I-029 through limited dilution cloning and confirmed using microsatellite genotyping(Tirrell et al. 2019). We performed whole-genome sequencing on these two isolate clones. In both CHT-S and CHT-R, the full K13 gene remained wild type (3D7), as verified by whole-genome and Sanger sequencing (Table 1). In 72-hour *in vitro* drug testing, both isolates demonstrated sensitivity to artemisinin, resistance to chloroquine and cycloguanil (active form of the antifolate proguanil), carrying key substitutions such as K76T in *pfCRT* and the *pfDHFR* quadruple mutation haplotype (IRNL) (Table 1). While CHT-S maintains a wild-type *Pfmdr1* genotype (3D7-like), CHT-R possesses the N86Y allele mutation. Additional genotypes in other candidate genes for ArtR are summarized in Supplementary Table S2.

**Table 1.**

| Strains | In vivo characteristics |  |  | In vitro drug phenotypes |  |  |  |  |  |  |  | Drug resistant marker genotypes |  |  |  |  |
| --- | --- | --- | --- | --- | --- | --- | --- | --- | --- | --- | --- | --- | --- | --- | --- | --- |
|  |  |  |  | RSA (% survival) | IC50 (nM) |  |  |  |  |  |  |  |  |  |  |  |
|  | Collection Year | Parance half-life | PC50 (h) | DHA | ART | CQ | PPQ | Cycloguanil | LUM | MFQ | AMDQ | K13<br>(PF3D7_1343700) | CRT<br>(PF3D7_0709000) | DHFR-TS<br>(PF3D7_0417200) | MDR1<br>(PF3D7_0523000) | PPPK-DHPS<br>(PF3D7_0810800) |
| CHT-S (I-001) | 2018 | 3.3 | 0.015 | 0.66±0.13 | 2.5±2 | 544±54.29 | 4.454±0.7427 | 7832±881.2 | 0.88±0.32 | 9.66±4.03 | 5.63±0.23 | WT | K76T, A220S, Q271E | N51I, C59R, S108N, I164L | N86 | A581G |
| CHT-R (I-029) | 2019 | 2.39 | 7.68 | 6.4±0.53 | 2.5±1.6 | 386.5±27.23 | 3.486±1.26 | 362.6±49.27 | 0.39±0.14 | 3.53±0.8 | 2.39±0.67 | WT | K76T, Q271E, R371I | N51I, C59R, S108N | N86Y | S436A, K540E |

To investigate the population structure and genetic relationships among CHT-S, CHT-R and neighboring regions, we conducted Principal Component Analysis (PCA) (Figure 3), comparing CHT-S and CHT-R with publicly available genomic sequences from MalariaGen Pf7 dataset, from the Chittagong division in Bangladesh, as well as neighboring Myanmar, India, Thailand, Cambodia, Vietnam and Indonesia, with representative samples from each country, collected from 2008-2018, including a total of 696 samples. Our CHT-S and CHT-R cluster closely with 93 other Bangladeshi and 198 Myanmar isolates. The samples from Myanmar, display a broader distribution in the PCA space, indicating a genetically more diverse or distinct population structure relative to Bangladesh. The additional pairwise PCA plots (PC1 vs. PC3 and PC2 vs. PC3) in Supplementary Figure 2A further illustrate these patterns, showing consistent clustering of the CHT samples with other Bangladeshi/Myanmar isolates across three principal component axes, which supports the robustness of this genetic grouping. Supplementary Figure 2B shows the variance explained by the first 10 principal components, out of which the first three explains the majority of the variance.

**Figure 3:**
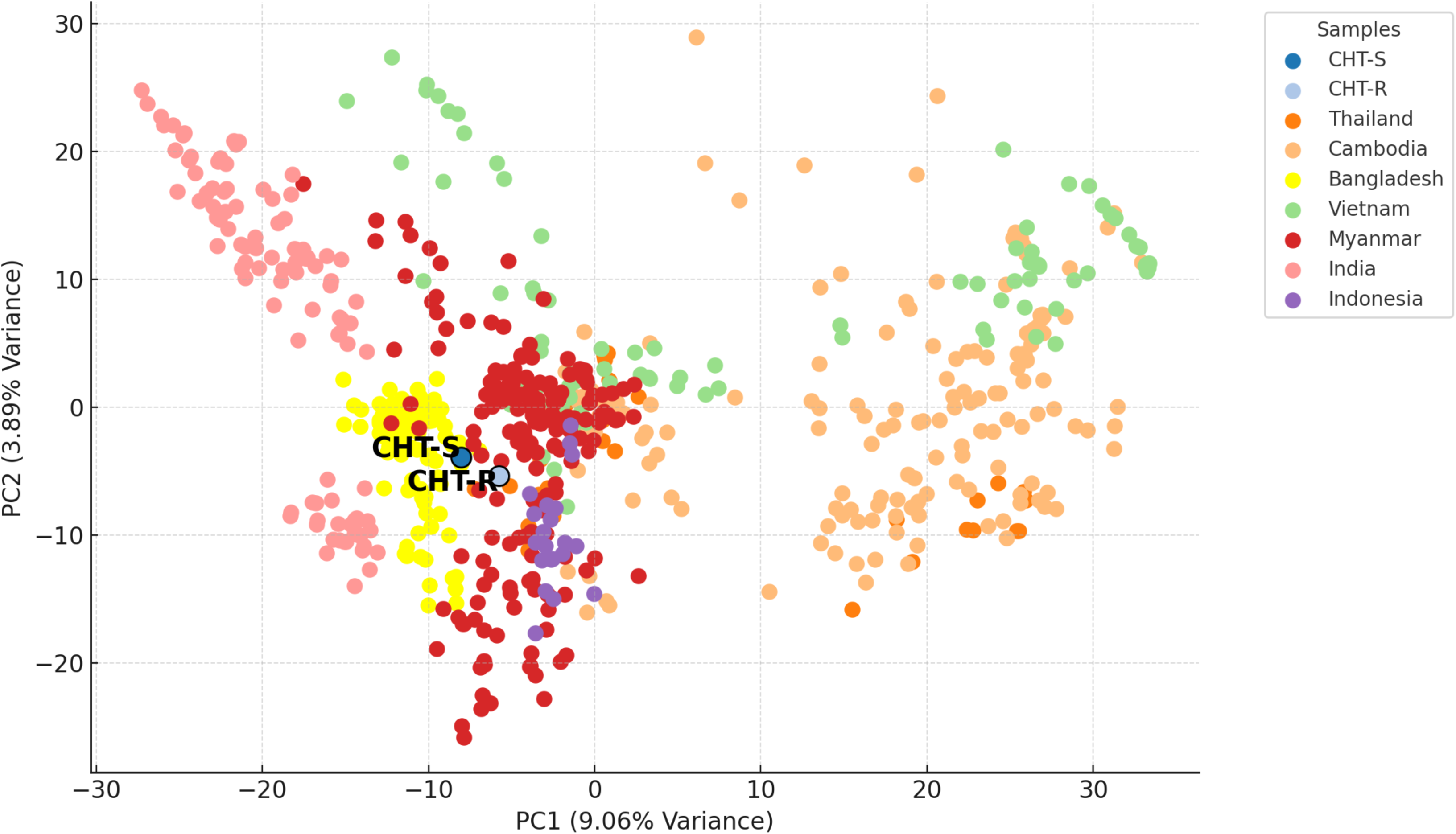
Principal Component Analysis (PCA) of our whole genome sequenced CHT samples merged with other available *P. falciparum* genomes from Bangladesh and neighboring countries. PCA of the CHT-S and CHT-R and publicly available genomic sequences from the MalariaGen Pf7 dataset, from Chittagong (a division covering the CHTs and the coastal plains) in Bangladesh and neighboring Myanmar, India, Thailand, Cambodia, Vietnam and Indonesia, with representative samples from each country, collected from 2008-2018, including a total of 696 samples. The first two Principal Components (PC1 and PC2) explains 9.06% and 3.89% variance of the multisample genomic data.

### C580Y, and R561H drive higher resistance in the CHT-S and CHT-R strains while F4461 shows a minimal effect

To evaluate the effect of the ArtR-conferring three dominant worldwide nonsynonymous K13 mutations (F446I, R561H, and C580Y) in the CHTs, we used CRISPR Cas9 to edit these mutations in the CHT-S and CHT-R isolates. As an editing control we knocked in two K13 synonymous shield mutations that do not alter amino acids. Successful knockins of K13 mutations were verified by Sanger sequencing. An independent clone for each background was obtained by limiting dilution of the bulk transfectant cultures, K13 domains sequenced (Supplementary Figure 3) and used in further analyses. To test for ArtR, we used RSA where 0 to 3 hour ring-stage parasites are subjected to 700nM DHA treatment for 6h and survival rates are calculated 72h later. A threshold of >1% survival is considered resistant(Witkowski, Amaratunga, et al. 2013). To control for our RSA survival data in the new edited lines, we also performed assays using previously characterized K13-edited laboratory strain Dd2 lines (R561H, and C580Y), whose RSA phenotypes have been published(Stokes et al. 2021).Our results (Supplementary Figure 4)match with the published RSA data from the Dd2 K13 edited lines. Our RSA results (Figure 4A and Source File-Figure 4A) compare the survival rates of K13-edited CHT strains. Control lines, CHT-S^ctrl^ and CHT-R^ctrl^, which contain only synonymous mutations, did not show a significant increase in resistance compared to their respective parent lines (unpaired t-test, ns, not shown) and were used as comparators in all statistical tests. Figure 4B and Source File-Figure 4B provide a meta-analysis of RSA survival rates of these K13 edits across different genetic backgrounds.

**Figure 4.**
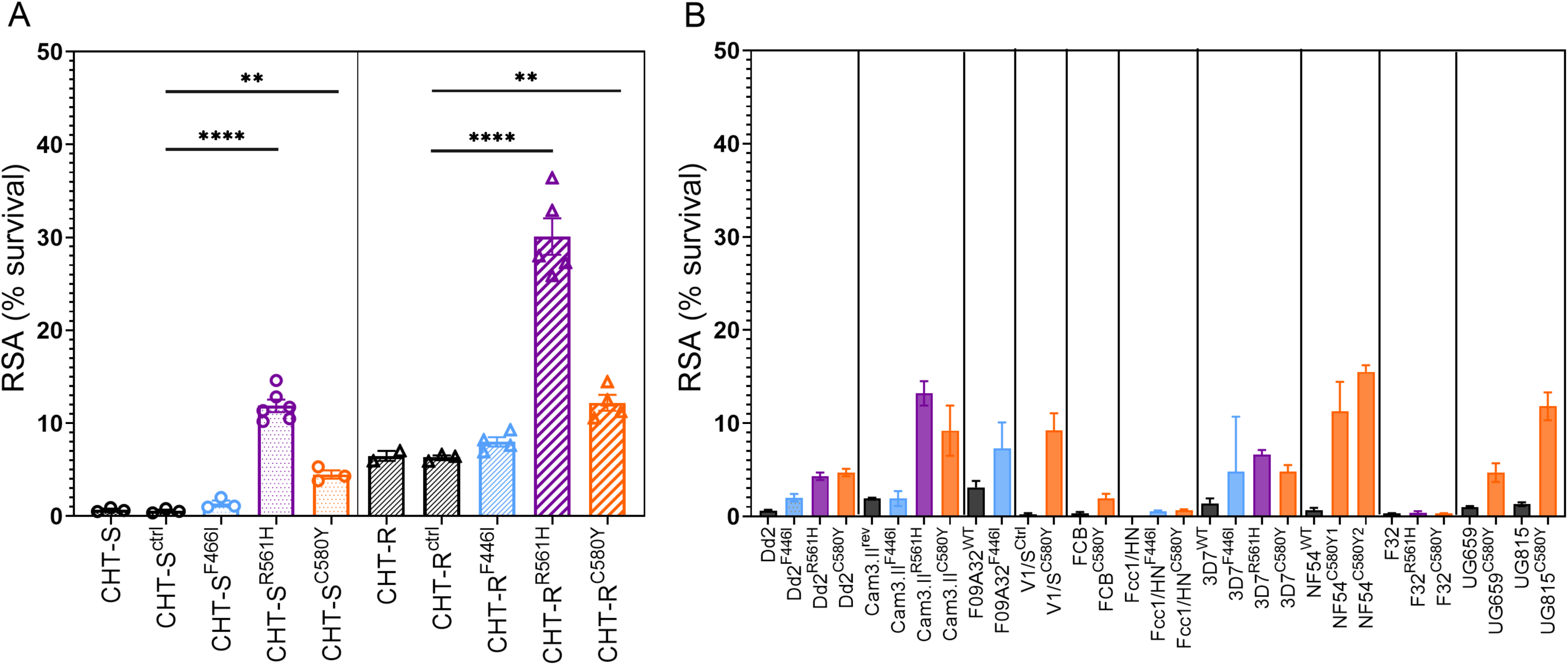
In vitro ART susceptibility (by RSAs) in genome edited K13 F446I, R561H and C580Y mutations in various genetic backgrounds A. RSA survival of F446I, R561H, C580Y and control silent edits in the CHT-S (left panel) and in the CHT-R (right panel) backgrounds. Statistical evaluations were performed with unpaired t-tests (with Welch’s correction) using CHT-S^ctrl^ and CHT-R^ctrl^ (K13 shield mutations/binding site control) as the comparators for all edits in the respective backgrounds. B. Meta-analysis of the RSAs from the same K13 genetic edits across different backgrounds, including laboratory strains and field-adapted strains from Cambodia, Tanzania, Uganda, China-Myanmar border and reference strains from Vietnam, Hainan province of China, Indo-China, West Africa(Stokes et al. 2021; Siddiqui et al. 2020; Uwimana et al. 2020; Ghorbal et al. 2014; Straimer et al. 2015; O’Neill et al. 2017). See Source Files-Figure 4A and Figure 4B for raw RSA data.

C580Y, the most prevalent and independently emerged mutation, produced RSA survival rates of 4.2% ± 0.6 [mean ± S.E.M] in CHT-S, representing an 8-fold increase in mean resistance compared to control line CHT-S^ctrl^ (unpaired t-test, p<0.01, Figure 4A) and comparable to the resistance potential of C580Y in Dd2 (Supplementary Figure 4), as well as 3D7 and Ugandan backgrounds (UG659C580Y and Dd2) in other studies (Figure 4B, orange bars, Source File-Figure 4B). In CHT-R, C580Y showed RSA levels of 12.22% ± 0.7 [mean ± SEM], indicating a 2- fold increase in resistance compared to CHT-R^ctrl^ (unpaired t-test, p<0.01), similar to resistance levels observed in other backgrounds such as NF54 and Cam.3.II (Figure 4B).

Notably, R561H, exhibited significantly high survival rates in both CHT-S and CHT-R backgrounds, with a 20-fold increase in resistance (12.33% ± 0.93, unpaired t-test, p<0.0001) in CHT-S and a 4.8-fold increase (30.09% ± 1.97, unpaired t-test, p<0.001) in CHT-R compared to their respective parent control lines. This unprecedented level of resistance has not been observed when introducing these mutations into any genetically engineered K13 line to date (Figure 4B).

F446I, had a minimal impact on elevating survival in both CHT-S and CHT-R backgrounds. In CHT-S, F446I demonstrated RSA of (1.5% ± 0.32, unpaired t-test, ns) similar to the resistance level observed in F446I engineered Dd2 and Cam3.II background, (Figure 4B, blue bars). In CHT-R, F446I had a modest and non-significant effect on elevating survival, with a rate (10.11% ± 1.67, unpaired t-test, ns) compared to its isogenic edited control.

The CHT-R background sustained the highest resistance upon acquiring R561H mutation compared to the sensitive background and previously studied K13-engineered lines (Figure 4B) (unpaired t-test, p < 0.0001, Source File-Figure 4B).

### K13 mutations confer minimal fitness costs in Bangladeshi *P. falciparum* isolates

To assess the asexual fitness impact of K13 substitutions, we performed replicated head-to-head growth competition assays. Tightly synchronized isogenic clones, differing by one only K13 allele (K13-edited control and the other a missense K13-edited variant)—were co-cultured at a 1:1 ratio and maintained for 40 days. The relative growth of each clone was determined by quantifying the proportion of the corresponding K13 allele every 48 hours using qPCR with a RhAmp SNP kit (IDT) (see Methods).

The competitive growth assay results (Figure 5A, B, Source File-Figure 5A) demonstrate that the acquisition of F446I, R561H, and C580Y mutations exerts minimal to no fitness costs in both CHT-S and CHT-R backgrounds. In the CHT-S background, F446I provides a fitness advantage, reaching 100% allele percentage by Day 20 (∼10 cycles). The proportion of R561H and C580Y remained relatively stable over time, with only slight fluctuations observed (Figure 5A). In contrast, in the CHT-R background, the proportion of F446I declined 28% by day 40 (20 cycles). C580Y showed minimal fluctuations over the 40 days of competition, while R561H provided a fitness advantage, reaching 95% allele percentage by Day 40.

**Figure 5.**
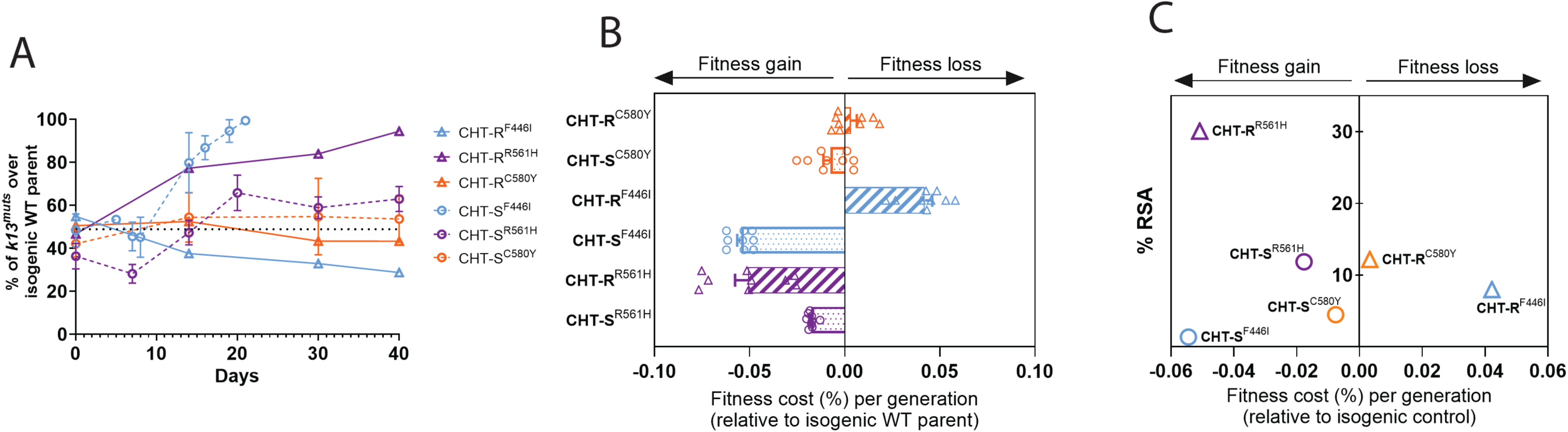
Head-to-head competitive fitness assays in the edited lines show no or minimal fitness costs. **A**. Percentages of mutant K13 alleles relative to the silent edited control in the respective background over time. CHT-S^ctrl^ and CHT-R^ctrl^ were mixed at 1:1 proportion with each of its edited lines (F446I/C580Y/R561H) on day 0 and cultured in 96 well plates for 40 days. Results shown as means ± SEM, were obtained from three biological replicates performed in technical triplicates. Values are provided in Source File-Figure 5A. **B.** Percent difference in growth rate per 48 hr generation, expressed as fitness costs, for CHT-S and CHT-R mutant lines relative to their isogenic silent-edited control lines, expressed as mean ± SEM. **C.** Fitness costs of the mutant lines plotted against their corresponding RSA survival values.

From these data, we calculated the percent increase in growth rate per 48-hour generation, expressed as fitness costs, for each K13 mutant line relative to its isogenic control (Figure 5B), where a negative fitness cost value signifies a fitness advantage. The fitness costs of the CHT-S mutant lines ranged from -0.05% to 0% per generation, while the CHT-R mutant lines exhibited fitness costs of -0.05% to 0.04% per generation. F446I in CHT-S demonstrated a fitness advantage (0.5% per generation) and showed minimal fitness cost in CHT-R (0.04% per generation), C580Y demonstrated fitness neutrality in both CHT-S and CHT-R with 0% fitness cost per generation. R561H demonstrated a fitness advantage in CHT-R (0.5% per generation) but was fitness neutral in CHT-S (0.1% growth per generation).

Plotting the fitness costs against the corresponding RSA survival values for each mutant line revealed no clear correlation between these two parameters in the CHT isolates (spearman r= 0.02, ns) (Figure 5C). Notably, this analysis reveals that R561H is both fit and exhibits high resistance (Figure 5C).

### K13 R561H edited lines recover fastest post DHA treatment

Following a 6-hour treatment of 0–3 h ring-stage parasites with 700 nM DHA, growth recovery was monitored by measuring parasitemia over 8 days (192 hours) (Figure 6). Linear regression analysis revealed distinct recovery patterns among edited lines. For CHT-S isolates, the R561H mutation displayed the steepest recovery trajectory with a slope of 0.5 and an R² value of 0.8, displaying significant post DHA treatment growth. This was notably higher compared to the other mutations: F446I and C580Y showed minimal growth recovery with slopes of 0.09 and 0.04, respectively, and an R² of 0.9, comparable to the transfected control line (slope=0.04, R²=0.9) suggesting CHT-S with K13 F446I and C580Y confer negligible to minimal growth advantage in the presence of DHA. In the CHT-R background, the R561H mutation again demonstrated the highest recovery rate (slope = 0.5, R² = 0.89), with no change between the CHT-S and CHT-R backgrounds. The C580Y mutation displayed a moderate recovery slope of 0.26 (R² = 0.85), while F446I had a similar recovery slope of 0.17 (R² = 0.84), compared to the control (slope=0.15, R² = 0.94).

**Figure 6.**
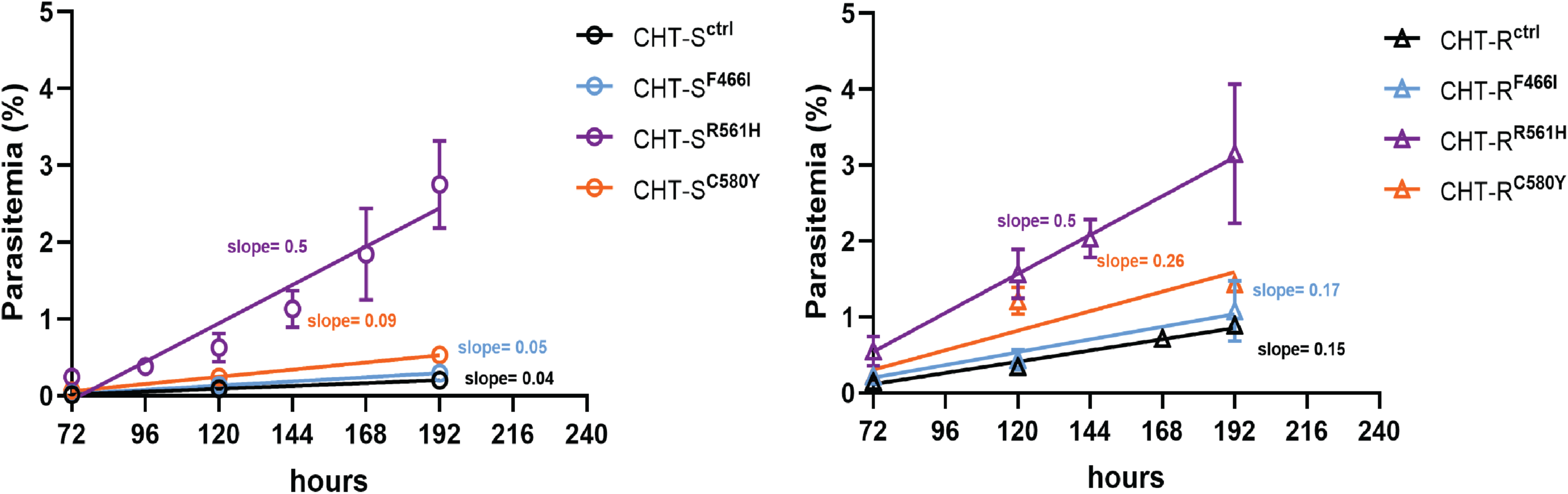
Post DHA-treatment recovery assays in the edited lines show differential impact of different mutations in recovery. Linear regression analysis of growth of K13 edited lines in CHT-S (left) and CHT-R (right) starting at 72h till 192h post treatment of 0-3h rings with 700nM DHA for 6 hours.

## Discussion

In line with the National Strategic Plan for Malaria Elimination (2021–2025), Bangladesh’s ongoing efforts are now intensifying towards eradicating the residual pockets of malaria transmission by 2030, particularly in the southeastern remote, densely forested Chittagong Hill Tracts persisting as stable malaria hotspots bordering Myanmar and India. A critical component of this strategy is enhancing surveillance and intervention measures to prevent the establishment and spread of artemisinin-resistant *Plasmodium falciparum* strains, ensuring the sustainability of elimination efforts(“National Strategic Plan_Malaria Elimination_Bangladesh_2021-2025.pdf,” n.d.; Haldar et al. 2023). CHTs are vulnerable to the cross-border introduction of drug-resistant malaria strains due to high population mobility through the movement of refugees and migrants(Sinha et al. 2020;

Noé et al. 2018; Wangdi, Pasaribu, and Clements 2021). ArtR associated K13 mutations have not yet been reported from the CHTs. We and others have found K13 mutations from Bangladesh at very low frequency(Nima et al. 2022; Mohon et al. 2014) (extended data 1) that are not ArtR candidate or validated markers. There’s a high likelihood of ArtR causing K13 mutations emerging and spreading factoring in 1) the historical dissemination of drug resistant mutations to previous chloroquine and sulfadoxine-pyrimethamine in this region(Mita, Tanabe, and Kita 2009) 2) independent emergence and expansion of ART selected K13 mutations in countries with similar span of ACT use(Takala-Harrison et al. 2015; Mathieu et al. 2020; Miotto et al. 2020; Lu et al. 2017; Chenet et al. 2016) and similar transmission settings(Huwe et al. 2022; Fola et al. 2017; Nkhoma et al. 2013; Kateera et al. 2016). Therefore, this study fills an important gap in knowledge and prepares the community to act if the most prevalent K13 mutations spread or emerge and expand.

Initially, ART-R was expected to emerge slowly, in the GMS, if at all, due to the combination of an artemisinin derivative with a partner drug. However, this hope waned with its first detection on the Thai-Cambodian border in 2008(Ashley et al. 2014; Dondorp et al. 2009) which over the two decades have spread widely in southeast Asia(Ménard et al. 2016; Kagoro et al. 2022).

Our meta-analysis (1997–2022) identified C580Y (5.65%) as the most prevalent *K13* mutation globally, followed by F446I (1.63%) and R561H (0.86%) (Supplementary Table S1). These findings align with a recent broader meta-analysis by Balmer et al(Balmer et al. 2024), covering a longer period (1980–2023) and a larger dataset, which similarly reported C580Y, F446I, and R561H among the top global K13 mutations.

By 2007, different validated K13 mutations were already prevalent, before increasing in Thailand, Cambodia, and Myanmar between 2007–2016(Kagoro et al. 2022) and Figure 2B. A strong selection pressure drove the KEL1/PLA1 co-lineage, harboring the K13 C580Y mutation, to dominance across the eastern Greater Mekong Subregion with migration eastward to Vietnam and separately westward to Thailand and Laos(Takala-Harrison et al. 2015; Imwong et al. 2017; Amato et al. 2018; Hamilton et al. 2019; Imwong et al. 2020). Over the next few years C580Y reached near fixation in the eastern GMS (Figure 2B) with K13 C580Y mutants displaying the classical clinical ArtR phenotypes of delayed clearance(Ashley et al. 2014; Ariey et al. 2014; “Association of Mutations in the Plasmodium Falciparum Kelch13 Gene (Pf3D7_1343700) with Parasite Clearance Rates after Artemisinin-Based Treatments—a WWARN Individual Patient Data Meta-Analysis” 2019) or 3 day parasite positivity(Ménard et al. 2016) and the *in vitro* ArtR phenotype of increased RSA survival(Ariey et al. 2014; Mukherjee et al. 2017; Witkowski, Khim, et al. 2013). Meanwhile in Myanmar and Thai-Myanmar border, C580Y emerged and spread independently on a different parasite genetic background(Imwong et al. 2017) with again evidence of clinical and *in vitro* ArtR phenotypes(Boullé et al. 2016).This is an example of a K13 mutation spreading in neighboring countries through both independent emergence and migration(Takala-Harrison et al. 2015; Imwong et al. 2017; Hamilton et al. 2019; Imwong et al. 2020). A similar scenario could occur in the CHTs. C580Y also emerged independently in Papua Guinea(Miotto et al. 2020), and Guyana associated with *in vitro* ArtR(Chenet et al. 2016; Mathieu et al. 2020) at lower frequencies.

F446I is the predominant K13 mutation in northern Myanmar (Figure 2A) and the China-Myanmar border region(Kyaw M. Tun et al. 2015; Z. Wang et al. 2015). However, its association with delayed parasite clearance remains unclear, with studies reporting conflicting evidence: no clear clinical ArtR association(Ashley et al. 2014) mild delayed parasite clearance (Kyaw Myo Tun et al. 2016; “Association of Mutations in the Plasmodium Falciparum Kelch13 Gene (Pf3D7_1343700) with Parasite Clearance Rates after Artemisinin-Based Treatments—a WWARN Individual Patient Data Meta-Analysis” 2019) or clear *in vivo* ArtR indicators such as prolonged parasite clearance(Huang et al. 2015) and day 3 parasitemia(Z. Wang et al. 2015).Its association with *in vitro* ArtR RSA phenotype also remains controversial(Ye et al. 2016; Zhang et al. 2019).

Similarly, R561H, prevalent at 30% in northern Thailand and in a recent study published in 2025 at 100% at the Thai-Myanmar border (Srisutham et al. 2025) and 27% in North-East Myanmar, has also emerged independently in Rwanda(Uwimana et al. 2020) with strong clinical and *in vitro* ArtR phenotypes both in the GMS(“Association of Mutations in the Plasmodium Falciparum Kelch13 Gene (Pf3D7_1343700) with Parasite Clearance Rates after Artemisinin-Based Treatments—a WWARN Individual Patient Data Meta-Analysis” 2019), Rwanda (Uwimana et al. 2021; Straimer et al. 2022; van Loon et al. 2022)and Tanzania(Ishengoma et al. 2024).

Although ArtR causing K13 mutations are not yet reported from Bangladesh, we have detected K13 mutation independent *in vitro* moderate ArtR in the CHTs(Nima et al. 2022) and Table 1. CHT-R, one such K13 independent resistant clone is genomically distinct from the CHT-S, the sensitive clone (Figure 3). Both CHT-S and CHT-R cluster closely with previous Bangladeshi (from the entire Chittagong region including the CHTs) and Myanmar *P. falciparum* genomes, suggesting significant genetic similarity between these samples and they belong to a shared or closely related genetic background. The extensive use of chloroquine and sulfadoxine-pyrimethamine (SP) in Bangladesh has led to pervasive resistance among *Plasmodium falciparum* strains. Our analysis from the MalariaGen Pf7 Bangladesh dataset (2008-2017) indicate that 100% of samples (n=1310) have the K76T mutation in the PfCRT gene, associated with chloroquine resistance, and 99% have the S108N mutation in pfDHFR, conferring pyrimethamine resistance, including our CHT isolates. While CHT-S maintains a wild-type *Pfmdr1* genotype (3D7-like), CHT-R possesses the N86Y allele mutation, which could be associated with increased sensitivity to lumefantrine (the partner drug used in the treatment), mefloquine(Windle et al. 2020).

Through gene editing, RSA phenotyping analysis we provide definitive evidence that the K13 R561H and C580Y mutations can confer *in vitro* ArtR in both CHT strains. K13-R561H mutation shows the highest level of resistance in both the CHT backgrounds (Figure 4A). In the CHT-S strain, the level of resistance is comparable to the highest levels of R561H-induced resistance reported in other Cambodian and Thai isogenic backgrounds, such as Cam3.II (13.2%), Thai 2 (14.2%), and Thai 5 (12.7%)(Stokes et al. 2021). However, the extreme resistance phenotype observed in the CHT-R background, with 30% survival mediated by the R561H mutation, has not been previously reported in any K13-engineered line. Even though the fold increase of resistance mediated by R561H is lower than in the sensitive strain (4.8 fold vs 12 fold), the final resistance level is much higher, indicating that the CHT-R background can support high-level resistance, with RSA values plateauing at extreme levels. This suggests that the R561H enhances pre-existing resistance mechanisms in the CHT-R background, the effect reaches a saturation rather than being multiplicative. C580Y, also generated a higher resistance level in the CHT-R, exhibited mean survival rates of 12.2% in CHT-R, compared to 4.2% in CHT-S, the resistance levels align with resistance levels observed in Southeast Asian (4.7-9.24% survival, Figure 4B) and African backgrounds (4.8-15.5% survival, Figure 4B), with the exception of Tanzanian (F32) and Chinese (FCC1/HN) isolates, where the introduction of C580Y did not result in ArtR (Figure 4B) suggesting the importance of a genetic background for a C580Y mutation to cause resistance. In contrast, F446I did not alter susceptibility in either of the CHT backgrounds, consistent with previous findings from F446I editing in Dd2, 3D7, and FCC1/HN (Figure 4B). The only instance where F446I was associated with resistance was reported by Siddiqui *et al*., (Siddiqui et al. 2020)where editing the mutant allele back to wild type reversed resistance, reducing ART susceptibility by two-fold in a Myanmar isolate (F09A32, Figure 4B). Our findings from R561H and C580Y editing and phenotyping emphasize that Bangladesh’s genetic backgrounds could support high resistance if these key mutations emerge or are spread from the GMS.

We posited that beyond the varied resistance levels imparted by distinct K13 alleles, the fitness costs associated with these mutations might also be crucial in determining the viability and spread of these mutations in the CHTs. Head to head growth assays (as a proxy for fitness) in this study indicate that all the three K13 mutations carry minimal to no fitness costs in both the CHT isolates, which may enable these mutations to persist in the CHT population without trade-offs in growth. C580Y was fitness-neutral in both CHT-S and CHT-R backgrounds (0% fitness cost per generation), as it was reported in Cambodian (Straimer et al. 2017). In contrast, C580Y mutations typically incurred significant fitness costs in Ugandan strains(Stokes et al. 2021), suggesting that K13 C580Y may not easily spread. F446I conferred a fitness advantage in CHT-S (0.5% per generation) and had a minimal cost in CHT-R (0.04% per generation), mirroring similar findings in the African origin lab strain 3D7(Siddiqui et al. 2020) and Dd2(Stokes et al. 2021). Notably, R561H—associated with the highest RSA survival in this study—was fitness neutral in CHT-S but advantageous in CHT-R (0.5% per generation). R561H prevalent in the Thai-Myanmar border, had been shown to have improved fitness compared to C580Y in a Thai-Myanmar isolate(Nair et al. 2018). In a Tanzanian lab strain (F32), the R561H mutation has been linked to fitness costs(Stokes et al. 2021); however, this mutation is expanding over time in Tanzania. This expansion could be due to compensatory mutations in the current Tanzanian genetic background that offset these costs and enhance growth fitness. Growth of these K13 engineered lines following DHA treatment might be another proxy for determining the resilience of spread under drug pressure in the population. R561H consistently exhibited the highest recovery rates in both backgrounds followed by C580Y (Figure 6) while F446I in contrast showed minimal recovery across both backgrounds, consistent with its limited resistance profile. The CHT-R background did not have an advantage for higher recovery than the CHT-S for any of the mutations (Figure 6).

Collectively, our in vitro results indicate that recent isolates from the CHTs are capable of developing and maintaining high-level ArtR when acquiring prevalent K13 mutations like R561H and C580Y. Moreover, K13-independent ArtR backgrounds have mechanisms to sustain higher levels of resistance mediated by K13-R561H and C580Y without fitness trade-offs for these mutations. This scenario suggests that if such parasitic genetic backgrounds acquire *K13* mutations in the CHTs, they could lead to more pronounced delayed clearance in patients, robust growth of the parasites, and potential spread, contingent on the transmission dynamics within the population. Given the high-risk factors of cross-border migration and environmental similarities to the GMS, it is imperative to intensify surveillance efforts in the CHTs to manage and mitigate the spread of ArtR.

## Materials and methods

### Data collection and analysis for K13 mutations

WWARN Artemisinin Molecular Surveyor K13 data(“Artemisinin Molecular Surveyor,” n.d.) was accessed in October 2023. In total, we included data from 242 WWARN publications covering 72,375 samples. To ensure our database was as comprehensive as possible, we performed an additional literature search for relevant, recent papers published after 2020 to supplement the WWARN database. We searched the Web of Science and PubMed with the following string: “K13 OR kelch OR kelch13 OR pfkelch13) AND (half life OR parasite clearance OR resistance)”, restricting results to articles only. We also searched “Kelch13 mutations falciparum artemisinin” in Google Scholar for articles published between 2022 - 2023. This search added 10 (He et al. 2019; Dwivedi et al. 2016; Conrad et al. 2023; Chaisatit et al. 2021; Kagoro et al. 2022; Matrevi et al. 2019; Aninagyei et al. 2020; L’Episcopia et al. 2020; Mihreteab et al. 2023; X. Wang et al. 2020). We excluded articles reporting K13 double mutations without genotyping data.

### Data limitations

Data collection in GMS declined in the last 5 years as transmission went down with additional focus in Africa, hence many SEA regions do not have data in recent years. In our analysis, we did not demarcate the sample proportion of Africa and South-East Asia.

### Sample collection

Patient isolates CHT-S (I-001) and CHT-R (I-029) were collected from *P. falciparum* infected patients from the Bandarban district in the CHTs in 2018 and 2019.

### Plasmodium falciparum culture

*P. falciparum* asexual parasites were cultured at 4% hematocrit with O+ human red blood cells (Biochemed Services, Winchester, VA and Interstate Blood Bank, Memphis, TN) suspended in complete medium containing RPMI 1640 with L-glutamine (Gibco, Life Technologies.), 50 mg/L hypoxanthine (Calbiochem, Sigma-Aldrich), 25 mM HEPES (Corning, VWR), 2g/L D-glucose (Sigma), 20 mg/L gentamicin (Gibco, Life Technologies) supplemented with 0.5% AlbuMAX II and 0.22% NaHCO_3_ in 5% CO_2_, 5% O_2_ and 95% CO_2_ gas mixture(Trager and Jensen 1976). Patient isolate CHT-R (I-029) was culture adapted using the above media at 10% hematocrit.

### DNA extraction for whole genome sequencing

Adapted *P. falciparum* cultures were expanded to 5% parasitemia with mature stages. RBCs were lysed with 0.05 % (w/v) saponin, washed with 1X PBS and genomic DNA was isolated after RNAse treatment from the parasite pellets. Genomic DNA was extracted using the QIAamp DNA Blood Mini Kit (Qiagen). The quantity and quality of gDNA was quantified by Nanodrop and Tapestation using the gDNA tape. 1-5 ug of gDNA was used for library preparation.

### Whole genome sequencing and variant calling

Whole genome libraries of the isolates were prepared and sequenced across one lane (along with 13 other samples not reported in this manuscript) of an Illumina NextSeq 2000 P2 (300 cycle) flow cell at the Genomics and Bioinformatics Core Facility, Notre Dame (https://genomics.nd.edu). Libraries were prepared using the NEBNext Ultra II DNA Library Prep kit with Covaris sonication. Libraries were sonicated to an average size of 350bp. Libraries were quality assessed using a combination of KAPA HiFi qPCR, TapeStation DNA HS Assay, and Qubit HS DNA assays. Libraries were pooled in equimolar amounts for sequencing on a NextSeq 2000 P2 (300 cycle) flowcell using paired 150bp reads. Secondary analysis was done using Illumina’s onboard DRAGEN software for BCL to Fastq conversion.

### Read alignment and variant calling

We followed the analysis from MalariaGEN Pf7 bioinformatics methods(MalariaGEN et al. 2023). The fastq files were trimmed and adaptors were removed using trimmomatic. The QC controlled reads were aligned to *Plasmodium falciparum* 3D7 reference genome using bwa mem with default parameters. The sam files were indexed and converted to bam files using samtools. GATK pipeline was used to remove PCR duplicates and Base Quality Score recalibration. GATK - HaplotypeCaller was used for variant calling using default parameters except the – sample_ploidy 1 parameter. All hypervariable, centromeric, and subtelomeric regions were discarded, and only the 21 MB core genome was considered for downstream analysis. Joint genotyping was performed using the – GenotypeVCF command. Hard filtering was used to remove low quality variants (DP < 20). SNPs and INDELS were separated with GATK -- selectVariant command. SNPs and INDELS were separated. SNPs were annotated using SnpEff. SNPs in candidate genes were evaluated. Only high quality biallelic SNPs in the core genome were used for downstream analysis.

### Principal Component Analysis

Principal Component Analysis of genomes from Myanmar, Bandarban and Ramu in CHTs, Bangladesh (MalariaGen Pf7) and our CHT samples. Biallelic core SNPs from Pf7 genomic data of selected samples (metadata and genotype matrix are included in Supplementary tables S3 and S4 respectively) were extracted by bcftools using Plink after linkage pruning with a 10 bp window step size for 50 kb window with a linkage threshold (r^2^)< 0.1. The % variance was explained by converting the ratio of eigenvalues to the sum of the eigenvalues to %. Principal Component Analysis was done using Python 3.0 (details in the Codes and data availability section). on 696 samples from seven countries (Supplementary table S3), namely from seven countries: Thailand (n_total_ = 33), Cambodia (n_total_ = 154), Bangladesh (n_total_ = 93), Vietnam (n_total_ = 95), Myanmar (n_total_ = 198), India (n_total_ = 100), Indonesia (n_total_ = 21) and newly isolated CHT-S (I-001) and CHT-R (I-029) collected. Yearwise breakdown of samples from each country; Bangladesh (n_2008_ = 13; n_2009_ = 15; n_2012_ = 49; n_2016_ = 4; n_2017_ = 12); Cambodia (n_1993_ = 5; n_2008_ = 1; n_2009_ = 2; n_2012_ = 18; n_2013_ = 2; n_2016_ = 60; n_2017_ = 66); India (n_2016_ = 30; n_2017_ = 64; n_2018_ = 6); Indonesia (n_2015_ = 7; n_2017_ = 14); Myanmar (n_2011_ = 9; n_2012_ = 33; n_2013_ = 11; n_2014_ = 29; n_2015_ = 14; n_2016_ = 46; n_2017_ = 56); Thailand (n_2001_ = 2; n_2002_ = 2; n_2003_ = 1; n_2004_ = 1; n_2005_ = 4; n_2007_ = 1; n_2011_ = 8; n_2012_ = 3; n_2017_ = 11); Vietnam (n_2000_ = 1; n_2009_ = 9; n_2010_ = 1; n_2012_ = 10; n_2015_ = 8; n_2016_ = 5; n_2017_ = 55; n_2018_ = 6).

### CRISPR/Cas9 editing of *K13* mutations

Editing of the K13 was performed using the pDC2-coSpCas9-k13guide-gRNA-h*dhfr* all-in-one plasmid that contains a *P. falciparum* codon-optimized Cas9 sequence, a human dihydrofolate reductase (h*dhfr*) gene expression cassette (conferring resistance to WR99210) and donor template (containing the respective nonsynonymous and shield/binding site mutations or only shield/binding site mutations) (plasmid generously provided by Dr. David Fidock)(Stokes et al. 2021; Adjalley and Lee 2022). 80uL of packed RBCs containing 5-8% ring parasitemia were electroprated with 50ug of the purified plasmid resuspended in Cytomix using a BioRad Gene Pulser Xcell Electroporation System. Transfected parasites were maintained under 5 nM WR99210 (Jacobus Pharmaceuticals) to select for edited parasites till 6 days post-transfection, after which the drug was removed from the media. Parasite cultures were monitored for recrudescence microscopically for up to six weeks post electroporation.

### Validation of K13 edits-PCR and sequencing

K13 edit was validated by both direct PCR using Phusion^®^ Blood Direct PCR Kit (ThermoFisher Inc.) and from genomic DNA isolated from the bulk transfected cultures and their clones. For 2uL culture/25ng genomic DNA was added as the template in a 20uL reaction. PCR product was purified using QIAquick PCR Purification Kit (Qiagen) and Sanger sequenced.The primers used for amplification and sanger sequencing are: k13-1270-Fw: GAAAGTGAAGCCTTGTTGAAAGAAGCAG and k13-UTR-rev: aaatgtgcatgaaaataaatattaaagaag

### Cloning of parasites by limiting dilution

Transfected K13-edited bulk culture or the CHT isolate (I-029) was diluted and low density seeding was performed on a 96 well plate at 0.25 parasites per well as previously described(Button-Simons et al. 2021) . Plates were maintained in a gassed incubator for a month with media changes and screening every week for parasite-positive wells. Screening was performed by qPCR using the Phusion Blood Direct PCR kit (ThermoFisher Inc.), supplemented with 1 × SYBR, beginning week 2 and continuing on weeks 3,4 and 6. Two microliters of culture was used in a 10 μl reaction and amplified using forward and reverse primers of the *pfcrt* gene. PCR amplification was measured using the ABI 7900HT, with a 5 min denaturation at 98 °C, followed by 40 cycles of 95 °C for 1 s, 53 °C for 5 s, and 65 °C for 15 s, then final extension at 65°C for 60s. Primer used for amplification are CRT-F: GGGTGATGTTGTAAGAGAACCA and CRT-R: ACGAACAAGCCATTTGATATTA.Positive wells with a CT score of 25 or lower were pulled to 1 mL wells at 5% Hct in 24 well plates for microsatellite analysis, expansion, cryopreservation and characterization.

### Ring Stage Survival Assay

The RSA_0–3h_ was performed as previously described(Mukherjee et al. 2017). Late-stage segmented schizonts, purified with Percoll, were briefly cultured with fresh red blood cells for 3 hours, followed by sorbitol synchronization. Cultures containing 0 to 3-hour rings were adjusted to a 2% hematocrit and 1% parasitemia and seeded into a 24-well plate with 2 mL complete medium per well, containing either dihydroartemisinin (DHA) at 700 nM or 0.1% dimethyl sulfoxide (DMSO) as a control. After 6 hours at 37°C, the cultures were washed and transferred to drug-free medium. Subsequently, after 66 hours (equivalent to 72 hours from initial seeding), thin blood smears were prepared, and survival rates were microscopically assessed by counting the proportion of viable rings with normal morphology after staining with giemsa. Parasitemia was estimated based on counts from 10,000 red blood cells in controls and 20,000 RBCs in treated. Survival % rates were determined by multiplying the ratios of viable parasitemia in DHA-exposed parasites to DMSO-treated controls by 100 and expressed as a percentage, with a survival rate ≥1% indicating the threshold for resistance(Witkowski, Amaratunga, et al. 2013).

### Post DHA Treated Growth Assay

Parasitemia for each edited line was estimated in parallel across at least three biological replicates over 8 days. Cultures were treated with 700 nM DHA at 1% parasitemia and 2% hematocrit in 24-well plates, and parasitemia was measured by microscopy.

### Pairwise head-to-head competitive growth assay

We conducted head-to-head pairwise competition growth assays between each K13-edited line with their corresponding isogenic control transfected line ( K13 shield mutations). We performed these co-culture assays in 96 well plates by mixing the lines at 1:1 ratio of synchronized rings, each strain at 0.5% with a total of 2% hematocrit in 200uL culture/well and measuring the proportion of one strain to another (sampled every 2 days) over a period of 40 days following established protocols from the Ferdig Lab (Tirrell et al. 2019; Kumar et al. 2022). Synchronized ring-stage parasites were used to initiate the assay (Day 0). At regular intervals (Days 0, 2, 4, 6, 8, 10, 12, etc.), a portion of each culture was collected and stored at –20°C for genotyping. Removed volume was replaced with fresh complete media and RBCs to maintain a consistent 200 μL volume. They were performed in 3 biological replicates with 3 technical replicates each in a co-culture competition experiment. Cultures were maintained in a gassed incubator with media changes or splitting every other day for 40 days. K13 Genomic gene was amplified using the Phusion RBC PCR kit using K13-1270-fw:GAAAGTGAAGCCTTGTTGAAAGAAGCAG and K13-UTR-rev: aaatgtgcatgaaaataaatattaaagaag primers. PCR products were quantified and diluted to 5 ng/μL, and 2 μL (10 ng) was used per reaction. Growth of the lines with different K13 SNPs in the same background were measured by qPCR using SNP genotyping RhAmp SNP kit (IDT) (Amambua-Ngwa et al. 2023) with primers targeting the K13 edited region, in a 10 μL reaction volume, including 5.3 μL of combined Genotyping Master Mix and Reporter Mix (20:1 ratio), 0.5 μL of 20X SNP Assay, 2.2 μL water, and 2 μL of PCR product. qPCR was run on an ABI system using the SNP genotyping protocol. Allele-specific primers were designed using IDT’s rhAMP genotyping tool and include universal primer binding tails (GT1, GT2, GT3, and GT4) required for amplification with rhAMP Master Mix. Mutant alleles were detected via the VIC channel and wild-type via FAM. Ct values were used to calculate fold change (2^ΔCt, where ΔCt = VIC – FAM), and percent competition win was calculated as:

% won = (fold change / (fold change + 1)) × 100

Day 0, 10, and 20, 30 and 40 samples were used for win/loss analysis.

Primers used for this study:

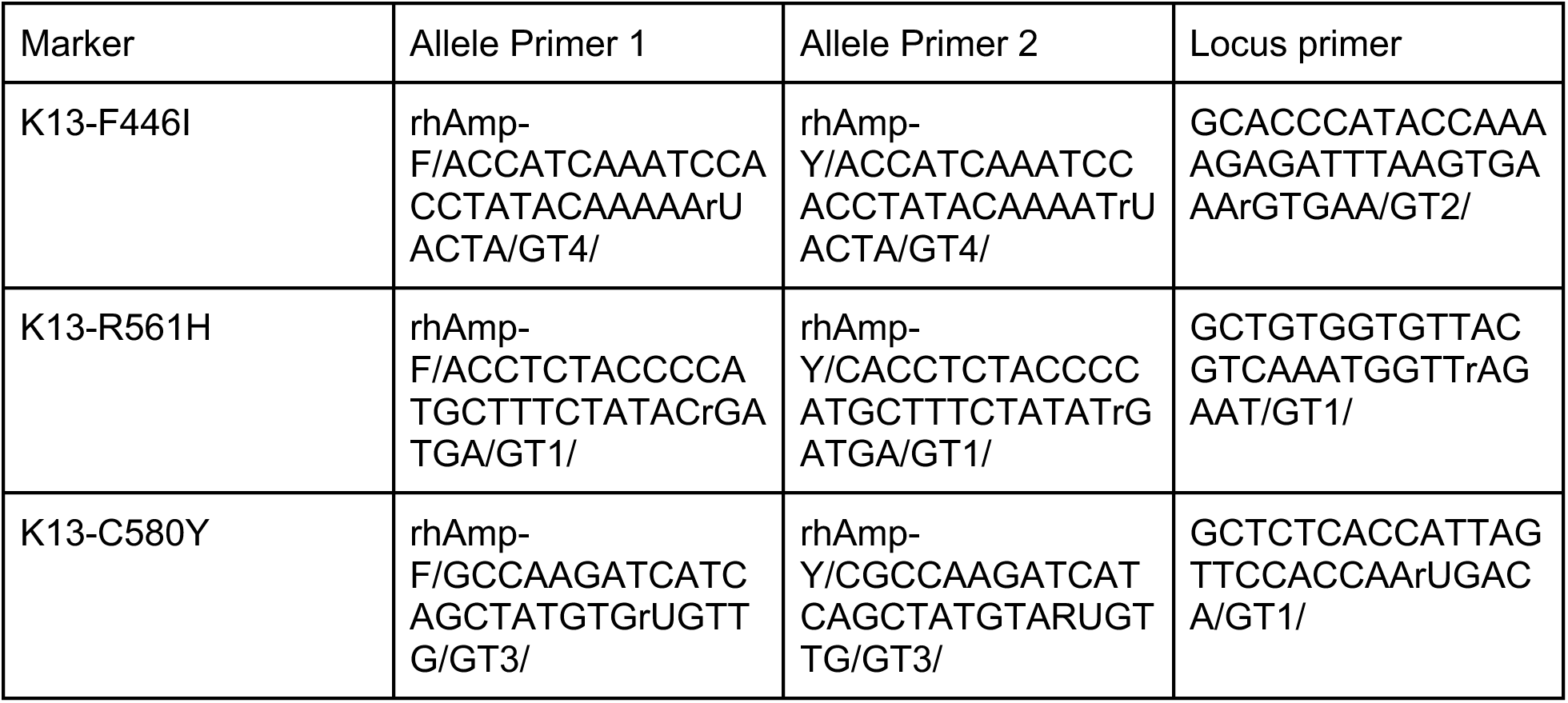

### Estimation of win/loss in pairwise competitive growth assay

Competition assays were conducted until one parasite line dominated with a population of ≥ 60% in the culture. If neither line achieved this dominance within 40 days, the competitions were halted. The winners were established at the conclusion of each competition assay. A definitive victory was declared if one line successfully outcompeted the other, reaching ≥ 60% of the total parasitaemia by day 40

## Supplementary Figures

**Supplementary Figure 1.**
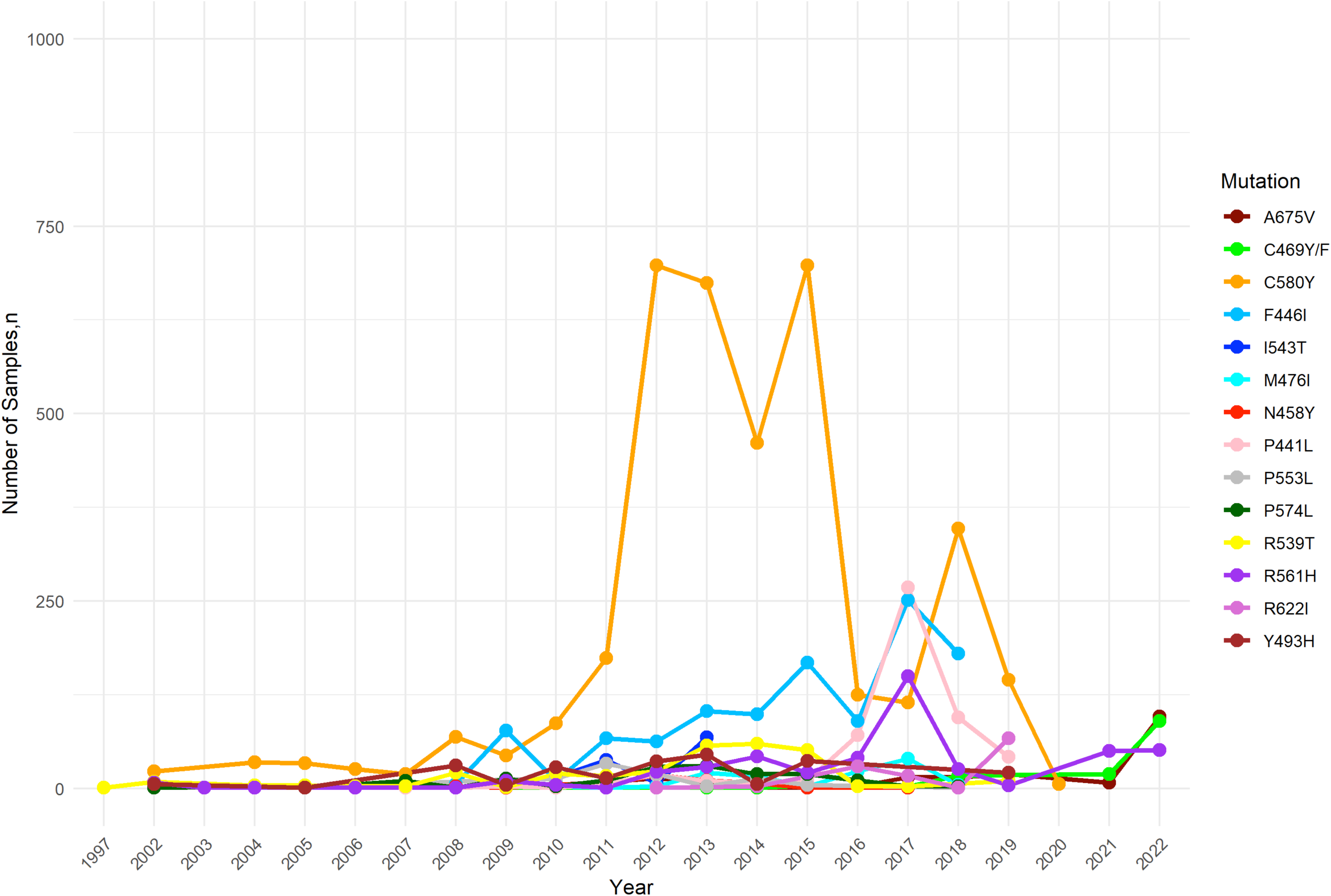
Global temporal progression of validated K13 mutations from 2007 to 2022. See Supplementary Table 1 for source data.

**Supplementary Figure 2.**
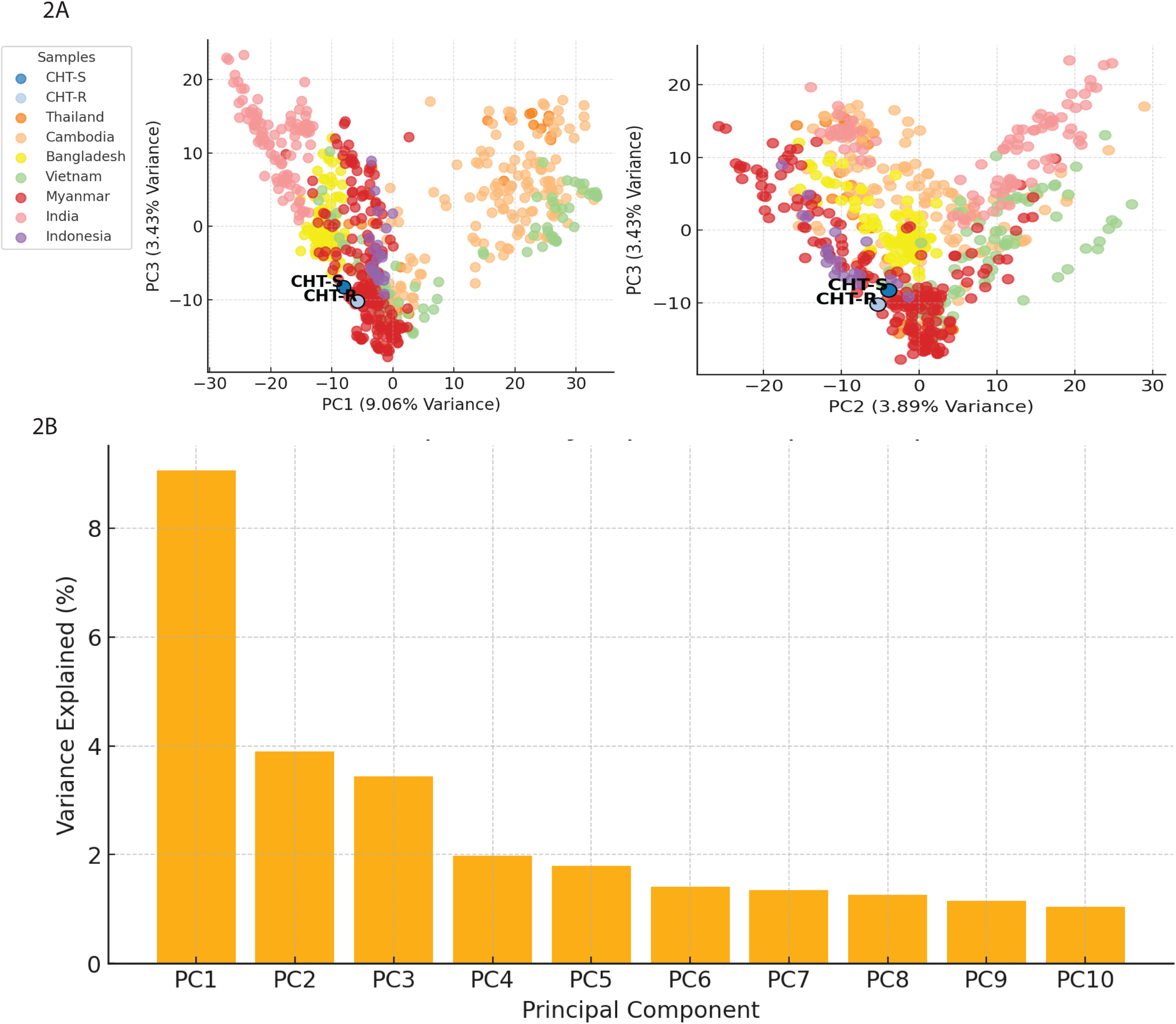
A. Pairwise Principal Component Analysis (PCA) using Principal Component 1, 2, and 3. PC1 and PC2 explain 9.06% and 3.43% variance and PC2 and PC3 explain 3.89% and 3.43% variance of the multisample genomic data. B. Variance explained in percentage (%) by the first ten principal components in the PCA.

**Supplementary Figure 3.**
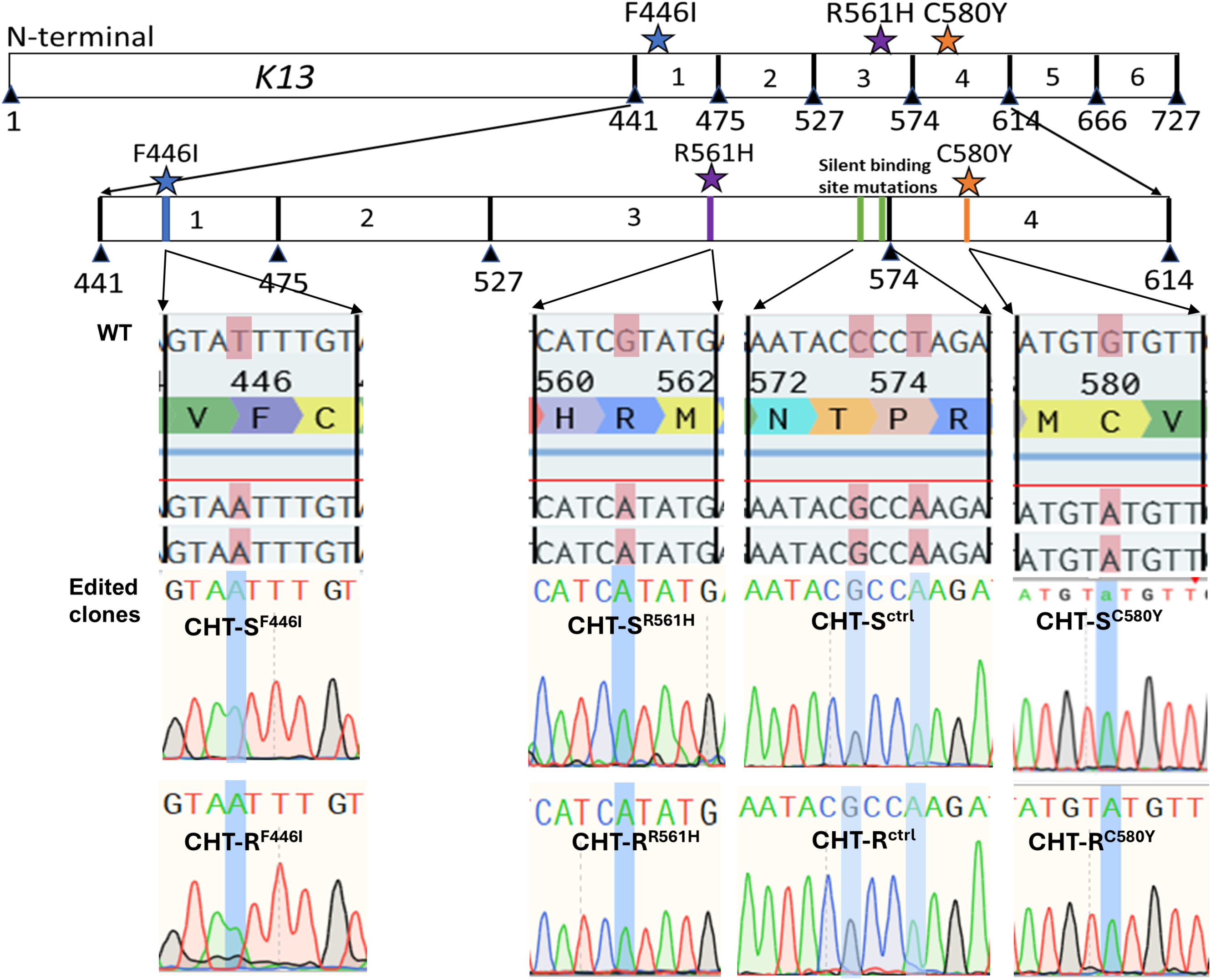
CRISPR-Cas9 genetic editing of K13 F446I, R561H, C580Y and silent binding site control mutations (cntrl) in CHT-S and CHT-R. Cartoon shows location of the K13-propeller domain mutations. Reference sequence of 3D7 on top and Sanger sequence and chromatogram analysis of an edited clone below. Highlighted bases show the WT and edited alleles of clones. CHT-S^ctrl^ and CHT-R^ctrl^ parasites contain only synonymous, phenotypically silent mutations.

**Supplementary figure 4.**
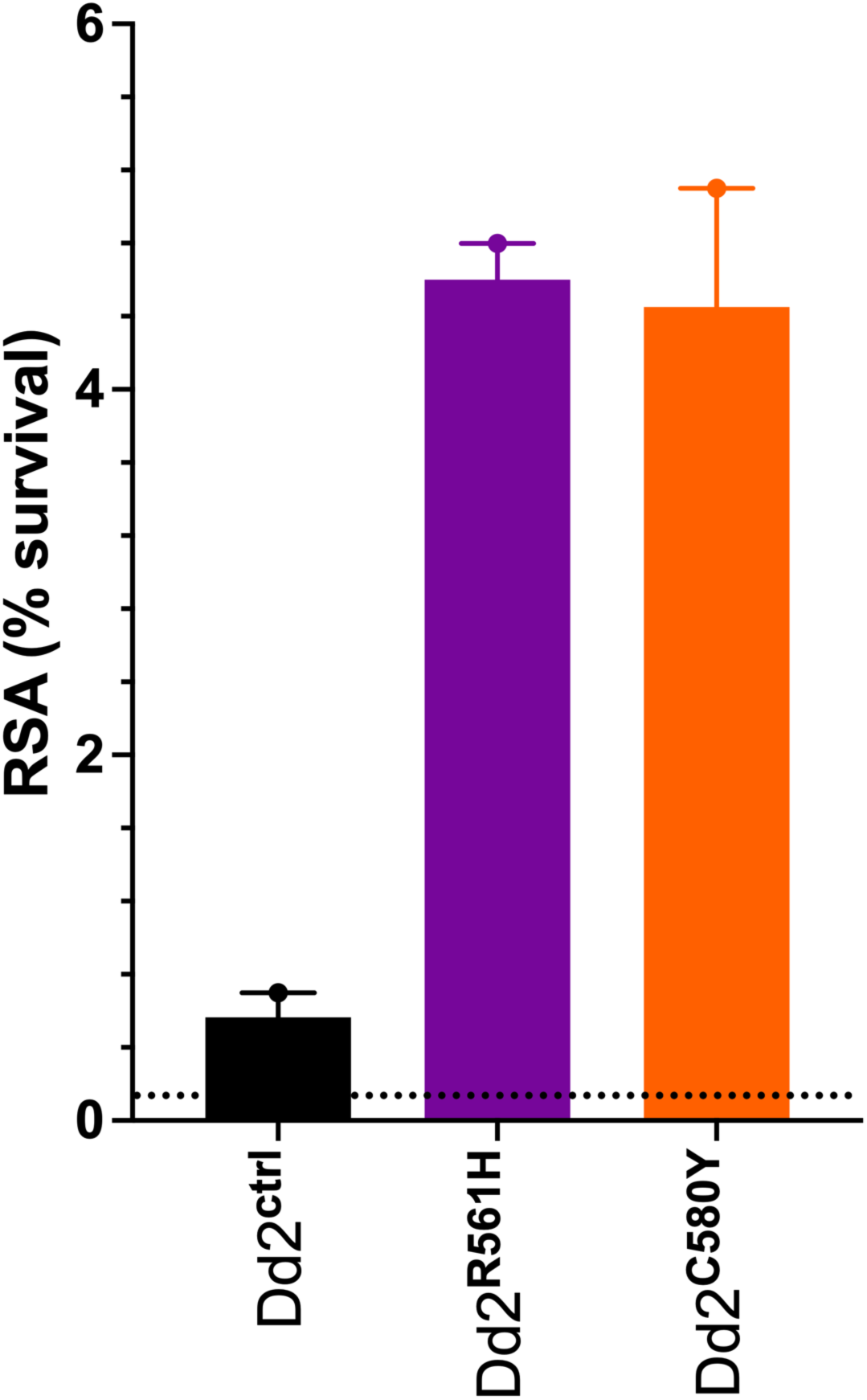
RSA survival of silent, R561H and C580Y K13 mutations in Dd2. Parasites were graciously donated by Dr. David Fidock.

## Codes and data availability

Codes used for Principal component Analysis are deposited at https://github.com/NirjharBhattacharyya/PCA_for_CRISPR_Paper-.

## Funding and Acknowledgements

This work was supported by the National Institutes of Health, NIAID (#1R21AI180663-01A1 to Mukherjee; 2024-2026), Indiana Clinical and Translational Sciences Institute (ICTSI) to Mukherjee (2023-2025) and University of Notre Dame, College of Science, Center for Rare and Neglected Diseases. M.K.N. was supported by a PhD fellowship from the Eck Institute of Global Health and College of Science (University of Notre Dame). We thank Dr. Kasturi Haldar for establishing the clinical capacity in the CHTs. We thank Dr. David Fidock for providing the K13 CRISPR constructs and the Dd2 edited strains. We thank the Genomics and Bioinformatics Core Facility of Notre Dame for the whole genome sequencing.

M.K.N. contributed to in vitro cultures, transfection and genome editing, execution of in vitro AR by RSA and IC50 assays, curation and analyses of all data, visualization of all results, and drafting and editing of the manuscript. N.B. contributed to all the bioinformatics analyses, IC50 assays, whole genome sequencing, variant calling, PCA analysis, visualization, drafting and editing of the manuscript. J.P. assisted with experimental assays. D.S. helped with the design of RhAmp genotyping assays. C.S.P collected patient isolates and performed in vivo characterization in patients. S.A.S supervised lab activity, sample preparation and initial culture establishment in the field. M.S.A supervised the study in icddr,b and donated isolates to M.K.N for dissertation research. S.K. contributed to reviewing and editing of the manuscript. All experiments were performed in M.T.F’s lab. A.M. contributed to conceptualization, study design, supervision, adaptation of field strains to *in vitro* cultures, transfection and genome editing, measurement of *in vitro* AR by RSA curation, and drafting and editing of the manuscript.

## Competing interests

The authors declare no competing interests.

---

**Supplementary Table S1:** Source Data for global temporal progression of validated K13 mutations from 2007 to 2022.

| mutation | total_mutation_samples | Total number of samples | % of mut in total samples | number of muts | %of mut in total number of validated muts |
| --- | --- | --- | --- | --- | --- |
| C580Y | 4095 | 72375 | 5.658031088 | 8210 | 49.87819732 |
| F446I | 1182 | 72375 | 1.633160622 | 8210 | 14.39707674 |
| R561H | 628 | 72375 | 0.867702936 | 8210 | 7.649208283 |
| P441L | 541 | 72375 | 0.747495682 | 8210 | 6.58952497 |
| R539T | 352 | 72375 | 0.486355786 | 8210 | 4.287454324 |
| Y493H | 254 | 72375 | 0.350949914 | 8210 | 3.093788063 |
| A675V | 214 | 72375 | 0.295682211 | 8210 | 2.606577345 |
| P574L | 203 | 72375 | 0.280483592 | 8210 | 2.472594397 |
| C469Y/F | 181 | 72375 | 0.250086356 | 8210 | 2.204628502 |
| I543T | 136 | 72375 | 0.18791019 | 8210 | 1.656516443 |
| M476I | 127 | 72375 | 0.175474957 | 8210 | 1.546894032 |
| R622I | 119.496 | 72375 | 0.165106736 | 8210 | 1.455493301 |
| P553L | 108 | 72375 | 0.149222798 | 8210 | 1.31546894 |
| N458Y | 69 | 72375 | 0.095336788 | 8210 | 0.84043849 |
| Total | 8209.496 |  | 11.34299965 |  |  |

% WT 88.65700035

**Supplementary Table S2:**
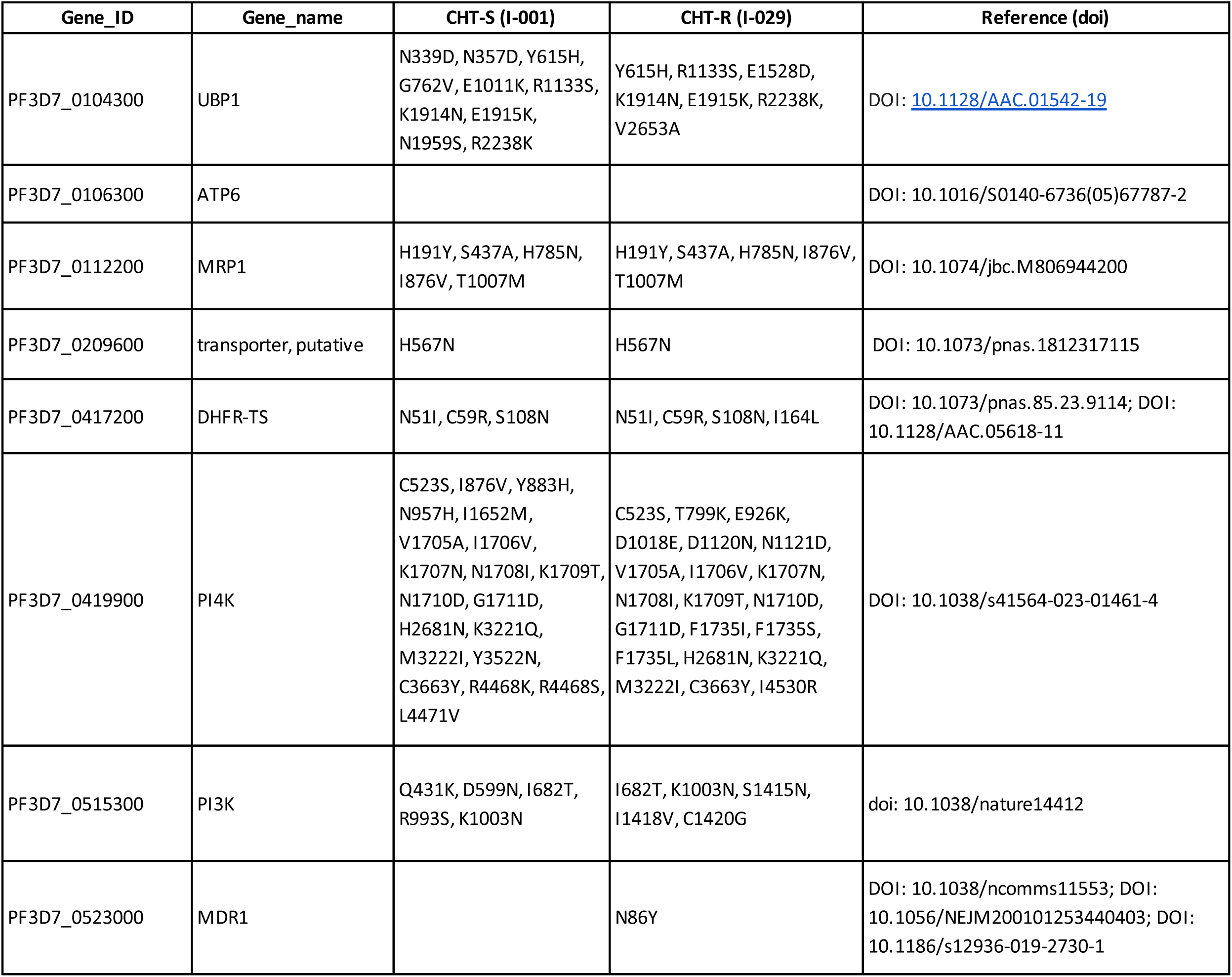
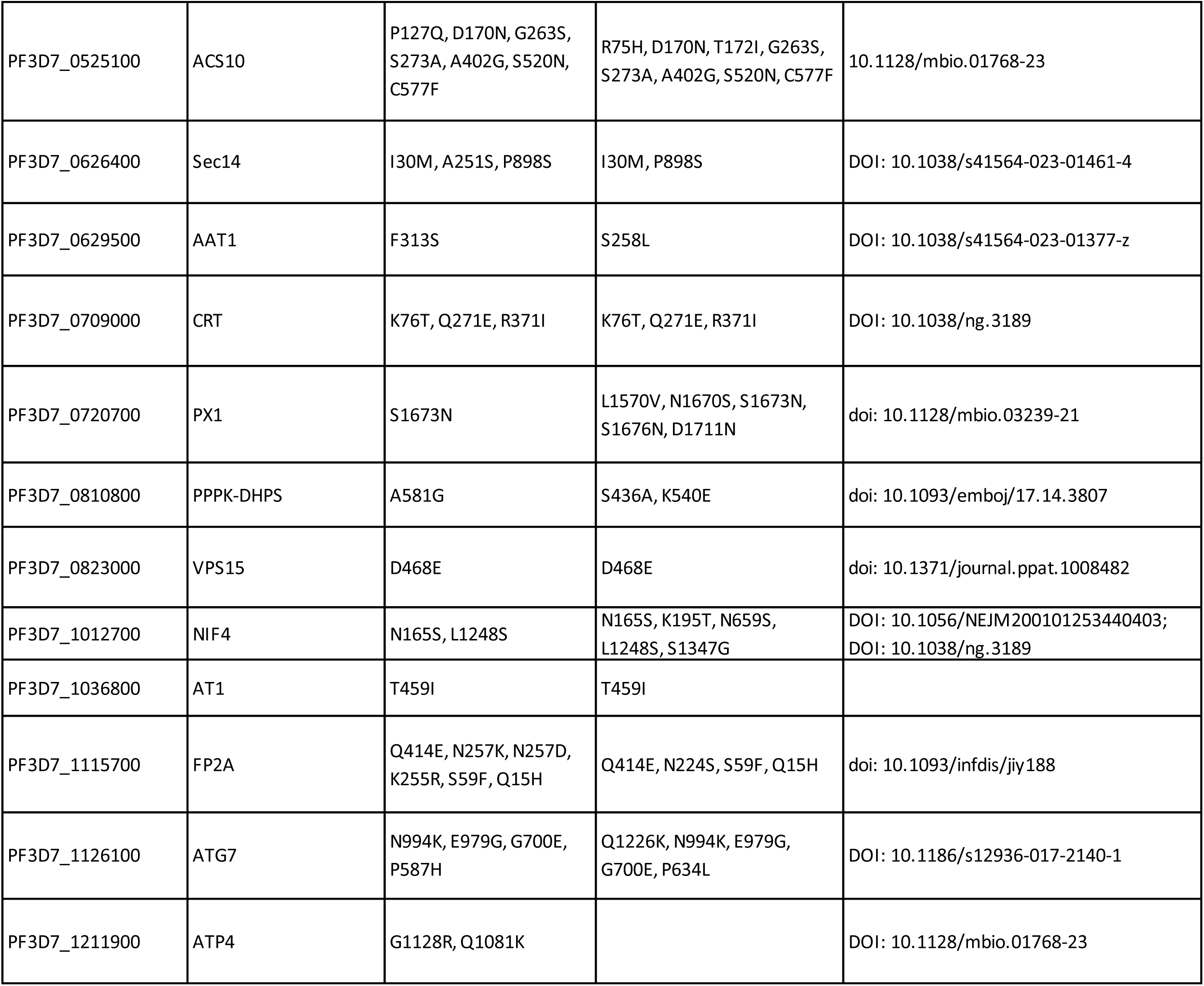
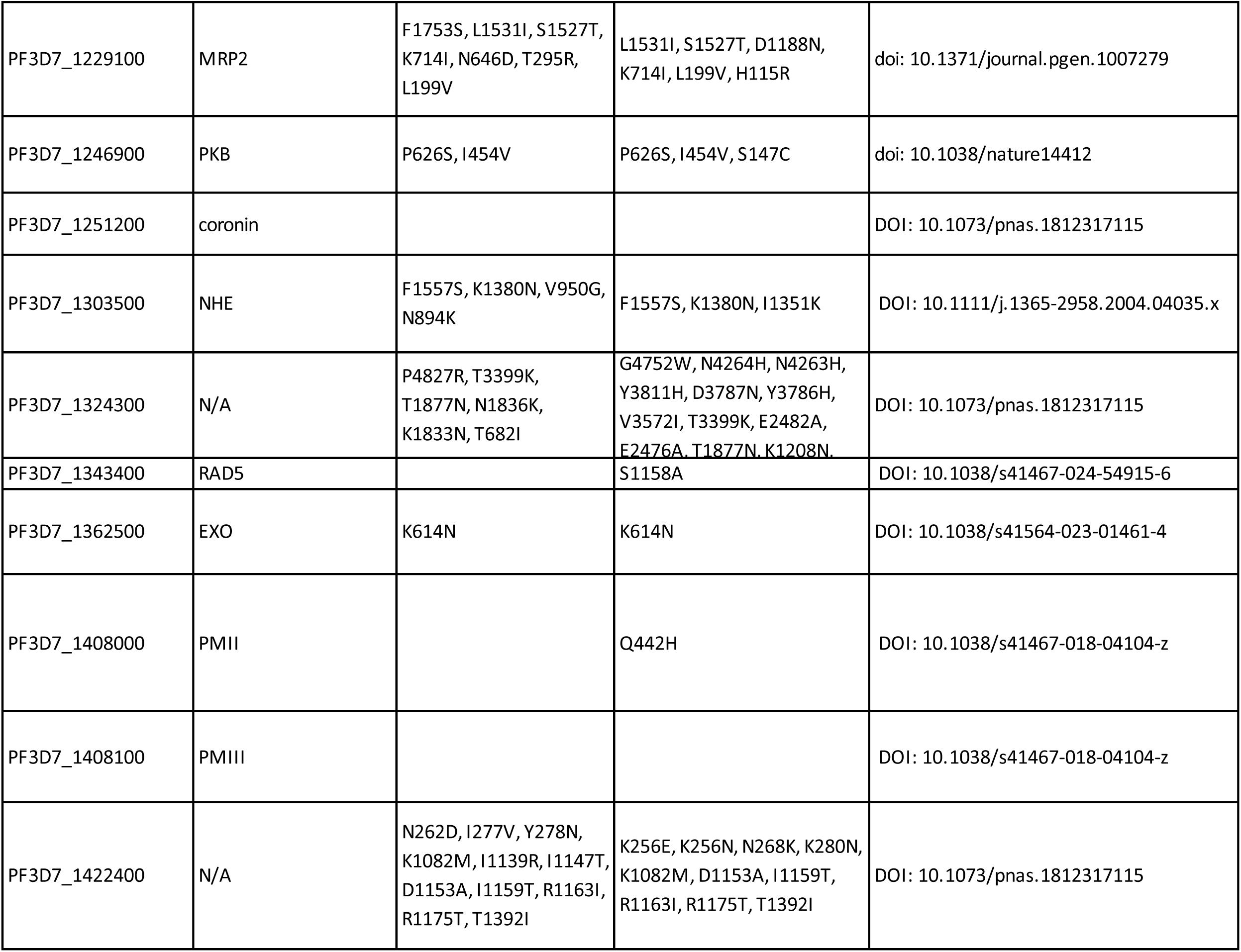
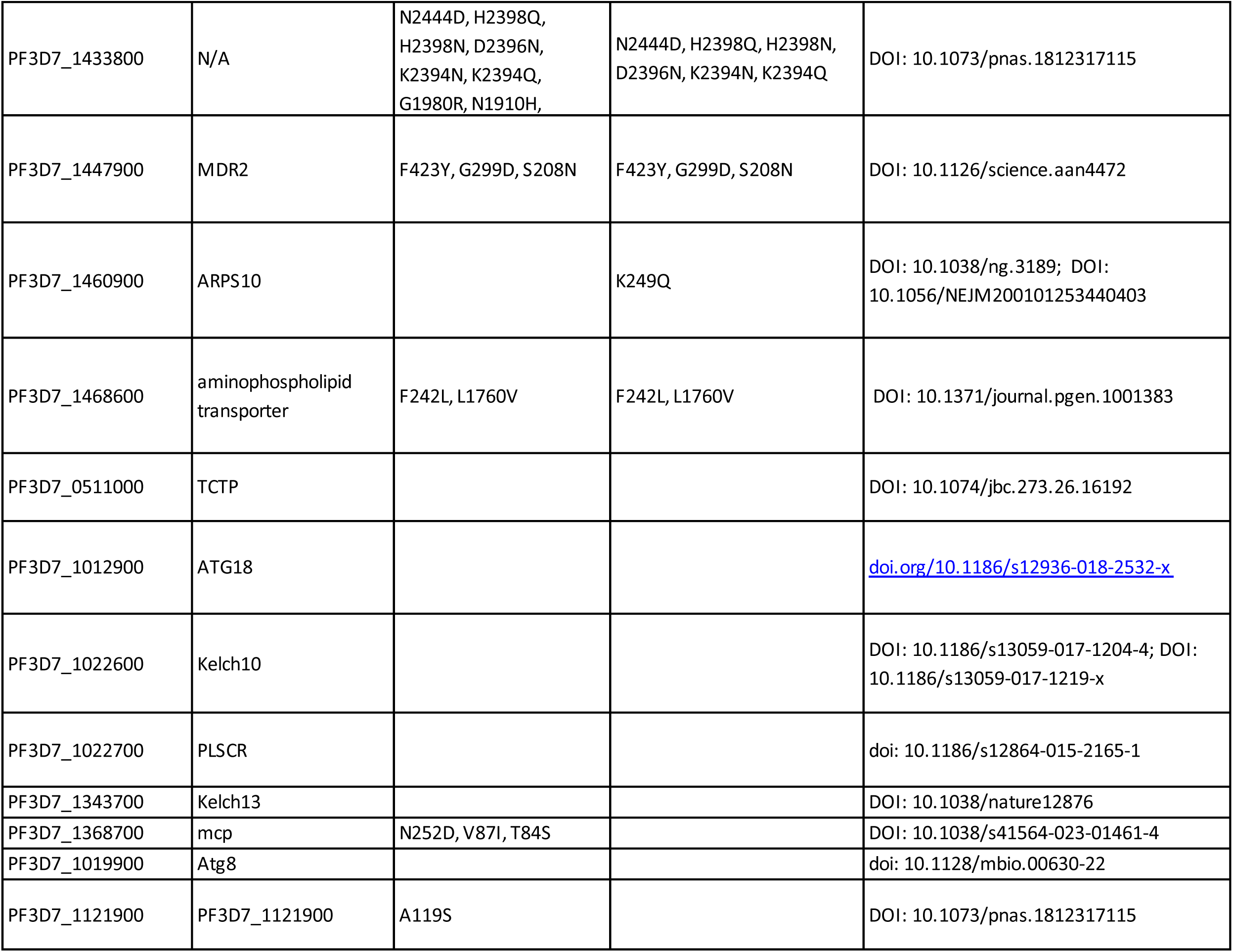
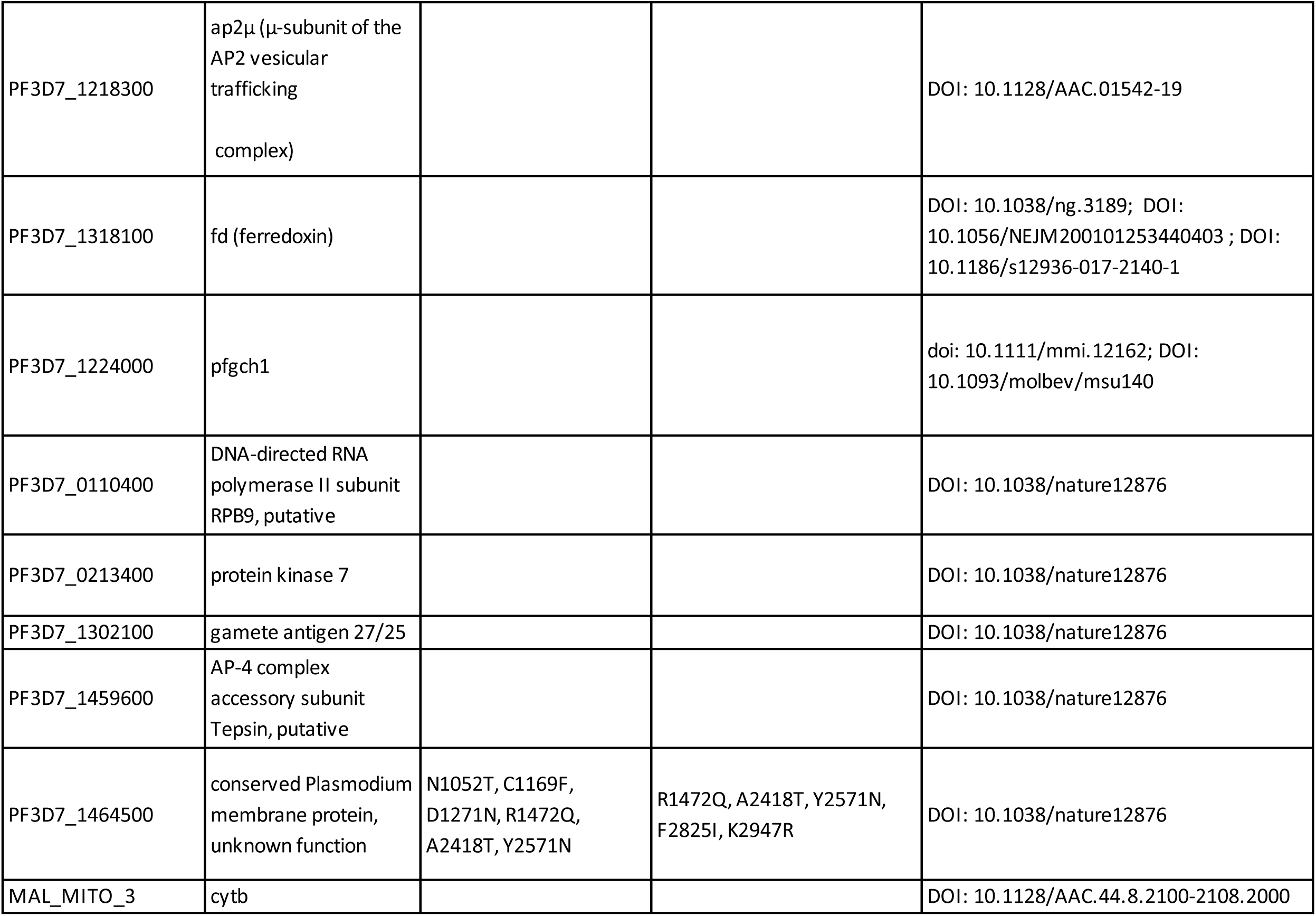
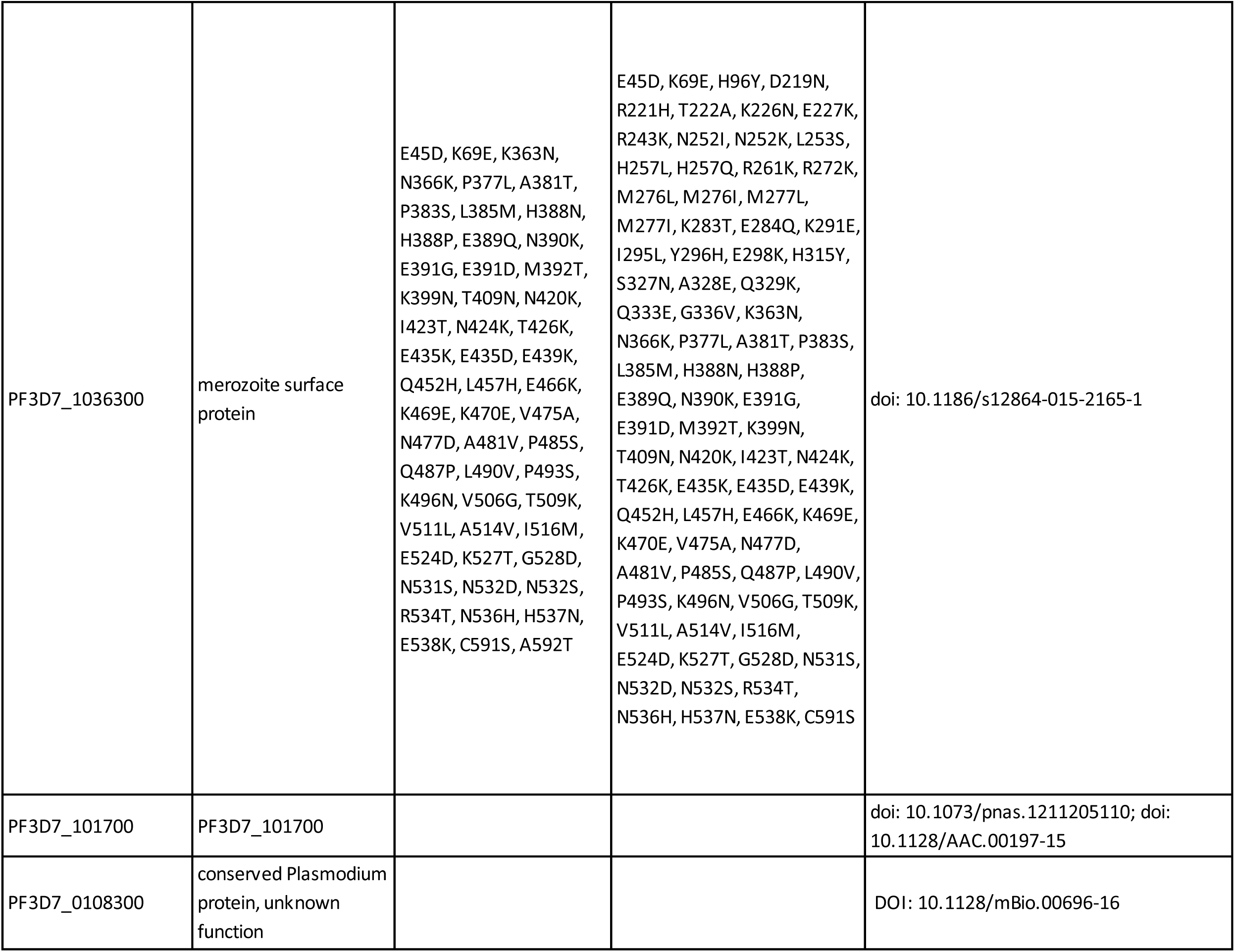
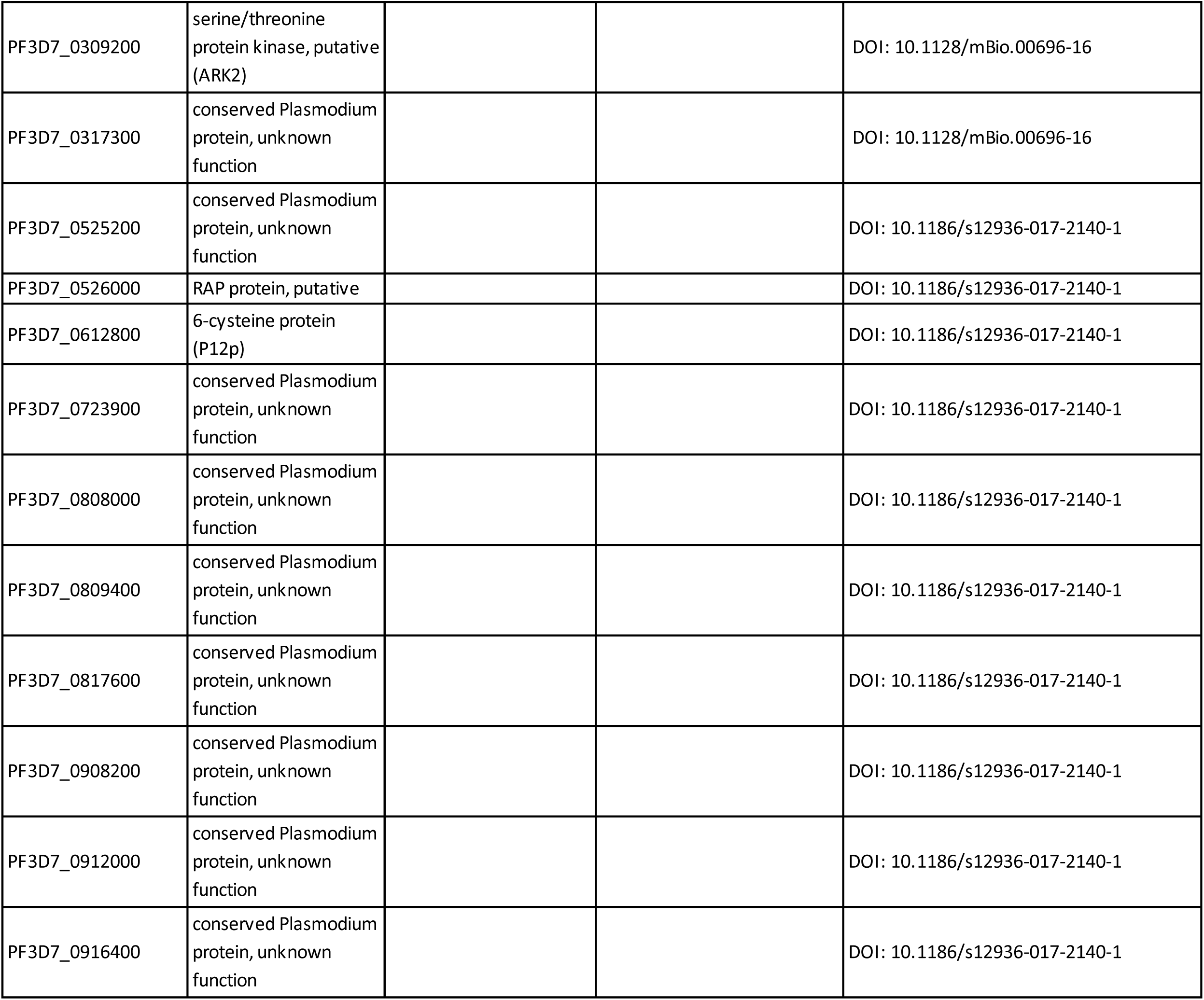
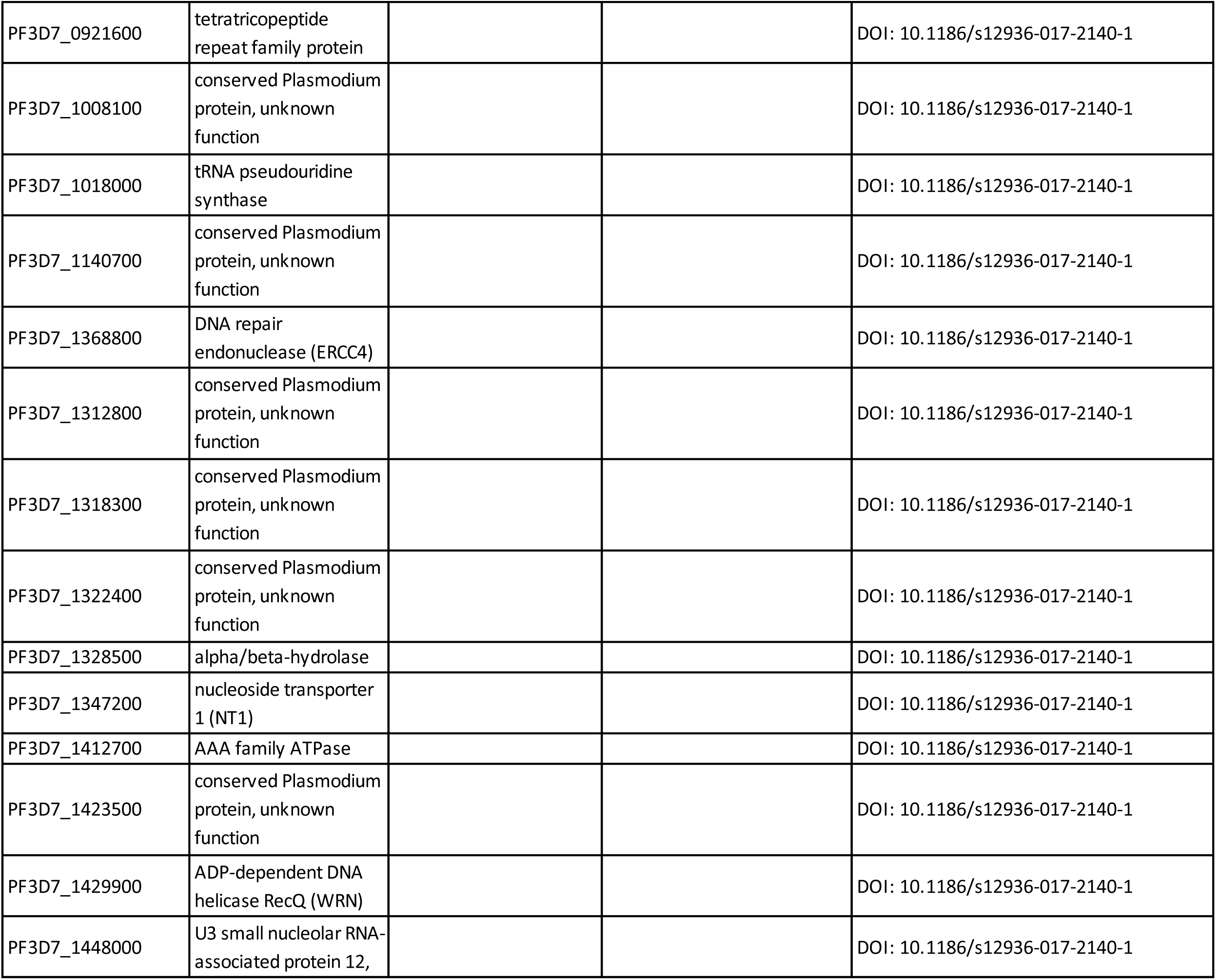
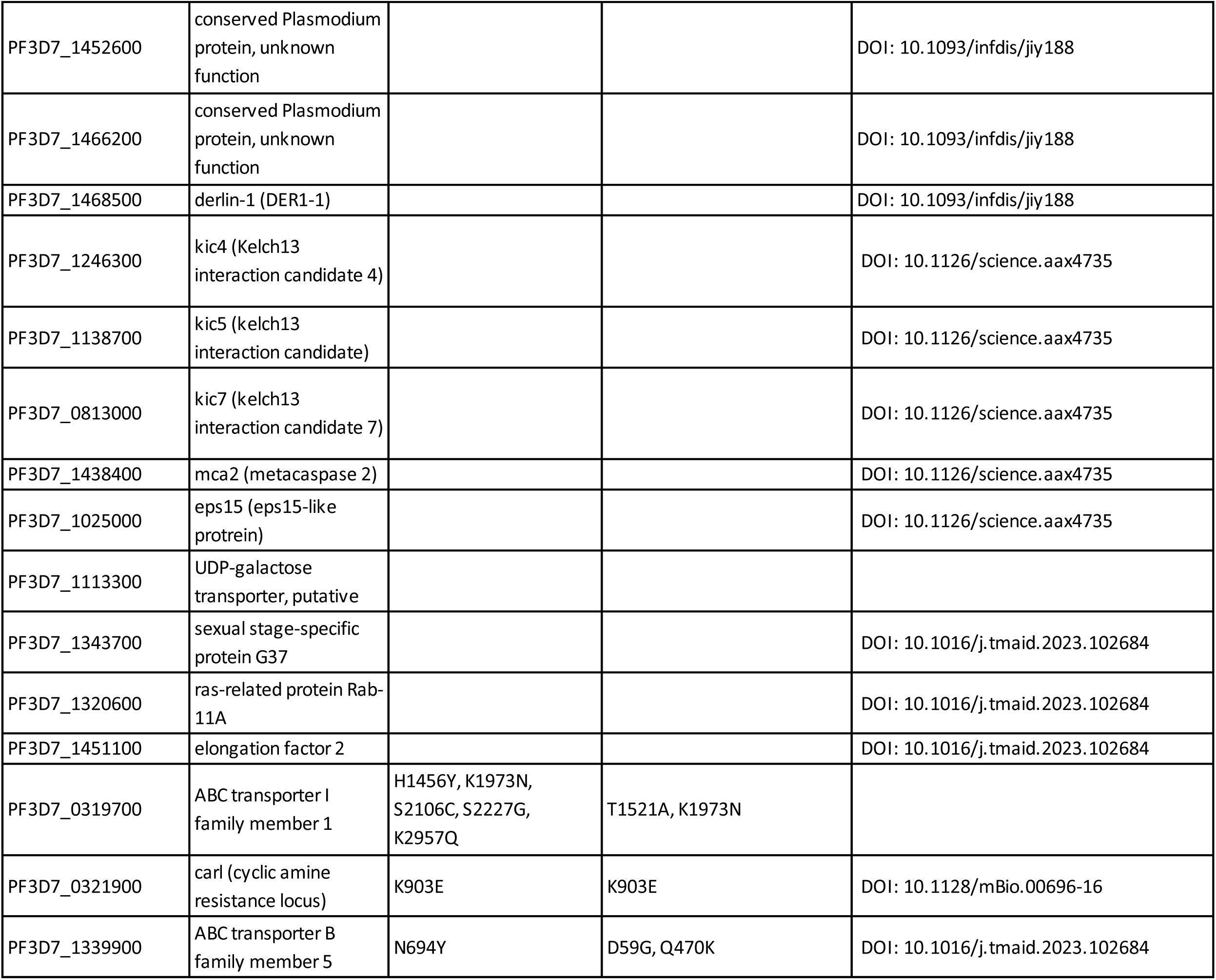
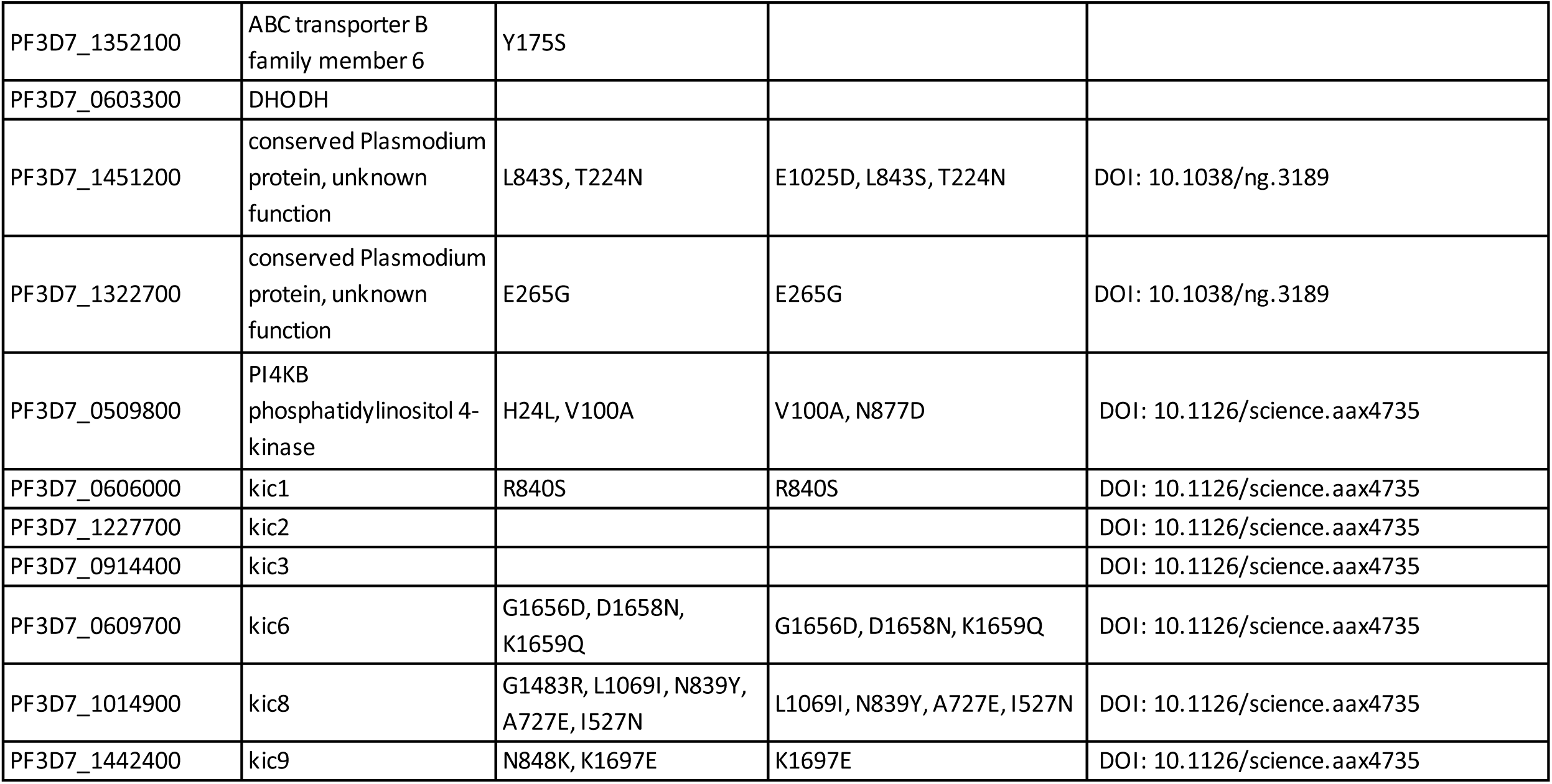
Genetic backgrounds of CHT-S and CHT-R of previously described nonsynonymous SNPs associated with ArtR.

**Supplementary Table S3:**
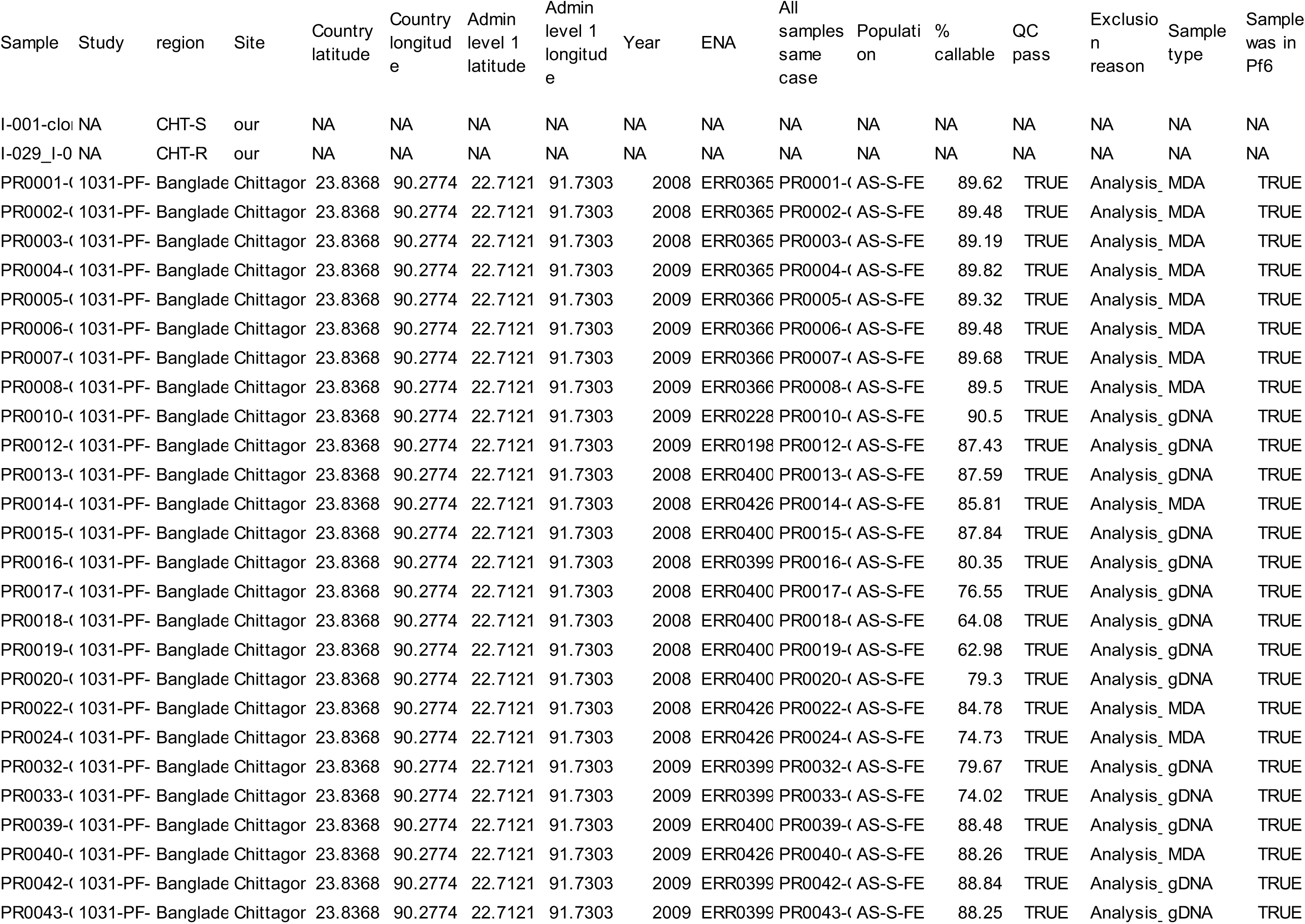
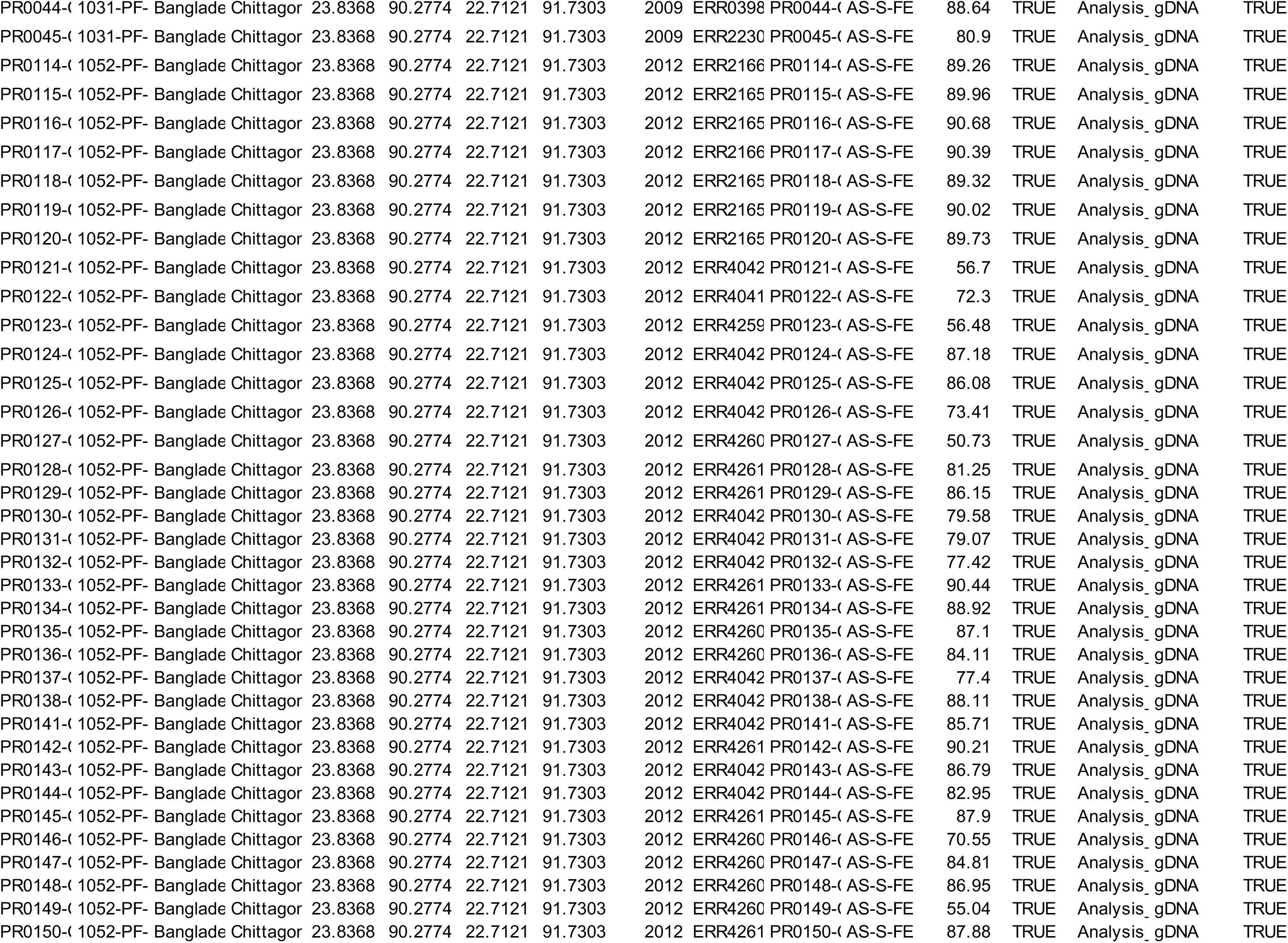
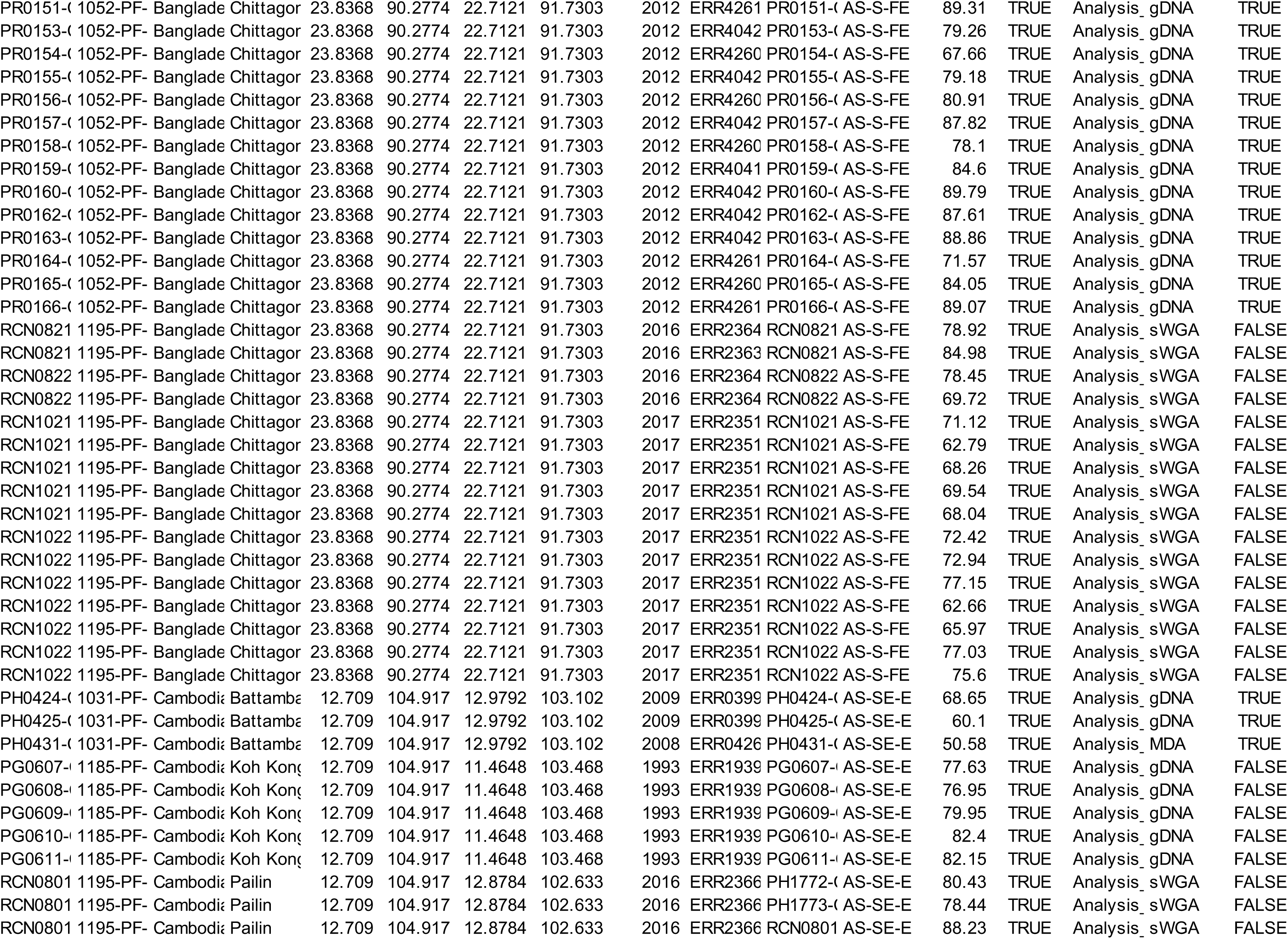
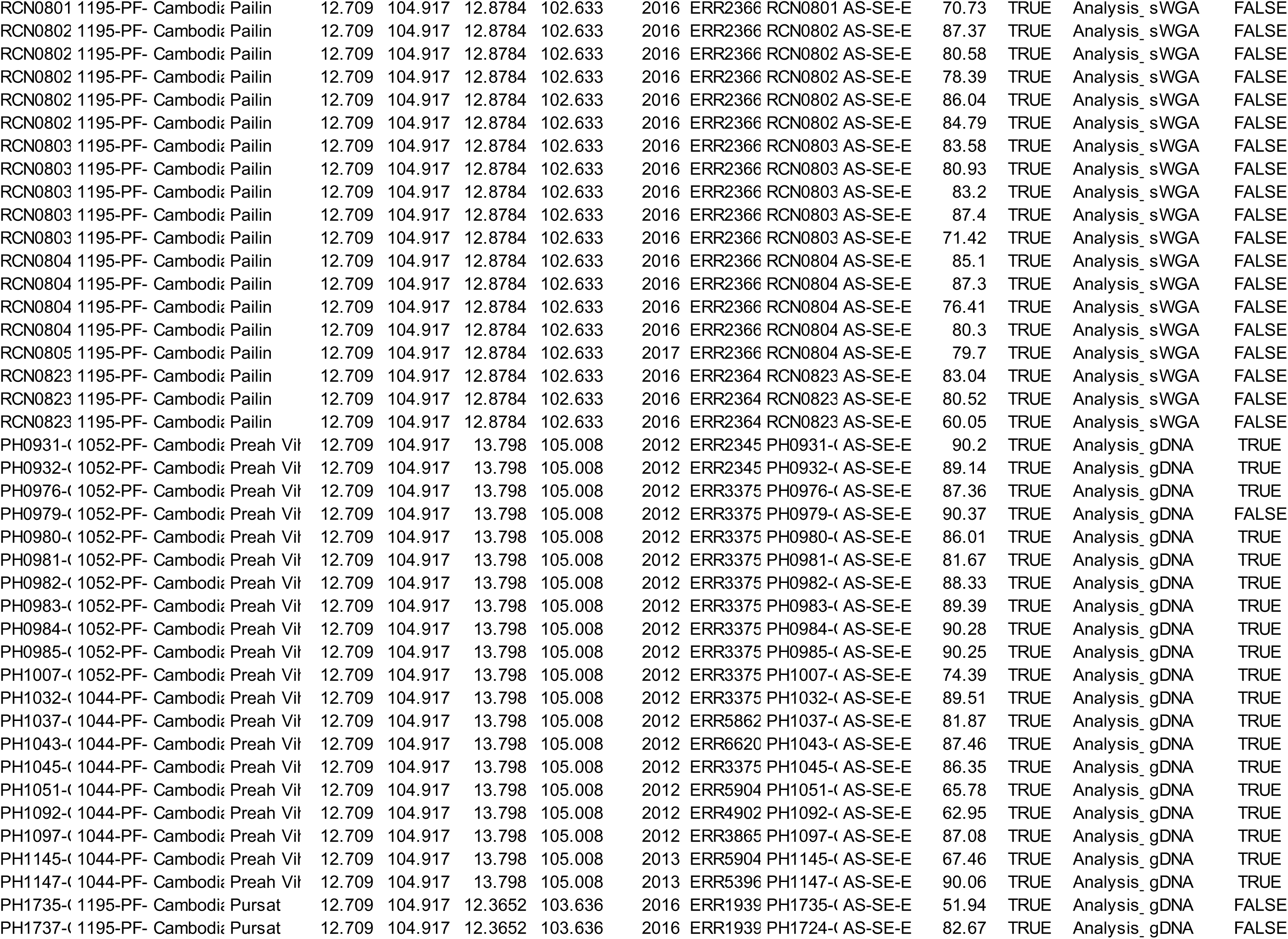
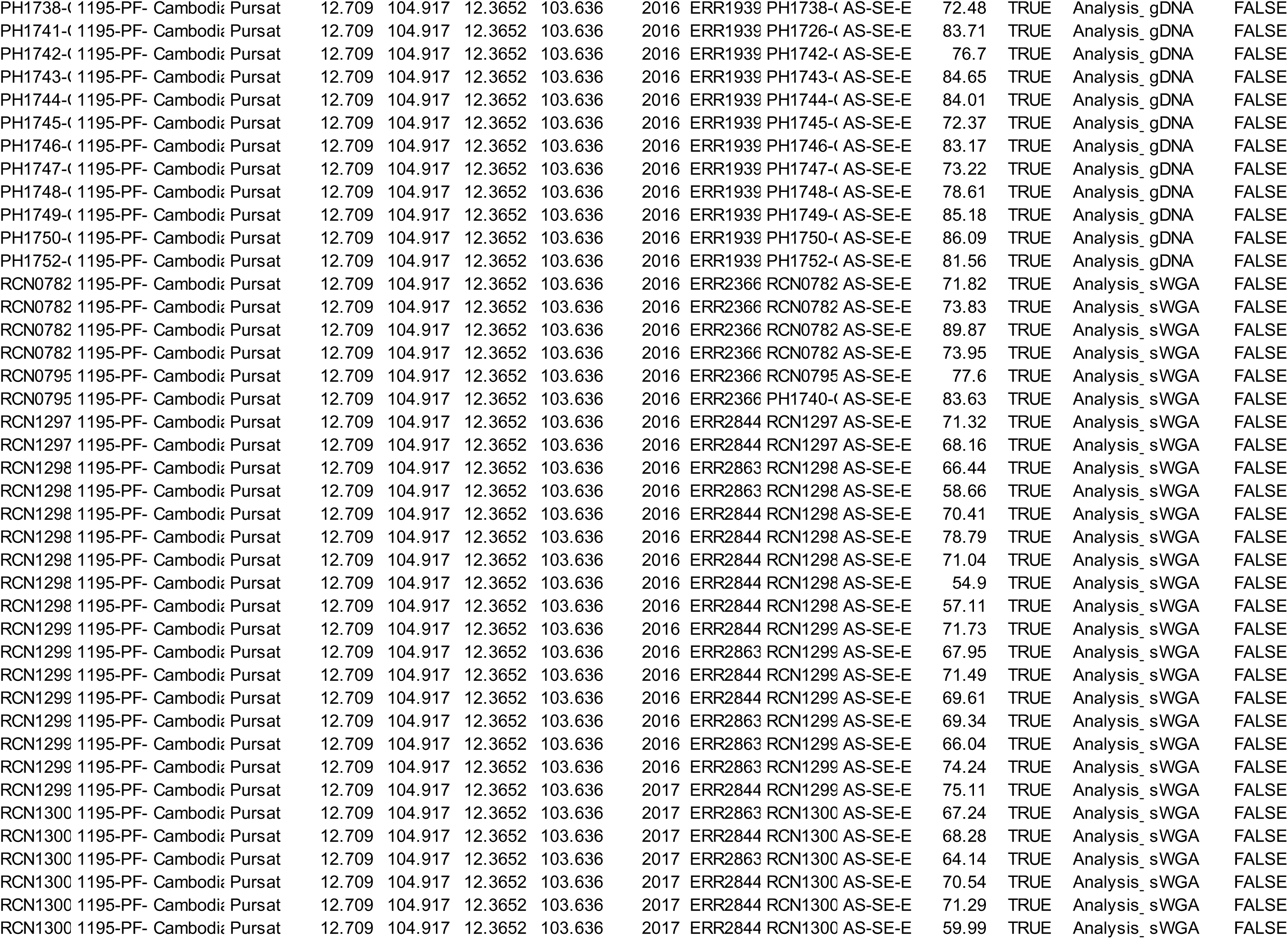
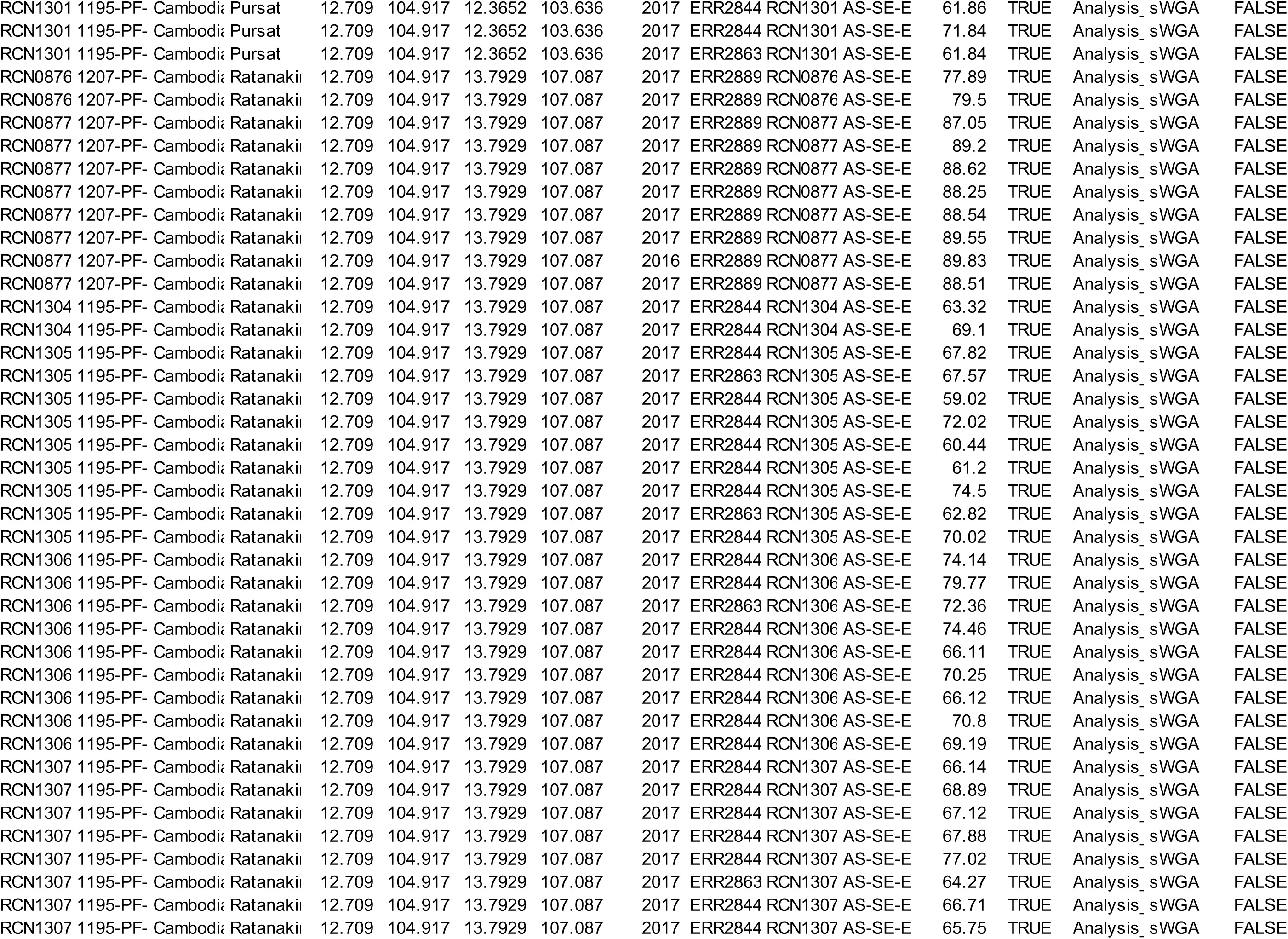
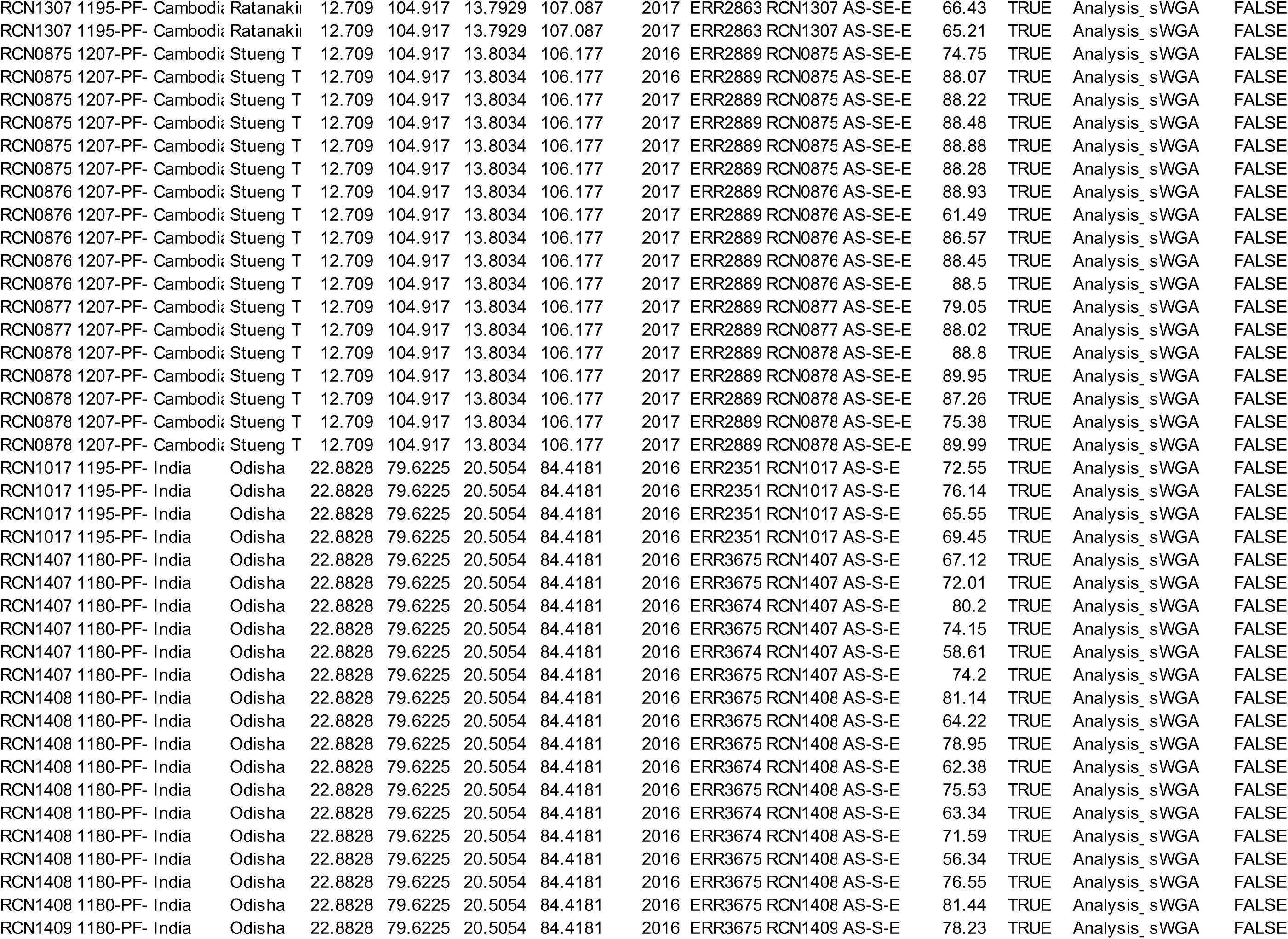
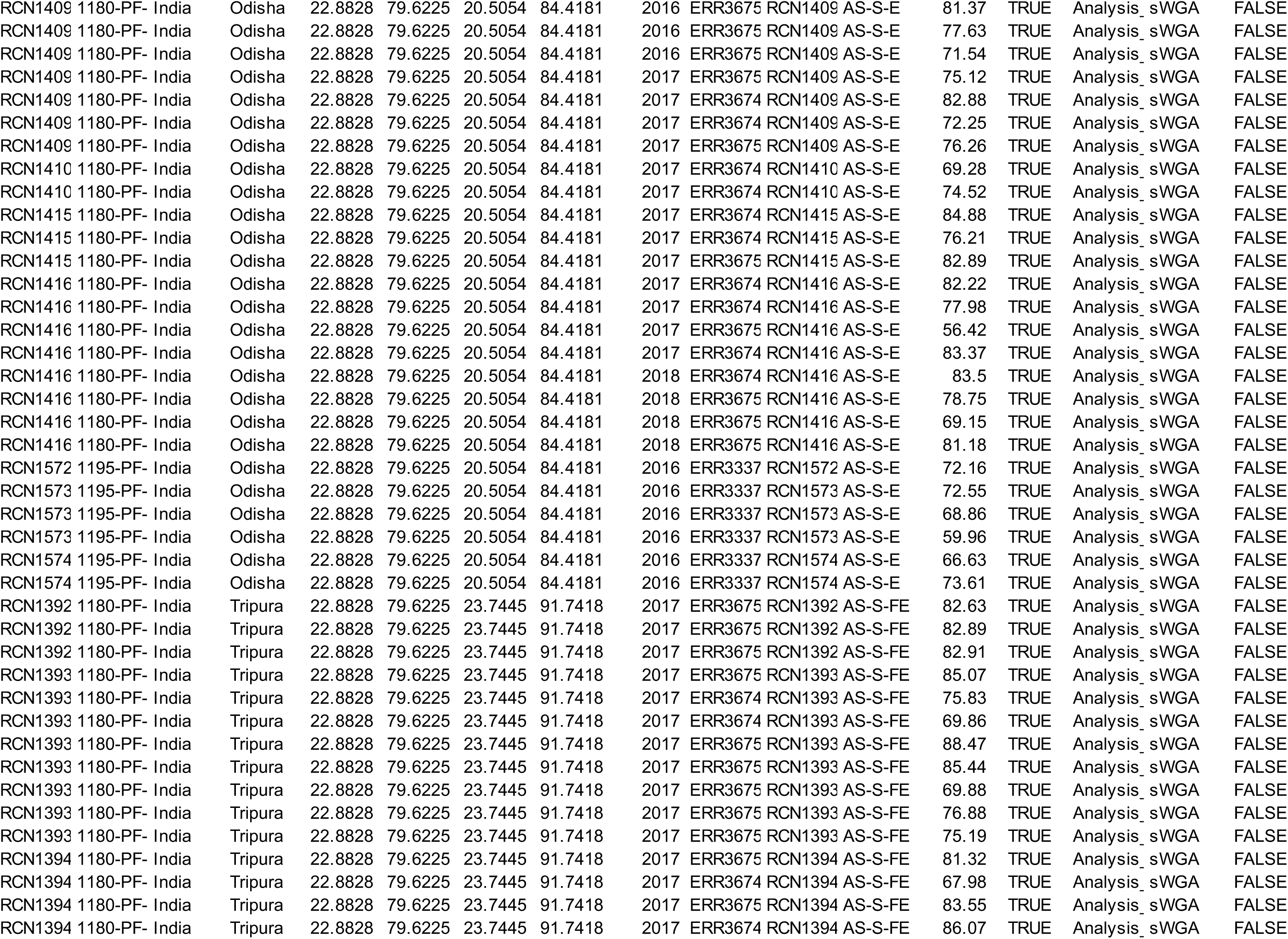
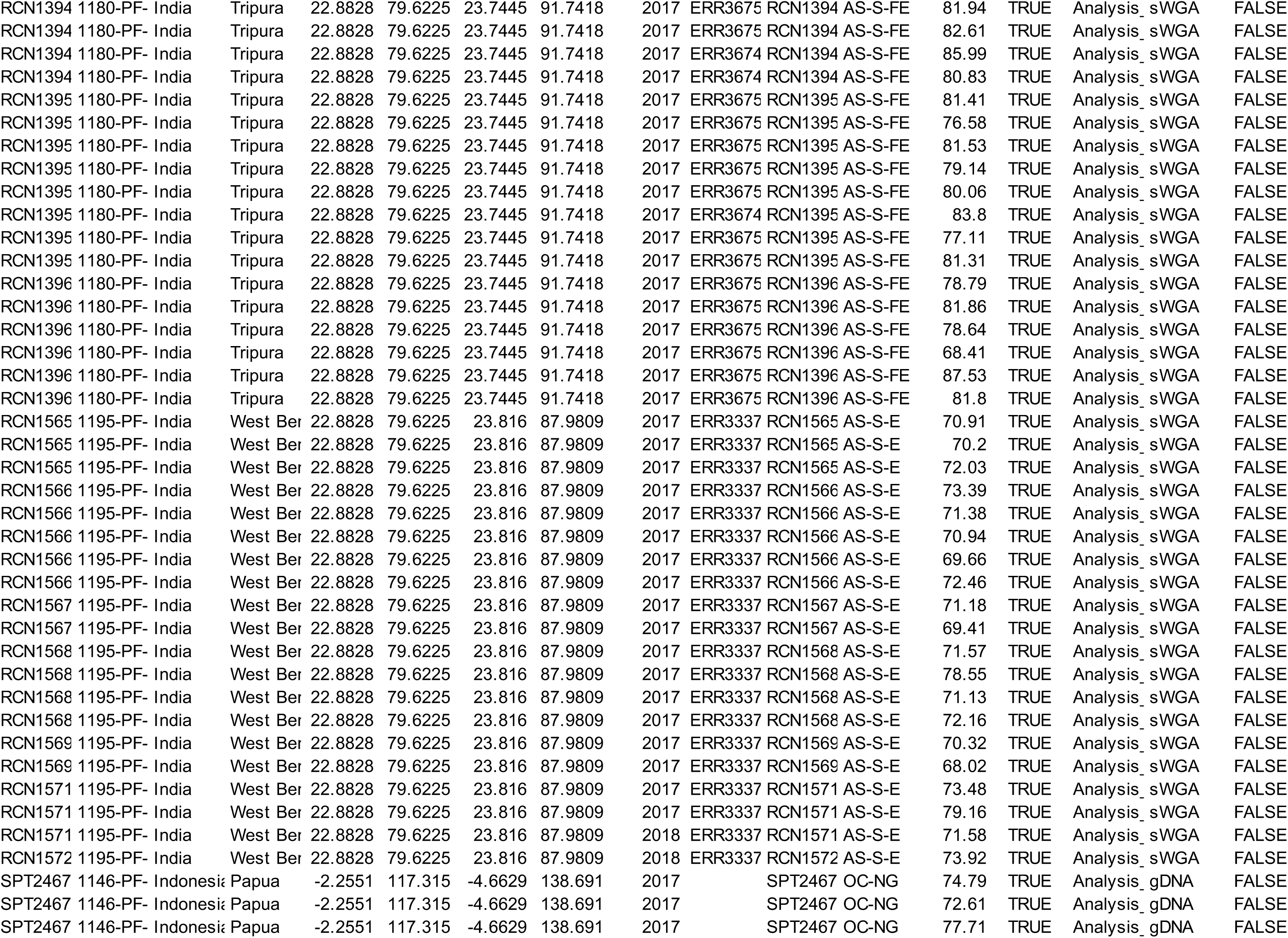
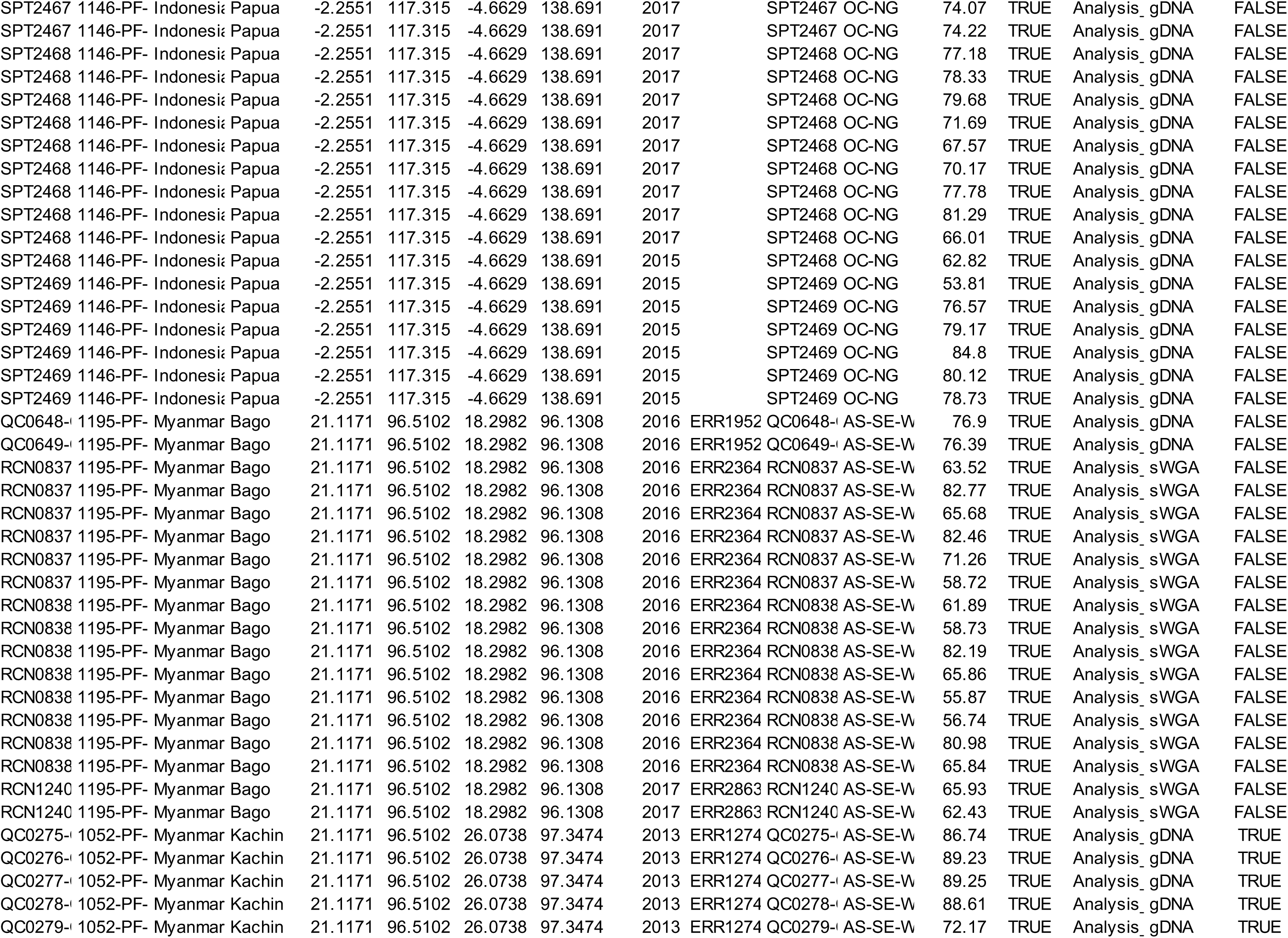
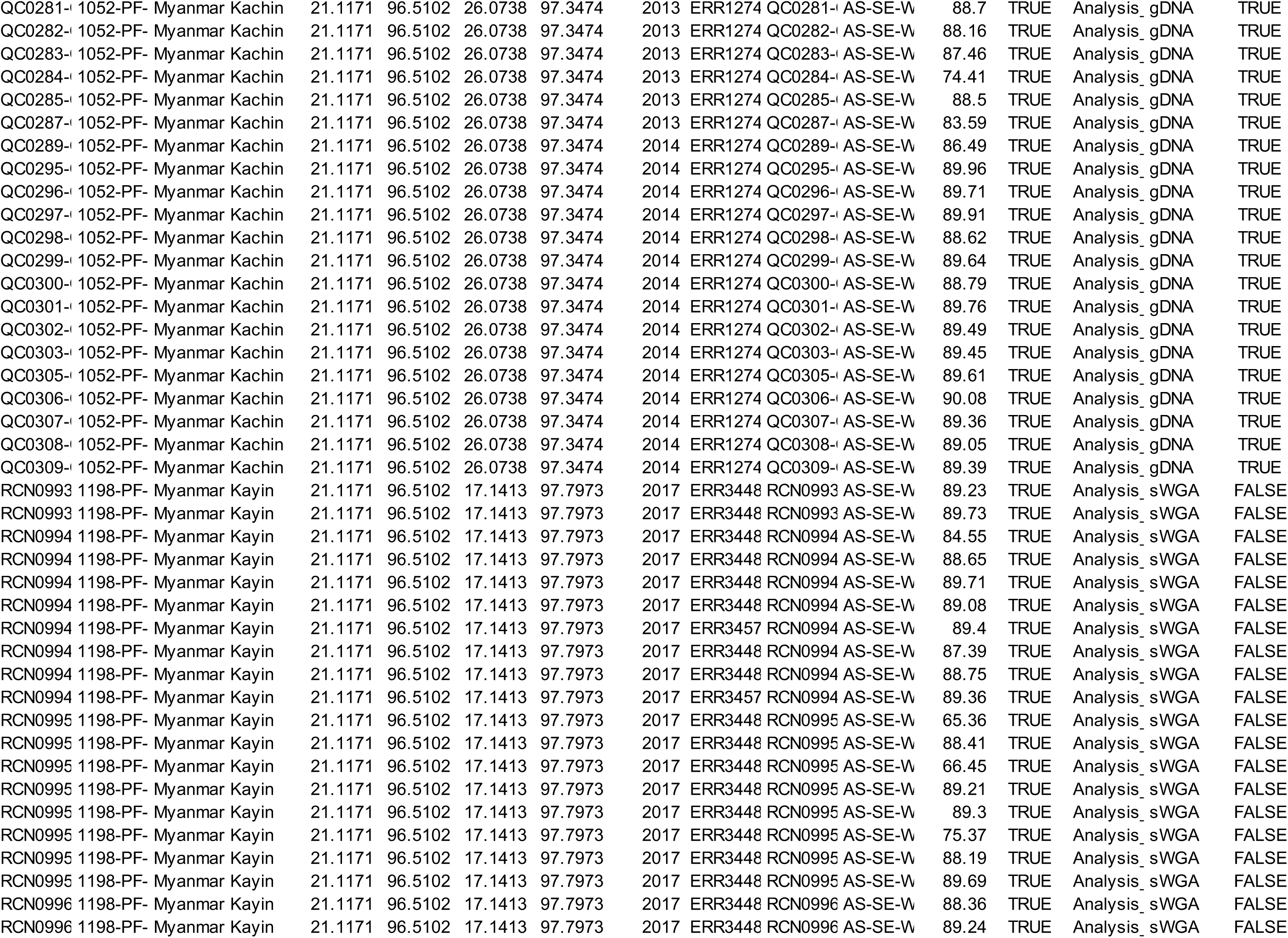
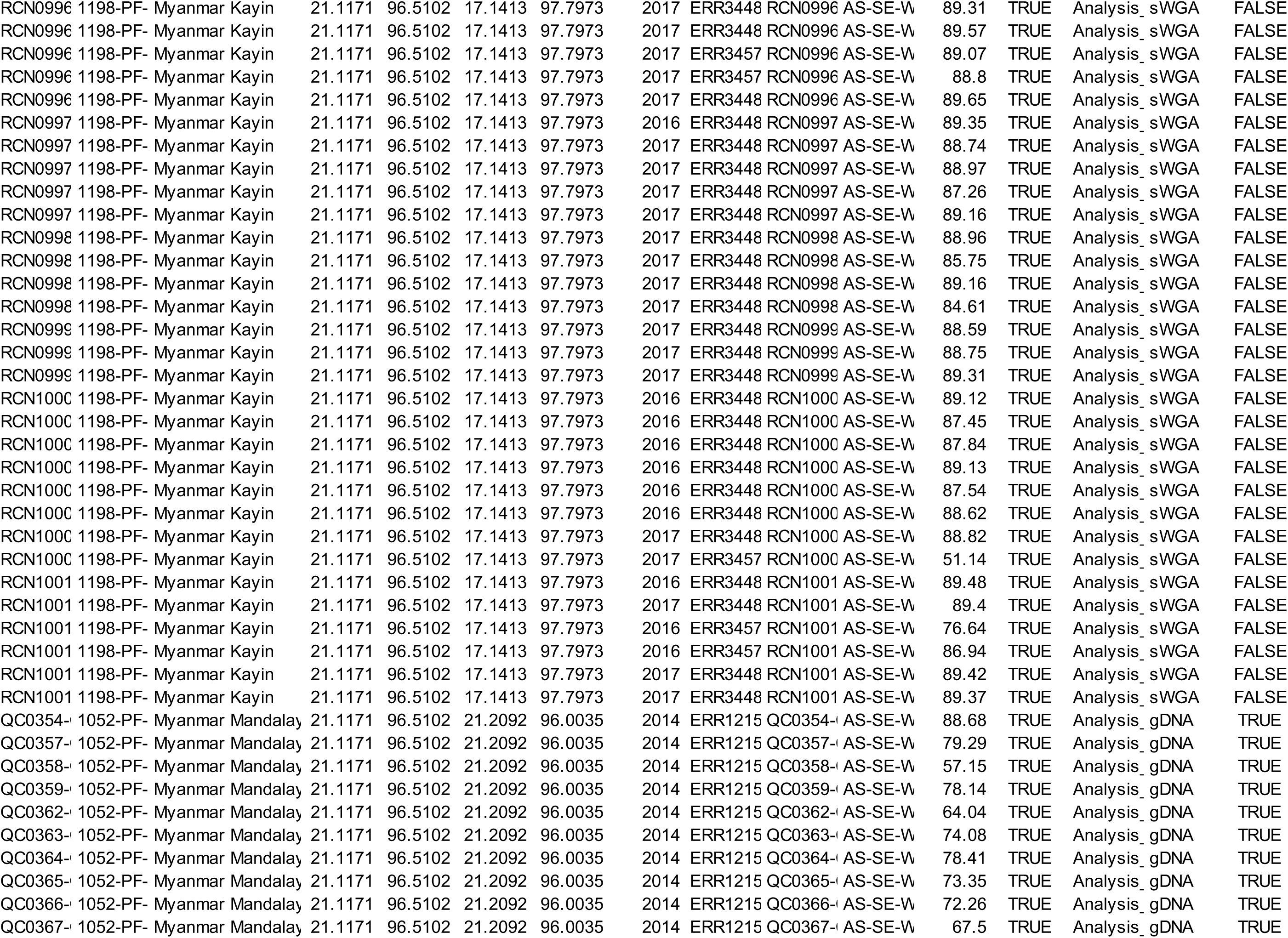
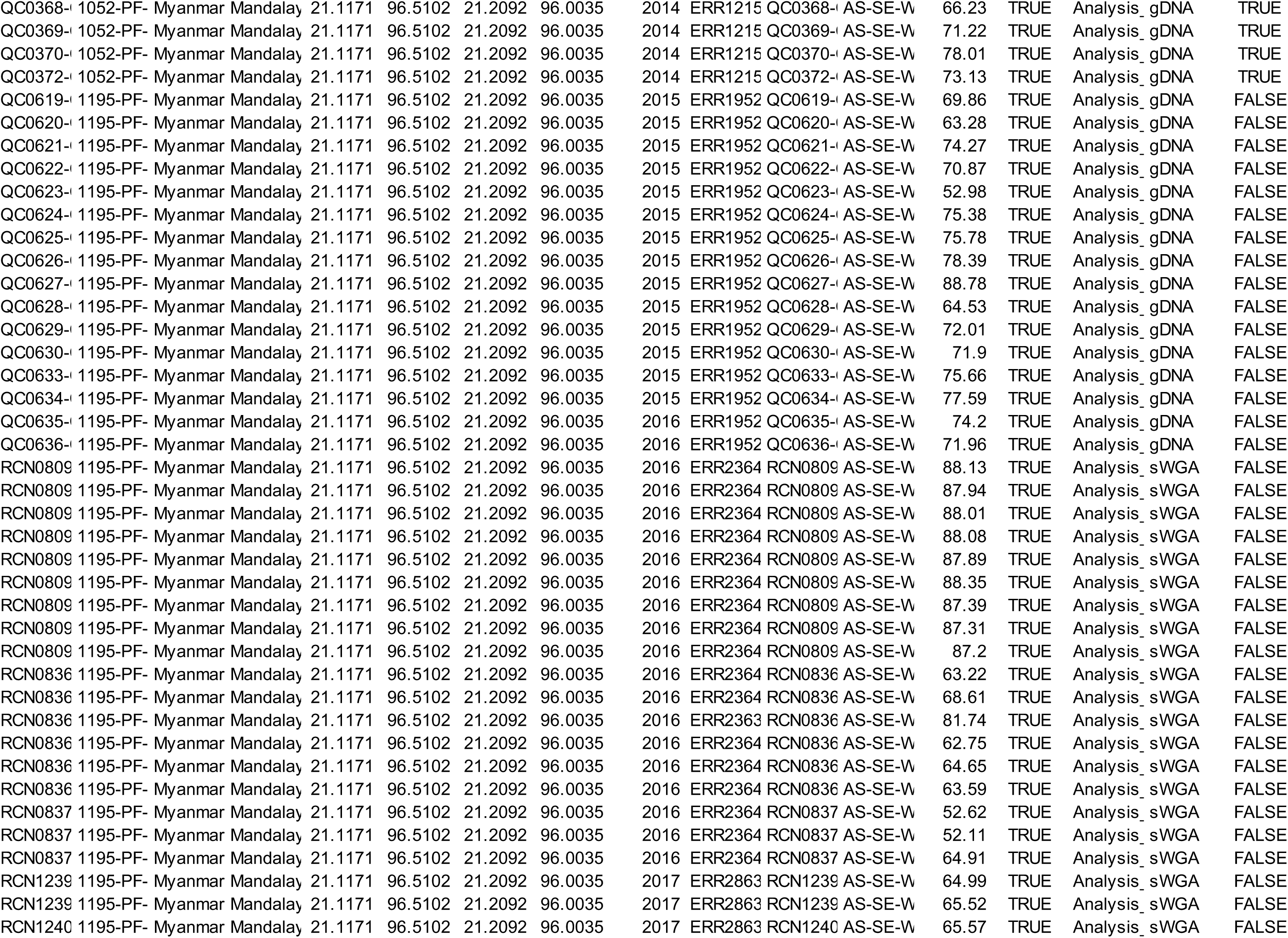
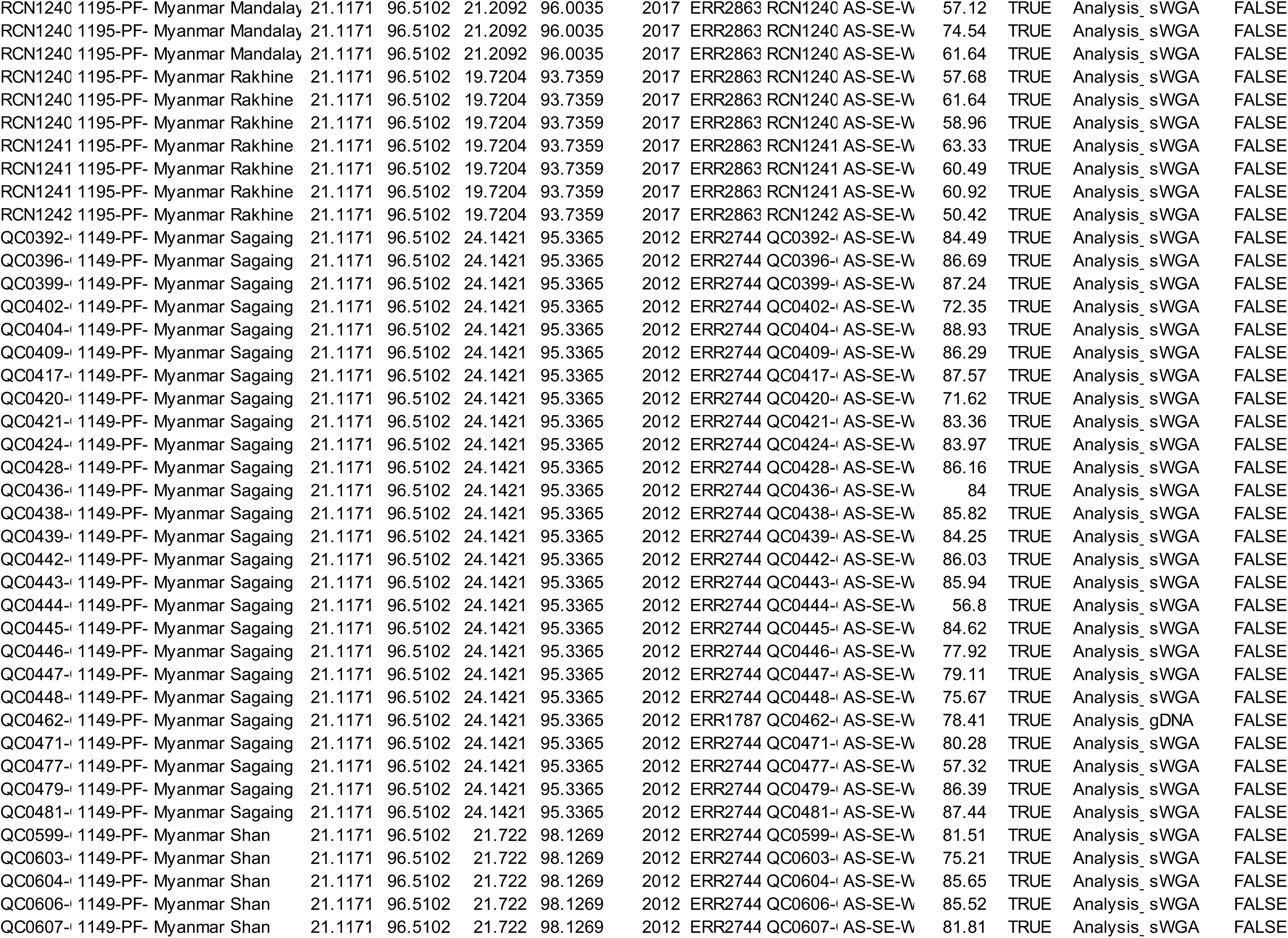
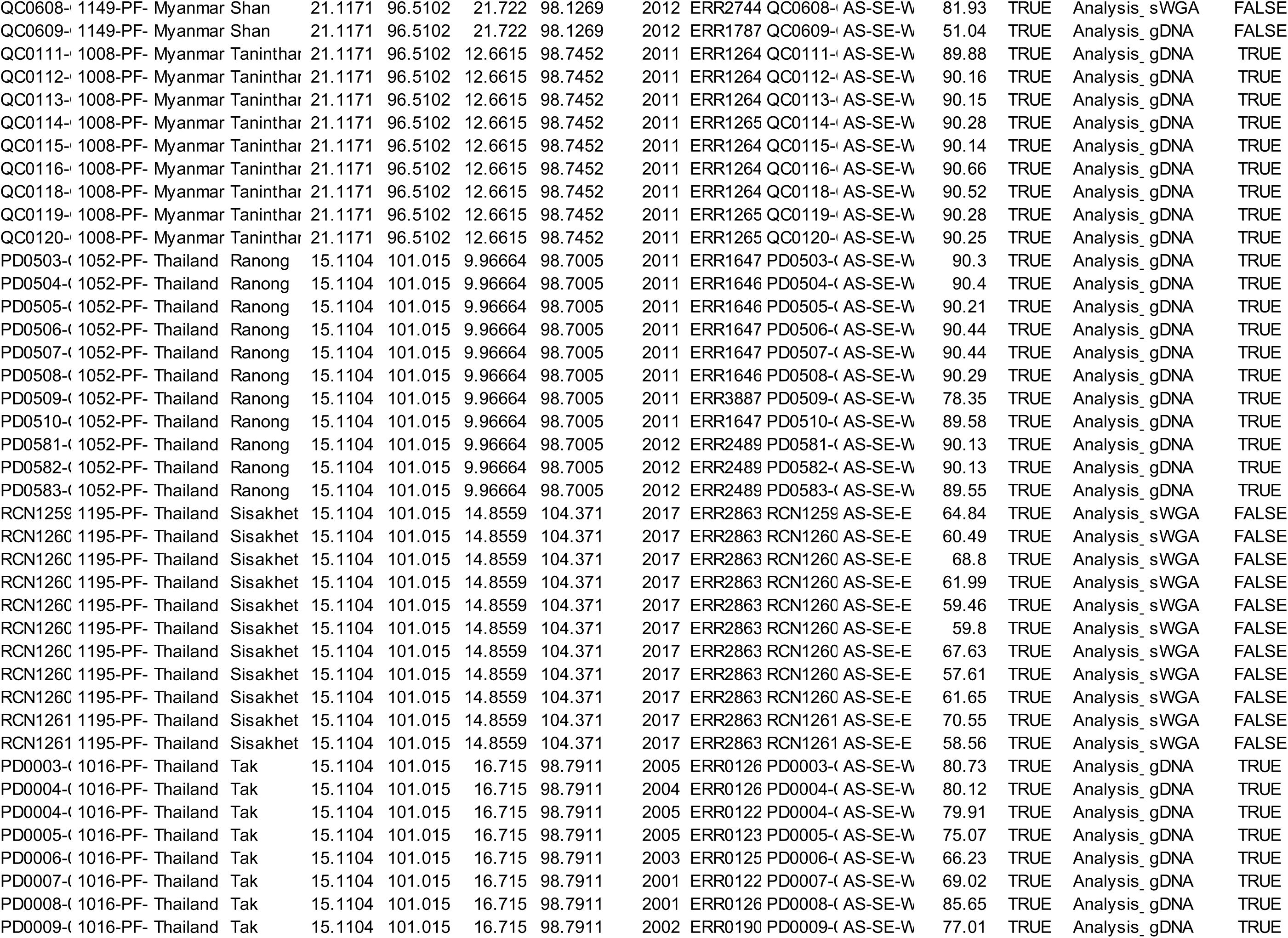
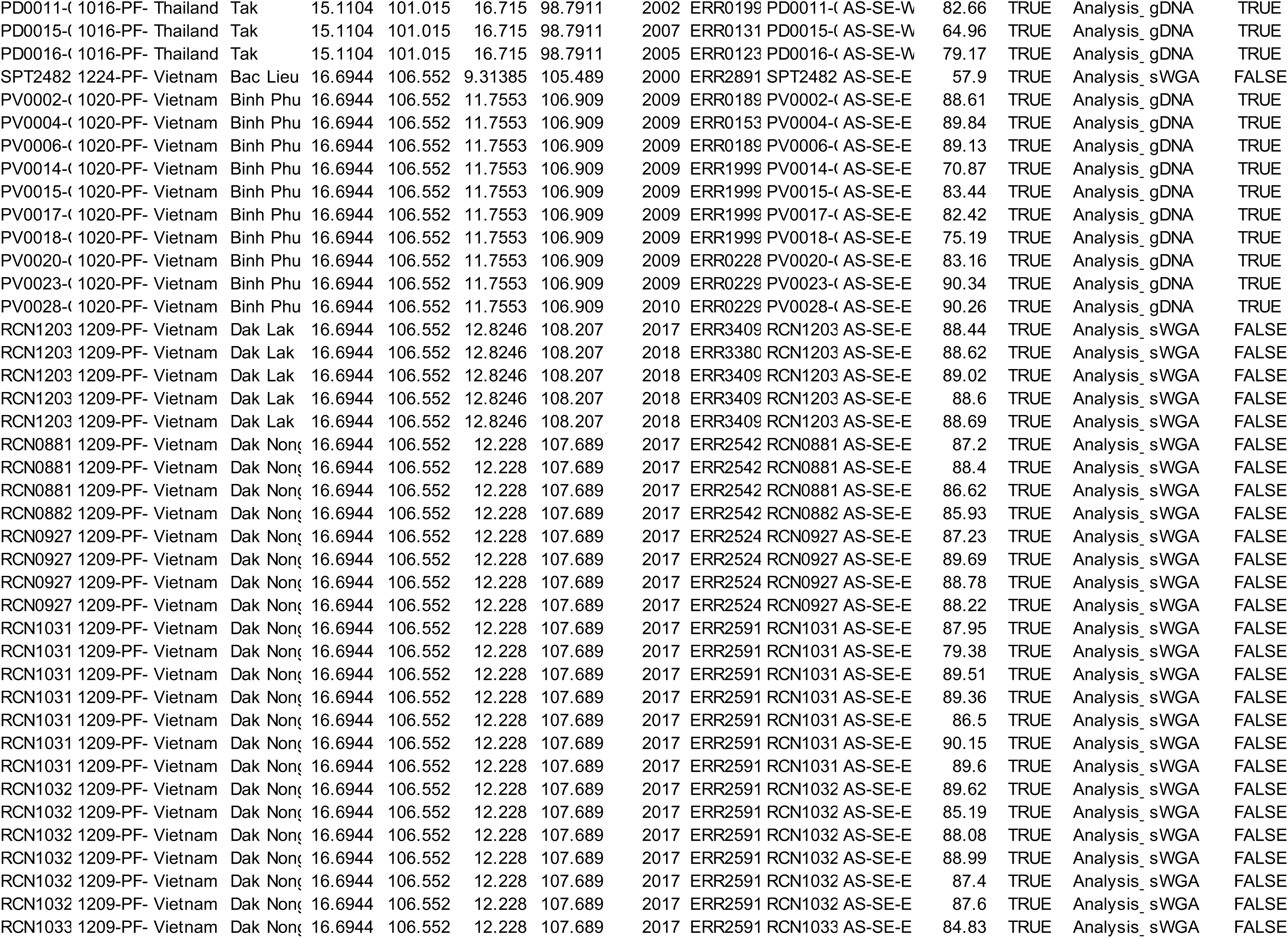
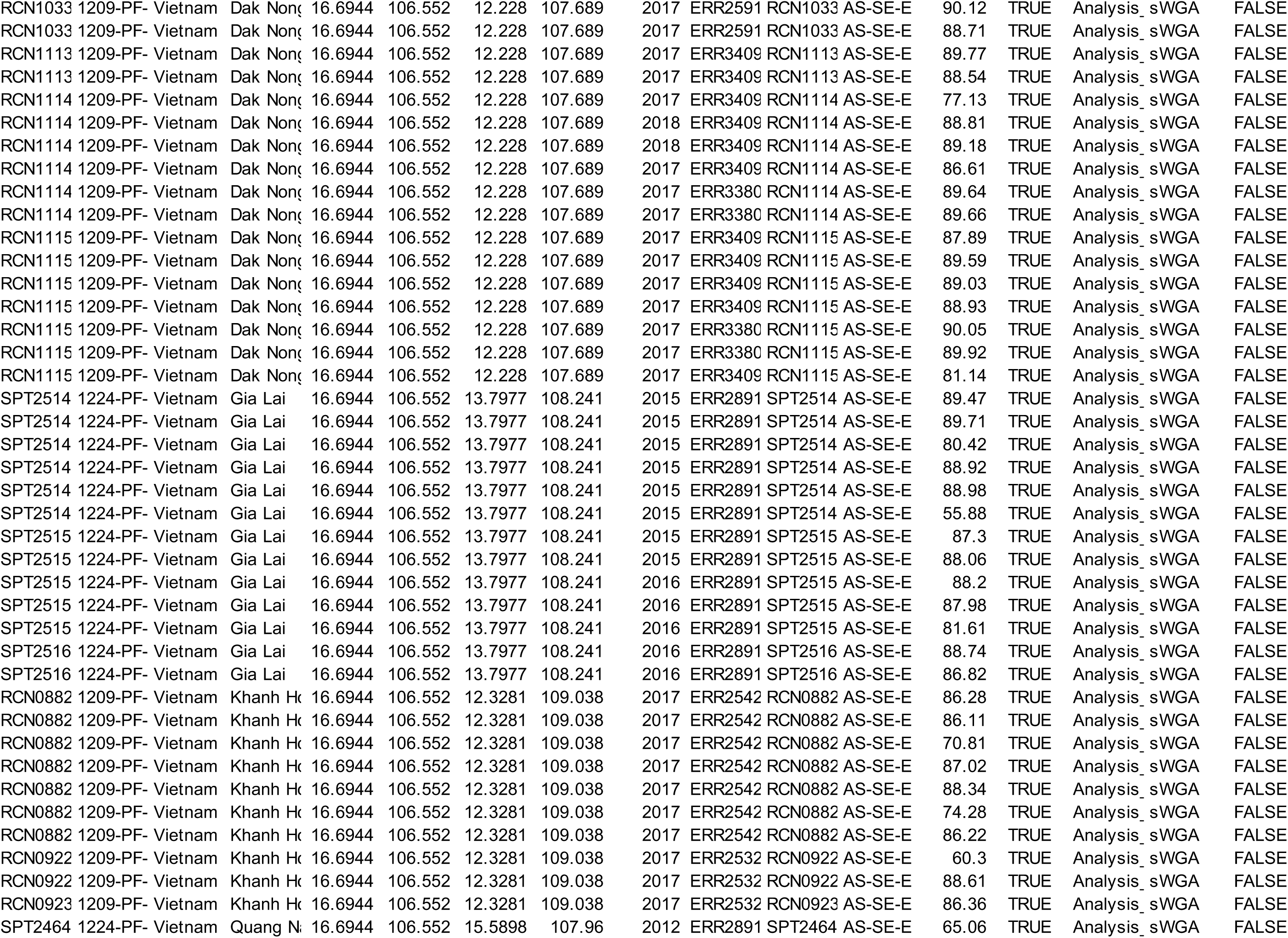
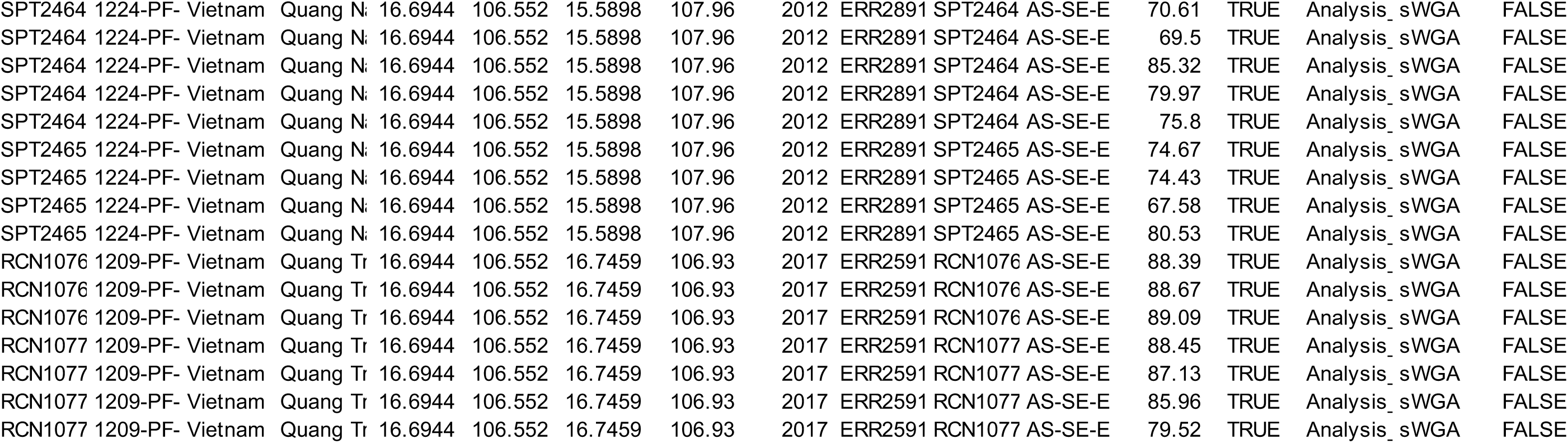
Metadata from Pf7 MalariaGEN for the PCA analysis.

## Notes

https://github.com/NirjharBhattacharyya/PCA_for_CRISPR_Paper-

### Competing Interest Statement

The authors have declared no competing interest.

## References

Adjalley, Sophie, and Marcus Chee San Lee. 2022. “CRISPR/Cas9 Editing of the Plasmodium Falciparum Genome.” In , 221–39.

Alam, Mohammad Shafiul, Benedikt Ley, Maisha Khair Nima, Fatema Tuj Johora, Mohammad Enayet Hossain, Kamala Thriemer, Sarah Auburn, Jutta Marfurt, Ric N. Price, and Wasif A. Khan. 2017. “Molecular Analysis Demonstrates High Prevalence of Chloroquine Resistance but No Evidence of Artemisinin Resistance in Plasmodium Falciparum in the Chittagong Hill Tracts of Bangladesh.” Malaria Journal 16 (1). 10.1186/s12936-017-1995-5.

Alker, Alisa P., Pharath Lim, Rithy Sem, Naman K. Shah, Poravuth Yi, Denis Mey Bouth, Reiko Tsuyuoka, et al. 2007. “Pfmdr1 and in Vivo Resistance to Artesunate-Mefloquine in Falciparum Malaria on the Cambodian-Thai Border.” The American Journal of Tropical Medicine and Hygiene 76 (4): 641–47.

Amambua-Ngwa, Alfred, Katrina A. Button-Simons, Xue Li, Sudhir Kumar, Katelyn Vendrely Brenneman, Marco Ferrari, Lisa A. Checkley, et al. 2023. “Chloroquine Resistance Evolution in Plasmodium Falciparum Is Mediated by the Putative Amino Acid Transporter AAT1.” Nature Microbiology 8 (7): 1213–26.

Amato, Roberto, Richard D. Pearson, Jacob Almagro-Garcia, Chanaki Amaratunga, Pharath Lim, Seila Suon, Sokunthea Sreng, et al. 2018. “Origins of the Current Outbreak of Multidrug-Resistant Malaria in Southeast Asia: A Retrospective Genetic Study.” The Lancet Infectious Diseases 18 (3). 10.1016/S1473-3099(18)30068-9.

Anderson, Tim J. C., Shalini Nair, Standwell Nkhoma, Jeff T. Williams, Mallika Imwong, Poravuth Yi, Duong Socheat, et al. 2010. “High Heritability of Malaria Parasite Clearance Rate Indicates a Genetic Basis for Artemisinin Resistance in Western Cambodia.” The Journal of Infectious Diseases 201 (9): 1326–30.

Aninagyei, Enoch, Kwabena Obeng Duedu, Tanko Rufai, Comfort Dede Tetteh, Margaretta Gloria Chandi, Paulina Ampomah, and Desmond Omane Acheampong. 2020. “Characterization of Putative Drug Resistant Biomarkers in Plasmodium Falciparum Isolated from Ghanaian Blood Donors.” BMC Infectious Diseases 20 (1): 533.

Ariey, Frédéric, Benoit Witkowski, Chanaki Amaratunga, Johann Beghain, Anne-Claire Langlois, Nimol Khim, Saorin Kim, et al. 2014. “A Molecular Marker of Artemisinin-Resistant Plasmodium Falciparum Malaria.” Nature 505 (7481): 50–55.

“Artemisinin Molecular Surveyor.” n.d. Accessed April 1, 2025. https://www.iddo.org/wwarn/tracking-resistance/artemisinin-molecular-surveyor.

Ashley, Elizabeth A., Mehul Dhorda, Rick M. Fairhurst, Chanaki Amaratunga, Parath Lim, Seila Suon, Sokunthea Sreng, et al. 2014. “Spread of Artemisinin Resistance in Plasmodium Falciparum Malaria.” The New England Journal of Medicine 371 (5). 10.1056/nejmoa1314981.

“Association of Mutations in the Plasmodium Falciparum Kelch13 Gene (Pf3D7_1343700) with Parasite Clearance Rates after Artemisinin-Based Treatments—a WWARN Individual Patient Data Meta-Analysis.” 2019. BMC Medicine 17 (1): 1.

Asua, Victor, Melissa D. Conrad, Ozkan Aydemir, Marvin Duvalsaint, Jennifer Legac, Elias Duarte, Patrick Tumwebaze, et al. 2021. “Changing Prevalence of Potential Mediators of Aminoquinoline, Antifolate, and Artemisinin Resistance across Uganda.” The Journal of Infectious Diseases 223 (6): 985–94.

Balikagala, Betty, Naoyuki Fukuda, Mie Ikeda, Osbert T. Katuro, Shin-Ichiro Tachibana, Masato Yamauchi, Walter Opio, et al. 2021. “Evidence of Artemisinin-Resistant Malaria in Africa.” The New England Journal of Medicine 385 (13): 1163–71.

Balmer, Andrew J., Nina F. D. White, Eyyüb S. Ünlü, Chiyun Lee, Richard D. Pearson, Jacob Almagro-Garcia, and Cristina V. Ariani. 2024. “Understanding the Global Spread of Artemisinin Resistance: Insights from over 100K*Plasmodium Falciparum*samples.” bioRxiv. 10.1101/2024.08.23.609367.

Bergmann, Clara, Welmoed van Loon, Felix Habarugira, Costanza Tacoli, Julia C. Jäger, Darius Savelsberg, Fabian Nshimiyimana, et al. 2021. “Increase in Kelch 13 Polymorphisms in Plasmodium Falciparum, Southern Rwanda.” Emerging Infectious Diseases 27 (1): 294–96.

Birnbaum, Jakob, Sarah Scharf, Sabine Schmidt, Ernst Jonscher, Wieteke Anna Maria Hoeijmakers, Sven Flemming, Christa Geeke Toenhake, et al. 2020. “A Kelch13-Defined Endocytosis Pathway Mediates Artemisinin Resistance in Malaria Parasites.” Science 367 (6473): 51–59.

Boullé, Mikael, Benoit Witkowski, Valentine Duru, Kanlaya Sriprawat, Shalini K. Nair, Marina McDew-White, Tim J. C. Anderson, Aung Pyae Phyo, Didier Menard, and François Nosten. 2016. “Artemisinin-Resistant Plasmodium Falciparum K13 Mutant Alleles, Thailand-Myanmar Border.” Emerging Infectious Diseases 22 (8): 1503–5.

Button-Simons, Katrina A., Sudhir Kumar, Nelly Carmago, Meseret T. Haile, Catherine Jett, Lisa A. Checkley, Spencer Y. Kennedy, et al. 2021. “The Power and Promise of Genetic Mapping from Plasmodium Falciparum Crosses Utilizing Human Liver-Chimeric Mice.” Communications Biology 4 (1): 734.

Chaisatit, Chaiyaporn, Piyaporn Sai-Ngam, Sasikanya Thaloengsok, Sabaithip Sriwichai, Krisada Jongsakul, Mark Fukuda, Michele Spring, et al. 2021. “Molecular Detection of Mutations in the Propeller Domain of Kelch 13 and pfmdr1 Copy Number Variation in Plasmodium Falciparum Isolates from Thailand Collected from 2002 to 2007.” The American Journal of Tropical Medicine and Hygiene 105 (4): 1093–96.

Cheeseman, Ian H., Becky A. Miller, Shalini Nair, Standwell Nkhoma, Asako Tan, John C. Tan, Salma Al Saai, et al. 2012. “A Major Genome Region Underlying Artemisinin Resistance in Malaria.” Science 336 (6077): 79–82.

Chenet, Stella M., Sheila Akinyi Okoth, Curtis S. Huber, Javin Chandrabose, Naomi W. Lucchi, Eldin Talundzic, Karanchand Krishnalall, et al. 2016. “Independent Emergence of the *Plasmodium Falciparum* Kelch Propeller Domain Mutant Allele C580Y in Guyana.” The Journal of Infectious Diseases 213 (9): 1472–75.

Chhibber-Goel, Jyoti, and Amit Sharma. 2019. “Profiles of Kelch Mutations in Plasmodium Falciparum across South Asia and Their Implications for Tracking Drug Resistance.” *International Journal for Parasitology*, Drugs and Drug Resistance 11 (December):49–58.

Conrad, Melissa D., Victor Asua, Shreeya Garg, David Giesbrecht, Karamoko Niaré, Sawyer Smith, Jane F. Namuganga, et al. 2023. “Evolution of Partial Resistance to Artemisinins in Malaria Parasites in Uganda.” The New England Journal of Medicine 389 (8): 722–32.

Demas, Allison R., Aabha I. Sharma, Wesley Wong, Angela M. Early, Seth Redmond, Selina Bopp, Daniel E. Neafsey, Sarah K. Volkman, Daniel L. Hartl, and Dyann F. Wirth. 2018. “Mutations in *Plasmodium Falciparum* Actin-Binding Protein Coronin Confer Reduced Artemisinin Susceptibility.” Proceedings of the National Academy of Sciences 115 (50): 12799–804.

Denis, Mey Bouth, Reiko Tsuyuoka, Pharath Lim, Niklas Lindegardh, Poravuth Yi, Sophoan Narann Top, Duong Socheat, et al. 2006. “Efficacy of Artemether?lumefantrine for the Treatment of Uncomplicated Falciparum Malaria in Northwest Cambodia.” Tropical Medicine & International Health: TM & IH 11 (12): 1800–1807.

Dondorp, Arjen M., François Nosten, Poravuth Yi, Debashish Das, Aung Phae Phyo, Joel Tarning, Khin Maung Lwin, et al. 2009. “Artemisinin Resistance in Plasmodium Falciparum Malaria.” The New England Journal of Medicine 361 (5). 10.1056/nejmoa0808859.

Dwivedi, Ankit, Nimol Khim, Christelle Reynes, Patrice Ravel, Laurence Ma, Magali Tichit, Christiane Bourchier, et al. 2016. “Plasmodium Falciparum Parasite Population Structure and Gene Flow Associated to Anti-Malarial Drugs Resistance in Cambodia.” Malaria Journal 15 (June):319.

Fola, Abebe A., G. L. Abby Harrison, Mita Hapsari Hazairin, Céline Barnadas, Manuel W. Hetzel, Jonah Iga, Peter M. Siba, Ivo Mueller, and Alyssa E. Barry. 2017. “Higher Complexity of Infection and Genetic Diversity of *Plasmodium Vivax* Than *Plasmodium Falciparum* across All Malaria Transmission Zones of Papua New Guinea.” *The American Journal of Tropical Medicine and Hygiene*, January, 16–0716.

Fola, Abebe A., Sindew M. Feleke, Hussein Mohammed, Bokretsion G. Brhane, Christopher M. Hennelly, Ashenafi Assefa, Rebecca M. Crudal, et al. 2023. “Plasmodium Falciparum Resistant to Artemisinin and Diagnostics Have Emerged in Ethiopia.” Nature Microbiology 8 (10): 1911–19.

Ghorbal, Mehdi, Molly Gorman, Cameron Ross Macpherson, Rafael Miyazawa Martins, Artur Scherf, and Jose-Juan Lopez-Rubio. 2014. “Genome Editing in the Human Malaria Parasite Plasmodium Falciparum Using the CRISPR-Cas9 System.” Nature Biotechnology 32 (8): 819–21.

Global Technical Strategy for Malaria 2016-2030, 2021 Update. 2021. World Health Organization.

Haldar, Kasturi, Mohammed Shafiul Alam, Cristian Koepfli, Neil F. Lobo, Ching Shwe Phru, Muhammad Nazmul Islam, Abul Faiz, Wasif Ali Khan, and Rashidul Haque. 2023. “Bangladesh in the Era of Malaria Elimination.” Trends in Parasitology 39 (9): 760–73.

Hamilton, William L., Roberto Amato, Rob W. van der Pluijm, Christopher G. Jacob, Huynh Hong Quang, Nguyen Thanh Thuy-Nhien, Tran Tinh Hien, et al. 2019. “Evolution and Expansion of Multidrug-Resistant Malaria in Southeast Asia: A Genomic Epidemiology Study.” The Lancet Infectious Diseases 19 (9): 943–51.

Haque, Ubydul, Hans J. Overgaard, Archie C. A. Clements, Douglas E. Norris, Nazrul Islam, Jahirul Karim, Shyamal Roy, et al. 2014. “Malaria Burden and Control in Bangladesh and Prospects for Elimination: An Epidemiological and Economic Assessment.” The Lancet Global Health 2 (2): e98–105.

He, Yan, Susana Campino, Ernest Diez Benavente, David C. Warhurst, Khalid B. Beshir, Inke Lubis, Ana Rita Gomes, et al. 2019. “Artemisinin Resistance-Associated Markers in Plasmodium Falciparum Parasites from the China-Myanmar Border: Predicted Structural Stability of K13 Propeller Variants Detected in a Low-Prevalence Area.” PloS One 14 (3): e0213686.

Huang, Fang, Shannon Takala-Harrison, Christopher G. Jacob, Hui Liu, Xiaodong Sun, Henglin Yang, Myaing M. Nyunt, et al. 2015. “A Single Mutation in K13 Predominates in Southern China and Is Associated with Delayed Clearance of Plasmodium Falciparum Following Artemisinin Treatment.” The Journal of Infectious Diseases 212 (10): 1629–35.

Huwe, Tiffany, Mohammad Golam Kibria, Fatema Tuj Johora, Ching Swe Phru, Nusrat Jahan, Mohammad Sharif Hossain, Wasif Ali Khan, et al. 2022. “Heterogeneity in Prevalence of Subclinical Plasmodium Falciparum and Plasmodium Vivax Infections but No Parasite Genomic Clustering in the Chittagong Hill Tracts, Bangladesh.” Malaria Journal 21 (1): 218.

Imwong, Mallika, Mehul Dhorda, Kyaw Myo Tun, Aung Myint Thu, Aung Pyae Phyo, Stephane Proux, Kanokon Suwannasin, et al. 2020. “Molecular Epidemiology of Resistance to Antimalarial Drugs in the Greater Mekong Subregion: An Observational Study.” The Lancet Infectious Diseases 20 (12): 1470–80.

Imwong, Mallika, Kanokon Suwannasin, Chanon Kunasol, Kreepol Sutawong, Mayfong Mayxay, Huy Rekol, Frank M. Smithuis, et al. 2017. “The Spread of Artemisinin-Resistant Plasmodium Falciparum in the Greater Mekong Subregion: A Molecular Epidemiology Observational Study.” The Lancet Infectious Diseases 17 (5): 491–97.

Ishengoma, Deus S., Celine I. Mandara, Catherine Bakari, Abebe A. Fola, Rashid A. Madebe, Misago D. Seth, Filbert Francis, et al. 2024. “Evidence of Artemisinin Partial Resistance in Northwestern Tanzania: Clinical and Molecular Markers of Resistance.” The Lancet Infectious Diseases 24 (11): 1225–33.

Juliano, Jonathan J., David J. Giesbrecht, Alfred Simkin, Abebe A. Fola, Beatus M. Lyimo, Dativa Pereus, Catherine Bakari, et al. 2023. “Country Wide Surveillance Reveals Prevalent Artemisinin Partial Resistance Mutations with Evidence for Multiple Origins and Expansion of High Level Sulfadoxine-Pyrimethamine Resistance Mutations in Northwest Tanzania.” *medRxiv : The Preprint Server for Health Sciences*, November. 10.1101/2023.11.07.23298207.

Kagoro, Frank M., Karen I. Barnes, Kevin Marsh, Nattwut Ekapirat, Chris Erwin G. Mercado, Ipsita Sinha, Georgina Humphreys, Mehul Dhorda, Philippe J. Guerin, and Richard J. Maude. 2022. “Mapping Genetic Markers of Artemisinin Resistance in Plasmodium Falciparum Malaria in Asia: A Systematic Review and Spatiotemporal Analysis.” The Lancet Microbe 3 (3). 10.1016/S2666-5247(21)00249-4.

Kateera, Fredrick, Sam L. Nsobya, Stephen Tukwasibwe, Petra F. Mens, Emmanuel Hakizimana, Martin P. Grobusch, Leon Mutesa, Nirbhay Kumar, and Michele van Vugt. 2016. “Malaria Case Clinical Profiles and Plasmodium Falciparum Parasite Genetic Diversity: A Cross Sectional Survey at Two Sites of Different Malaria Transmission Intensities in Rwanda.” Malaria Journal 15 (1): 237.

Kumar, Sudhir, Xue Li, Marina McDew-White, Ann Reyes, Elizabeth Delgado, Abeer Sayeed, Meseret T. Haile, et al. 2022. “A Malaria Parasite Cross Reveals Genetic Determinants of Plasmodium Falciparum Growth in Different Culture Media.” Frontiers in Cellular and Infection Microbiology 12 (May). 10.3389/fcimb.2022.878496.

Lalmalsawma, Pachuau, K. Balasubramani, Meenu Mariya James, Lalfakzuala Pautu, Kumar Arun Prasad, Devojit Kumar Sarma, and Praveen Balabaskaran Nina. 2023. “Malaria Hotspots and Climate Change Trends in the Hyper-Endemic Malaria Settings of Mizoram along the India-Bangladesh Borders.” Scientific Reports 13 (1): 4538.

L’Episcopia, Mariangela, Julia Kelley, Dhruviben Patel, Sarah Schmedes, Shashidahar Ravishankar, Michela Menegon, Edvige Perrotti, et al. 2020. “Targeted Deep Amplicon Sequencing of Kelch 13 and Cytochrome B in Plasmodium Falciparum Isolates from an Endemic African Country Using the Malaria Resistance Surveillance (MaRS) Protocol.” Parasites & Vectors 13 (1): 137.

Loon, Welmoed van, Rafael Oliveira, Clara Bergmann, Felix Habarugira, Jules Ndoli, Augustin Sendegeya, Claude Bayingana, and Frank P. Mockenhaupt. 2022. “In Vitro Confirmation of Artemisinin Resistance in Plasmodium Falciparum from Patient Isolates, Southern Rwanda, 2019.” Emerging Infectious Diseases 28 (4): 852–55.

Lu, Feng, Richard Culleton, Meihua Zhang, Abhinay Ramaprasad, Lorenz von Seidlein, Huayun Zhou, Guoding Zhu, et al. 2017. “Emergence of Indigenous Artemisinin-Resistant *Plasmodium Falciparum* in Africa.” The New England Journal of Medicine 376 (10): 991– 93.

MalariaGEN, Muzamil Mahdi Abdel Hamid, Mohamed Hassan Abdelraheem, Desmond Omane Acheampong, Ambroise Ahouidi, Mozam Ali, Jacob Almagro-Garcia, et al. 2023. “Pf7: An Open Dataset of Plasmodium Falciparum Genome Variation in 20,000 Worldwide Samples.” Wellcome Open Research 8 (January):22.

Mathieu, Luana C., Horace Cox, Angela M. Early, Sachel Mok, Yassamine Lazrek, Jeanne-Celeste Paquet, Maria-Paz Ade, et al. 2020. “Local Emergence in Amazonia of Plasmodium Falciparum k13 C580Y Mutants Associated with in Vitro Artemisinin Resistance.” eLife 9 (May). 10.7554/eLife.51015.

Matrevi, Sena A., Philip Opoku-Agyeman, Neils B. Quashie, Selassie Bruku, Benjamin Abuaku, Kwadwo A. Koram, Anne Fox, Andrew Letizia, and Nancy O. Duah-Quashie. 2019. “Plasmodium Falciparum Kelch Propeller Polymorphisms in Clinical Isolates from Ghana from 2007 to 2016.” Antimicrobial Agents and Chemotherapy 63 (11). 10.1128/AAC.00802-19.

Maude, Richard James, Mahtab Uddin Hasan, Md Amir Hossain, Abdullah Abu Sayeed, Sanjib Kanti Paul, Waliur Rahman, Rapeephan Rattanawongnara Maude, et al. 2012. “Temporal Trends in Severe Malaria in Chittagong, Bangladesh.” Malaria Journal 11 (1): 323.

Mbengue, Alassane, Souvik Bhattacharjee, Trupti Pandharkar, Haining Liu, Guillermina Estiu, Robert V. Stahelin, Shahir S. Rizk, et al. 2015. “A Molecular Mechanism of Artemisinin Resistance in Plasmodium Falciparum Malaria.” Nature 520 (7549): 683–87.

Ménard, Didier, Nimol Khim, Johann Beghain, Ayola A. Adegnika, Mohammad Shafiul-Alam, Olukemi Amodu, Ghulam Rahim-Awab, et al. 2016. “A Worldwide Map of Plasmodium Falciparum K13-Propeller Polymorphisms.” The New England Journal of Medicine 374 (25). 10.1056/nejmoa1513137.

Mihreteab, Selam, Lucien Platon, Araia Berhane, Barbara H. Stokes, Marian Warsame, Pascal Campagne, Alexis Criscuolo, et al. 2023. “Increasing Prevalence of Artemisinin-Resistant HRP2-Negative Malaria in Eritrea.” The New England Journal of Medicine 389 (13): 1191– 1202.

Miotto, Olivo, Roberto Amato, Elizabeth A. Ashley, Bronwyn Macinnis, Jacob Almagro-Garcia, Chanaki Amaratunga, Pharath Lim, et al. 2015. “Genetic Architecture of Artemisinin-Resistant Plasmodium Falciparum.” Nature Genetics 47 (3). 10.1038/ng.3189.

Miotto, Olivo, Makoto Sekihara, Shin-Ichiro Tachibana, Masato Yamauchi, Richard D. Pearson, Roberto Amato, Sonia Gonçalves, et al. 2020. “Emergence of Artemisinin-Resistant Plasmodium Falciparum with kelch13 C580Y Mutations on the Island of New Guinea.” PLoS Pathogens 16 (12): e1009133.

Mishra, Neelima, Surendra Kumar Prajapati, Kamlesh Kaitholia, Ram Suresh Bharti, Bina Srivastava, Sobhan Phookan, Anupkumar R. Anvikar, et al. 2015. “Surveillance of Artemisinin Resistance in Plasmodium Falciparum in India Using the kelch13 Molecular Marker.” Antimicrobial Agents and Chemotherapy 59 (5): 2548–53.

Mita, Toshihiro, Kazuyuki Tanabe, and Kiyoshi Kita. 2009. “Spread and Evolution of Plasmodium Falciparum Drug Resistance.” Parasitology International 58 (3): 201–9.

Mohon, Abu Naser, Mohammad Shafiul Alam, Abebe Genetu Bayih, Asongna Folefoc, Dea Shahinas, Rashidul Haque, and Dylan R. Pillai. 2014. “Mutations in Plasmodium Falciparum K13 Propeller Gene from Bangladesh (2009-2013).” Malaria Journal 13 (1). 10.1186/1475-2875-13-431.

“More on Artemisinin-Resistant *Plasmodium Falciparum* in Eastern India.” 2019. The New England Journal of Medicine 380 (11): e14.

Mukherjee, Angana, Selina Bopp, Pamela Magistrado, Wesley Wong, Rachel Daniels, Allison Demas, Stephen Schaffner, et al. 2017. “Artemisinin Resistance without pfkelch13 Mutations in Plasmodium Falciparum Isolates from Cambodia.” Malaria Journal 16 (1). 10.1186/s12936-017-1845-5.

Nair, Shalini, Xue Li, Grace A. Arya, Marina McDew-White, Marco Ferrari, François Nosten, and Tim J. C. Anderson. 2018. “Fitness Costs and the Rapid Spread of kelch13 -C580Y Substitutions Conferring Artemisinin Resistance.” Antimicrobial Agents and Chemotherapy 62 (9): AAC.00605–18.

“National Strategic Plan_Malaria Elimination_Bangladesh_2021-2025.pdf.” n.d.

Ndwiga, Leonard, Kelvin M. Kimenyi, Kevin Wamae, Victor Osoti, Mercy Akinyi, Irene Omedo, Deus S. Ishengoma, et al. 2021. “A Review of the Frequencies of Plasmodium Falciparum Kelch 13 Artemisinin Resistance Mutations in Africa.” *International Journal for Parasitology*, Drugs and Drug Resistance 16 (August):155–61.

Nima, Maisha Khair, Angana Mukherjee, Saiful Arefeen Sazed, Muhammad Riadul Haque Hossainey, Ching Swe Phru, Fatema Tuj Johora, Innocent Safeukui, et al. 2022. “Assessment of Plasmodium Falciparum Artemisinin Resistance Independent of kelch13 Polymorphisms and with Escalating Malaria in Bangladesh.” mBio 13 (1). 10.1128/MBIO.03444-21.

Nkhoma, Standwell C., Shalini Nair, Salma Al-Saai, Elizabeth Ashley, Rose McGready, Aung P. Phyo, François Nosten, and Tim J. C. Anderson. 2013. “Population Genetic Correlates of Declining Transmission in a Human Pathogen.” Molecular Ecology 22 (2): 273–85.

Noé, Andrés, Sazid Ibna Zaman, Mosiqure Rahman, Anjan Kumar Saha, M. M. Aktaruzzaman, and Richard James Maude. 2018. “Mapping the Stability of Malaria Hotspots in Bangladesh from 2013 to 2016.” Malaria Journal 17 (1). 10.1186/s12936-018-2405-3.

Noedl, Harald, Youry Se, Kurt Schaecher, Bryan L. Smith, Duong Socheat, and Mark M. Fukuda. 2008. “Evidence of Artemisinin-Resistant Malaria in Western Cambodia.” The New England Journal of Medicine 359 (24). 10.1056/nejmc0805011.

Nyunt, Myat Htut, Thaung Hlaing, Htet Wai Oo, Lu Lu Kyaw Tin-Oo, Hnin Phyu Phway, Bo Wang, Ni Ni Zaw, et al. 2015. “Molecular Assessment of Artemisinin Resistance Markers, Polymorphisms in the K13 Propeller, and a Multidrug-Resistance Gene in the Eastern and Western Border Areas of Myanmar.” Clinical Infectious Diseases: An Official Publication of the Infectious Diseases Society of America 60 (8). 10.1093/cid/ciu1160.

O’Neill, Paul M., Richard K. Amewu, Susan A. Charman, Sunil Sabbani, Nina F. Gnädig, Judith Straimer, David A. Fidock, et al. 2017. “A Tetraoxane-Based Antimalarial Drug Candidate That Overcomes PfK13-C580Y Dependent Artemisinin Resistance.” Nature Communications 8 (May):15159.

Pillai, Dylan R., Abebe Alemu, Sisay Getie, Abebe Genetu Bayih, Abu Naser Mohon, and Gebeyaw Getnet. 2016. “A Unique Plasmodium Falciparum K13 Gene Mutation in Northwest Ethiopia.” The American Journal of Tropical Medicine and Hygiene 94 (1): 132– 35.

Pluijm, Rob W. van der, Mallika Imwong, Nguyen Hoang Chau, Nhu Thi Hoa, Nguyen Thanh Thuy-Nhien, Ngo Viet Thanh, Podjanee Jittamala, et al. 2019. “Determinants of Dihydroartemisinin-Piperaquine Treatment Failure in Plasmodium Falciparum Malaria in Cambodia, Thailand, and Vietnam: A Prospective Clinical, Pharmacological, and Genetic Study.” The Lancet Infectious Diseases 19 (9): 952–61.

Rocamora, Frances, Lei Zhu, Kek Yee Liong, Arjen Dondorp, Olivo Miotto, Sachel Mok, and Zbynek Bozdech. 2018. “Oxidative Stress and Protein Damage Responses Mediate Artemisinin Resistance in Malaria Parasites.” PLoS Pathogens 14 (3): e1006930.

Shannon, Kerry L., Wasif A. Khan, David A. Sack, M. Shafiul Alam, Sabeena Ahmed, Chai Shwai Prue, Jacob Khyang, et al. 2016. “Subclinical Plasmodium Falciparum Infections Act as Year-Round Reservoir for Malaria in the Hypoendemic Chittagong Hill Districts of Bangladesh.” International Journal of Infectious Diseases: IJID: Official Publication of the International Society for Infectious Diseases 49 (August):161–69.

Siddiqui, Faiza Amber, Rachasak Boonhok, Mynthia Cabrera, Huguette Gaelle Ngassa Mbenda, Meilian Wang, Hui Min, Xiaoying Liang, et al. 2020. “Role of Plasmodium Falciparum Kelch 13 Protein Mutations in P. Falciparum Populations from Northeastern Myanmar in Mediating Artemisinin Resistance.” mBio 11 (1). 10.1128/mBio.01134-19.

Sinha, Ipsita, Abdullah Abu Sayeed, Didar Uddin, Amy Wesolowski, Sazid Ibna Zaman, M. Abul Faiz, Aniruddha Ghose, et al. 2020. “Mapping the Travel Patterns of People with Malaria in Bangladesh.” BMC Medicine 18 (1). 10.1186/s12916-020-1512-5.

Srisutham, Suttipat, Aungkana Saejeng, Nardlada Khantikul, Rungniran Sugaram, Raweewan Sangsri, Arjen M. Dondorp, Nicholas P. J. Day, and Mallika Imwong. 2025. “Advancing Artemisinin Resistance Monitoring Using a High Sensitivity ddPCR Assay for Pfkelch13 Mutation Detection in Asia.” Scientific Reports 15 (1): 4869.

Stokes, Barbara H., Satish K. Dhingra, Kelly Rubiano, Sachel Mok, Judith Straimer, Nina F. Gnädig, Ioanna Deni, et al. 2021. “Plasmodium Falciparum k13 Mutations in Africa and Asia Impact Artemisinin Resistance and Parasite Fitness.” eLife 10. 10.7554/eLife.66277.

Straimer, Judith, Preetam Gandhi, Katalin Csermak Renner, and Esther K. Schmitt. 2022. “High Prevalence of *Plasmodium Falciparum* K13 Mutations in Rwanda Is Associated With Slow Parasite Clearance After Treatment With Artemether-Lumefantrine.” The Journal of Infectious Diseases 225 (8): 1411–14.

Straimer, Judith, Nina F. Gnädig, Barbara H. Stokes, Michelle Ehrenberger, Audrey A. Crane, and David A. Fidock. 2017. “*Plasmodium Falciparum* K13 Mutations Differentially Impact Ozonide Susceptibility and Parasite Fitness *in Vitro*.” mBio 8 (2). 10.1128/mbio.00172-17.

Straimer, Judith, Nina F. Gnädig, Benoit Witkowski, Chanaki Amaratunga, Valentine Duru, Arba Pramundita Ramadani, Mélanie Dacheux, et al. 2015. “K13-Propeller Mutations Confer Artemisinin Resistance in Plasmodium Falciparum Clinical Isolates.” Science 347 (6220). 10.1126/science.1260867.

Sutherland, Colin J., Ryan C. Henrici, and Katerina Artavanis-Tsakonas. 2021. “Artemisinin Susceptibility in the Malaria Parasite Plasmodium Falciparum: Propellers, Adaptor Proteins and the Need for Cellular Healing.” FEMS Microbiology Reviews 45 (3). 10.1093/femsre/fuaa056.

Takala-Harrison, Shannon, Taane G. Clark, Christopher G. Jacob, Michael P. Cummings, Olivo Miotto, Arjen M. Dondorp, Mark M. Fukuda, et al. 2013. “Genetic Loci Associated with Delayed Clearance of *Plasmodium Falciparum* Following Artemisinin Treatment in Southeast Asia.” Proceedings of the National Academy of Sciences 110 (1): 240–45.

Takala-Harrison, Shannon, Christopher G. Jacob, Cesar Arze, Michael P. Cummings, Joana C. Silva, Arjen M. Dondorp, Mark M. Fukuda, et al. 2015. “Independent Emergence of Artemisinin Resistance Mutations among Plasmodium Falciparum in Southeast Asia.” The Journal of Infectious Diseases 211 (5): 670–79.

Tirrell, Abigail R., Katelyn M. Vendrely, Lisa A. Checkley, Sage Z. Davis, Marina McDew-White, Ian H. Cheeseman, Ashley M. Vaughan, François H. Nosten, Timothy J. C. Anderson, and Michael T. Ferdig. 2019. “Pairwise Growth Competitions Identify Relative Fitness Relationships among Artemisinin Resistant Plasmodium Falciparum Field Isolates.” Malaria Journal 18 (1): 295.

Trager, W., and J. B. Jensen. 1976. “Human Malaria Parasites in Continuous Culture.” *Science (New York*, N.Y.) 193 (4254): 673–75.

Tumwebaze, Patrick K., Melissa D. Conrad, Martin Okitwi, Stephen Orena, Oswald Byaruhanga, Thomas Katairo, Jennifer Legac, et al. 2022. “Decreased Susceptibility of Plasmodium Falciparum to Both Dihydroartemisinin and Lumefantrine in Northern Uganda.” Nature Communications 13 (1): 6353.

Tun, Kyaw M., Mallika Imwong, Khin M. Lwin, Aye A. Win, Tin M. Hlaing, Thaung Hlaing, Khin Lin, et al. 2015. “Spread of Artemisinin-Resistant Plasmodium Falciparum in Myanmar: A Cross-Sectional Survey of the K13 Molecular Marker.” The Lancet Infectious Diseases 15 (4). 10.1016/S1473-3099(15)70032-0.

Tun, Kyaw Myo, Atthanee Jeeyapant, Mallika Imwong, Min Thein, Sai Soe Moe Aung, Tin Maung Hlaing, Prayoon Yuentrakul, et al. 2016. “Parasite Clearance Rates in Upper Myanmar Indicate a Distinctive Artemisinin Resistance Phenotype: A Therapeutic Efficacy Study.” Malaria Journal 15 (1): 185.

Uwimana, Aline, Eric Legrand, Barbara H. Stokes, Jean Louis Mangala Ndikumana, Marian Warsame, Noella Umulisa, Daniel Ngamije, et al. 2020. “Emergence and Clonal Expansion of in Vitro Artemisinin-Resistant Plasmodium Falciparum kelch13 R561H Mutant Parasites in Rwanda.” Nature Medicine 26 (10). 10.1038/s41591-020-1005-2.

Uwimana, Aline, Noella Umulisa, Meera Venkatesan, Samaly S. Svigel, Zhiyong Zhou, Tharcisse Munyaneza, Rafiki M. Habimana, et al. 2021. “Association of Plasmodium Falciparum kelch13 R561H Genotypes with Delayed Parasite Clearance in Rwanda: An Open-Label, Single-Arm, Multicentre, Therapeutic Efficacy Study.” The Lancet Infectious Diseases 21 (8): 1120–28.

Wangdi, Kinley, Ayodhia Pitaloka Pasaribu, and Archie C. A. Clements. 2021. “Addressing Hard-to-reach Populations for Achieving Malaria Elimination in the Asia Pacific Malaria Elimination Network Countries.” Asia & the Pacific Policy Studies 8 (2): 176–88.

Wang, Xiaoxiao, Wei Ruan, Shuisen Zhou, Fang Huang, Qiaoyi Lu, Xinyu Feng, and He Yan. 2020. “Molecular Surveillance of Pfcrt and k13 Propeller Polymorphisms of Imported Plasmodium Falciparum Cases to Zhejiang Province, China between 2016 and 2018.” Malaria Journal 19 (1): 59.

Wang, Zenglei, Sony Shrestha, Xiaolian Li, Jun Miao, Lili Yuan, Mynthia Cabrera, Caitlin Grube, Zhaoqing Yang, and Liwang Cui. 2015. “Prevalence of K13-Propeller Polymorphisms in Plasmodium Falciparum from China-Myanmar Border in 2007-2012.” Malaria Journal 14(1). 10.1186/s12936-015-0672-9.

White, Nicholas J. 2004. “Antimalarial Drug Resistance.” The Journal of Clinical Investigation 113 (8): 1084–92.

“WHO Guidelines for Malaria.” 2017. World Health Organization. July 14, 2017. https://www.who.int/publications/i/item/guidelines-for-malaria.

Windle, Sean T., Kristin D. Lane, Nahla B. Gadalla, Anna Liu, Jianbing Mu, Ramoncito L. Caleon, Rifat S. Rahman, Juliana M. Sá, and Thomas E. Wellems. 2020. “Evidence for Linkage of pfmdr1, Pfcrt, and pfk13 Polymorphisms to Lumefantrine and Mefloquine Susceptibilities in a Plasmodium Falciparum Cross.” International Journal for Parasitology, Drugs and Drug Resistance 14 (December):208–17.

Witkowski, Benoit, Chanaki Amaratunga, Nimol Khim, Sokunthea Sreng, Pheaktra Chim, Saorin Kim, Pharath Lim, et al. 2013. “Novel Phenotypic Assays for the Detection of Artemisinin-Resistant Plasmodium Falciparum Malaria in Cambodia: In-Vitro and Ex-Vivo Drug-Response Studies.” The Lancet Infectious Diseases 13 (12). 10.1016/S1473-3099(13)70252-4.

Witkowski, Benoit, Nimol Khim, Pheaktra Chim, Saorin Kim, Sopheakvatey Ke, Nimol Kloeung, Sophy Chy, et al. 2013. “Reduced Artemisinin Susceptibility of Plasmodium Falciparum Ring Stages in Western Cambodia.” Antimicrobial Agents and Chemotherapy 57 (2): 914– 23.

World Health Organization. 2023. *World Malaria Report* 2023. World Health Organization. “World Malaria Report 2024.” n.d. Accessed March 7, 2025. https://www.who.int/teams/global-malaria-programme/reports/world-malaria-report-2024.

Ye, Run, Dongwei Hu, Yilong Zhang, Yufu Huang, Xiaodong Sun, Jian Wang, Xuedi Chen, et al. 2016. “Distinctive Origin of Artemisinin-Resistant Plasmodium Falciparum on the China-Myanmar Border.” Scientific Reports 6 (1): 20100.

Zhang, Jie, Na Li, Faiza A. Siddiqui, Shiling Xu, Jinting Geng, Jiaqi Zhang, Xi He, et al. 2019. “In Vitro Susceptibility of Plasmodium Falciparum Isolates from the China-Myanmar Border Area to Artemisinins and Correlation with K13 Mutations.” *International Journal for Parasitology*, Drugs and Drug Resistance 10 (August):20–27.

